# A landing pad system for multicopy gene integration in *Issatchenkia orientalis*

**DOI:** 10.1101/2023.05.21.541627

**Authors:** Zia Fatma, Shih-I Tan, Aashutosh Girish Boob, Huimin Zhao

## Abstract

The robust nature of the non-conventional yeast *Issatchenkia orientalis* allows it to grow under highly acidic conditions and therefore, has gained increasing interest in producing organic acids using a variety of carbon sources. Recently, the development of a genetic toolbox for *I. orientalis*, including an episomal plasmid, characterization of multiple promoters and terminators, and CRISPR-Cas9 tools, has eased the metabolic engineering efforts in *I. orientalis*. However, multiplex engineering is still hampered by the lack of efficient multicopy integration tools. To facilitate the construction of large, complex metabolic pathways by multiplex CRISPR-Cas9-mediated genome editing, we developed a bioinformatics pipeline to identify and prioritize genome-wide intergenic loci and characterized 47 sites. These loci are screened for guide RNA cutting efficiency, integration efficiency of a gene cassette, the resulting cellular fitness, and GFP expression level. We further developed a landing pad system using components from these well-characterized loci, which can aid in the integration of multiple genes using single guide RNA and multiple repair templates of the user’s choice. We have demonstrated the use of the landing pad for simultaneous integrations of 2, 3, 4, or 5 genes to the target loci with efficiencies greater than 80%. As a proof of concept, we showed how the production of 5-aminolevulinic acid can be improved by integrating five copies of genes at multiple sites in one step. We have further demonstrated the efficiency of this tool by constructing a metabolic pathway for succinic acid production by integrating five gene expression cassettes using a single guide RNA along with five different repair templates, leading to the production of 9 g/L of succinic acid in batch fermentations. This study demonstrates the effectiveness of a single gRNA-mediated CRISPR platform to build complex metabolic pathways in non-conventional yeast. This landing pad system will be a valuable tool for the metabolic engineering of *I. orientalis*.

**HIGHLIGHTS:**

- *In silico* screening was performed to identify 204 unique guide RNAs in the intergenic regions of the genome.
- 27 loci demonstrated high integration efficiency (>80%) and can be used for efficient gene or long pathway (∼18 kb) integration.
- An array of landing pad systems was installed at four loci for multiplex engineering.
- Multicopy integration of the gene cassettes (GFP, ALAS) resulted in a proportional increase in GFP fluorescence and 5-ALA production.
- A five-gene biosynthetic pathway was integrated into the chromosome in one step.
- This is the first study reporting the development of the landing pad system in *Issatchenkia orientalis*.

## 1. Introduction

Nature has evolved various microbial chassis by adapting them to certain adverse conditions that make them highly convenient hosts for the production of specific chemicals (Bader et al., 2020; Ekas et al., 2019; Fatma et al., 2020; Hug et al., 2020; Llamas et al., 2020; Madzak, 2021; Matsushika et al., 2016; Nielsen et al., 2013). These hosts can be further domesticized by metabolic engineering efforts (Choi et al., 2019; Lee et al., 2011; Volk et al., 2022), which requires the development of genetic tools to engineer platform organisms to produce a wide variety of value-added compounds from renewable raw materials. Among these microbial chassis, a few are already well-studied and come under the category of model organisms, i.e., *Escherichia coli* for bacterial system (Adamczyk and Reed, 2017; Jeschek et al., 2017) and *Saccharomyces cerevisiae* for yeast (Nielsen, 2019), while others that are not well-characterized fall under the list of non-model or non-conventional organisms (Russell et al., 2017).

In recent years, many non-conventional yeasts have emerged as attractive production hosts due to their highly unusual characteristics. Among them, *Issatchenkia orientalis* with high tolerance to multiple stresses, especially low pH, has demonstrated its potential as a robust organism for producing organic acids from various carbon sources (Cao et al., 2020; Lee et al., 2022; Matsushika et al., 2016; Sun et al., 2020; Suthers and Maranas, 2022; Tran et al., 2019; Xiao et al., 2014). So far, *I. orientalis* has been explored to produce ethanol (Isono et al., 2012), succinic acid (Xiao et al., 2014), D-lactic acid (Park et al., 2018), and recently 199.4 g/L of L-malic acid (Xi et al., 2023). A wide variety of genetic tools has been developed to engineer *I. orientalis*, such as a replicative and stable plasmid, promoters and terminators, and CRISPR-Cas9-mediated tools for gene disruption and integration (Cao et al., 2020; Tran et al., 2019). Apart from these genetic tools, the genome-scale metabolic model (Suthers et al., 2020; Suthers and Maranas, 2022) has provided insights to better understand the genotype-to-phenotype relationship and guide the metabolic engineering efforts in *I. orientalis*.

However, in the current scenario, editing different loci simultaneously for multiple deletions or insertions remains inefficient, and performing sequential gene editing is laborious. Also, with current genetic tools, performing the systematic screening of enzyme orthologs in conjunction with DNA copy number tuning is time-consuming. One approach to overcome this drawback is to create a landing pad system that will help efficiently integrate multiple cassettes at different loci effortlessly. A landing pad comprises a DNA fragment (mostly synthetic), a designed sequence placed in the genome of interest in single or multiple copies that are used to achieve a genomic integration of one or multiple genes in a precise and stable manner. The landing pad system has been established in a mammalian cell line, where the authors installed a landing pad at three loci and achieved the insertion of heterologous DNA by using a BxB1 recombinase to integrate nine copies of a monoclonal antibody (Gaidukov et al., 2018). Similarly, a series of synthetic DNA sequences were introduced into the genome of *S. cerevisiae* to facilitate multiple copy gene integration (Baek et al., 2021; Bourgeois et al., 2018). So far, multiplexing for constructing the biosynthetic pathways in *I. orientalis* has not been demonstrated, especially when targeting multiple sites in the genome. During genetic tool development, it has been shown that *I. orientalis* prefers homology-directed repair (HDR) over the non-homologous end-joining (NHEJ) mechanism to repair chromosomal double-strand breaks (DSB) (Tran et al., 2019; Cao et al., 2020). This offers a precise and efficient gene insertion to a specific locus and allows us to further extend our effort of genetic tool development towards controlling gene copy number and expediting strain and pathway engineering.

In this work, we first developed an *in-silico* screening approach to characterize a genome-wide sgRNA library to serve as integration sites for the development of a Cas9-facilitated landing pad system. By applying our in-house developed bioinformatics pipeline, we selected 47 gRNAs that were either located next to a potential essential gene or had a high gene density (>8) for experimental validation. We characterized these loci based on guide RNA cutting efficiency, integration of fluorescent proteins, and GFP expression level. We further developed a landing pad system using specific guide RNAs and intergenic regions and demonstrated simultaneous integrations of 2, 3, 4, or 5 genes to the targeted loci with more than 80% efficiency. This platform also resulted in a highly efficient metabolic pathway assembly for succinic acid production by integrating five genes using single guide RNA with five different repair templates, leading to 9 g/L of succinic acid produced in batch fermentation.

## 2. Materials and Methods

### 2.1 *In silico* screening of genome-wide intergenic sites for constructing a landing pad system

The genome sequence of *I. orientalis* (SD108) comprising 4,925 genes spread across 130 scaffolds, gene table, and necessary annotations were procured from the JGI Genome Portal (https://genome.jgi.doe.gov/portal/Issorie2/Issorie2.download.html). All scripts were developed in-house in Python 3.7.11. The design rules incorporated in these scripts are explained in detail in Section 3.1. The on-target activity prediction tool, CRISPRon (Xiang et al., 2021), was installed based on the guidelines provided in the following GitHub repository (https://github.com/RTH-tools/crispron), while Rule Set 2 scores (Doench et al., 2016) for the guides were obtained by running the supplementary code (https://www.nature.com/articles/nbt.3437#MOESM9) in Python-2.7. The scripts developed in this study to identify the integration sites and perform the analysis of the experimental data are made available at: https://github.com/HuiminZhao/Landing-pad-model.

### 2.2 Strains, plasmids, primers, and media

All the strains and plasmids are listed in **Tables S1** and **S2**, respectively. The guide RNAs and primers used in this study are also listed in **Tables S3** and **S4**, respectively. Assembled plasmids were transformed in the commercial *Escherichia coli* DH5α competent cells (New England Biolabs (NEB), USA). For propagation of the recombinant *E. coli* strains, LB medium (1.5% tryptone, 1.5% NaCl, and 0.5% yeast extract) supplemented with 100 mg/L ampicillin was utilized. For culturing the *I. orientalis* SD108 Δ*ura3* strains, YPAD medium (1% yeast extract, 2% peptone, 0.01% adenine hemisulphate, and 2% glucose) was used. Recombinant *I. orientalis* strains harbouring the plasmid, were grown in Synthetic Complete (SC) dropout medium lacking uracil (SC-URA) with 50 g/L glucose was used. LB broth, bacteriological grade agar, yeast extract, peptone, yeast nitrogen base (w/o amino acid and ammonium sulfate), ammonium sulfate, and D-glucose were obtained from Difco (BD, MD), while complete synthetic medium was purchased from MP Biomedicals (Solon, OH).

### 2.3 Plasmid construction

Oligonucleotides including gBlocks and primers were all synthesized by Integrated DNA Technologies (IDT, Coralville, IA). For construction of the CRISPR plasmid carrying the spacer, the complementary oligos were phosphorylated using T4 polynucleotide kinase from New England Biolabs (NEB, USA) first and then ligated to pVT36b using Golden Gate Assembly (Engler et al., 2009) by *Bsa*I enzyme and T4 DNA ligase (NEB, USA). For integration of the 5-aminolevulinic acid synthase (ALAS) gene, the ALAS gene from *I. orientalis* was amplified by Q5 DNA polymerase (NEB, USA) and cloned to the pZF backbone using HiFi assembly (NEB, USA). The plasmids were verified by digestion by restriction endonucleases (NEB). Plasmid mapping and sequencing alignments were carried out using SnapGene software (GSL Biotech, available at snapgene.com).

### 2.4 Large fragment integration

For obtaining the long fragments of DNA, the PKS4 plasmid (Enghiad et al., 2021) was digested by different sets of restriction enzymes. For the 9 kb fragment, *Nco*I and *Apa*I were used. For the 13 kb fragment, *Nco*I and *Pvu*I were selected. For the 18 kb fragment, *Nco*I and *Fsp*I were utilized. After one hour of digestion, the resulting mixture was directly subjected to column purification to obtain the integration fragments. As for assembly using multiple DNA fragments, Q5 DNA polymerase (NEB, USA) was used for DNA amplification containing an 80 bp overlapped segment with the adjacent fragments. For integration, a modified lithium acetate transformation method was used (Gietz and Schiestl, 2007). Plasmid pVT36b-EG14 was co-transformed with the donor fragments obtained from plasmid PKS4 by restriction digestion or PCR amplification along with 250 bp upstream and downstream homology arms for increasing the integration efficiency. For verification of the integration, colony PCR was performed at the junction of integration using the Phire Hot Start II DNA polymerase (Thermo Fisher Scientific).

### 2.5 Construction and integration of gene of interest into the *I. orientalis* landing pad strain

For the construction of the landing pad strain, the landing pad sequence was synthesized from IDT as a gBlock (**Fig. S1**). In the landing pad sequence, seven different sites (EG9, EG4, EG6, GD5, GD11, GD21, and GD27) were included and each site contained the 50 bp left and right homology arms, 20 bp spacer sequence, and 3 bp PAM sequence. The landing pad sequence was integrated into four different chromosomal sites (i.e., EG14, EG15, EG2, and GD1 sites) in two steps. In the first step, the plasmid pVT36-EG14-EG15 (carrying two guides RNA, cutting at loci EG14 and EG15) was co-transformed with a landing pad sequence flanked with the corresponding homology arm targeting EG14 and EG15 into the SD108 to construct the LPA strain. After confirmation by genotyping and plasmid removal by the 5-Fluoroorotic Acid (5-FOA) method, the LPA strain was further transformed with the plasmid pVT36-EG2-GD1 and the landing pad sequence flanked with the corresponding homology arms targeting EG2 and GD1 to construct the final landing pad strain, LPB strain. For the integration of multiple copies of the gene of interest, 1000 ng of corresponding CRISPR plasmid and 2000 ng of GFP and ALAS cassettes flanked with the homology arm ranging from 38 bp to 42 bp were co-transformed into the LPB strain.

### 2.6 Measurement of cell density and GFP fluorescence

After the integration of GFP into the genome, three different colonies were randomly selected and cultured in SC-URA medium (20 g/L glucose) for three days. The cell culture was diluted 20-fold with phosphate buffered saline (PBS, pH 7.4) and loaded into a 96-well plate. Cell density (OD_600_) was measured using the microplate reader (Tecan Trading AG, Switzerland) and the wtGFP fluorescence was measured using excitation at 488 nm and emission at 509 nm.

### 2.7 Production and quantification of 5-ALA

To produce 5-ALA in *I. orientalis*, the recombinant strains integrated with one to five copies of the ALAS gene and harbouring the CRISPR plasmid were cultured in SC-URA medium (50 g/L glucose) supplemented with 3 g/L glycine, 1 g/L succinate and 50 mM pyridoxal 5′-phosphate (PLP). After culturing for five days, the supernatant was collected and subjected to 5-ALA quantification. For analyzing 5-ALA production, modified Ehrlich’s reagent (0.1 g/mL p-dimethylaminobenzaldehyde (DMAB) and 0.16 mg/mL hypochlorous acid in glacial acetic acid) was used. 500 μL samples or standard solutions were well mixed with 500 μL 1 M acetate buffer (pH 4.6) and 100 mL acetylacetone. Subsequently, the mixture was heated at 100 °C for 10 minutes. After cooling to room temperature, a 200 μL mixture was mixed with 200 μL of freshly prepared modified Ehrlich’s reagent in the dark. After 10 minutes, absorbance at 553 nm was measured with a spectrophotometer (NanoDrop 2000c, Thermo Scientific, USA).

### 2.8 Production and quantification of succinate

For succinic acid production, the recombinant strain integrated with the succinic acid pathway and harbouring the CRISPR plasmid were cultured in SC-URA medium (50 g/L glucose). After two days of culture, the supernatant was collected, and a 10-fold dilution was performed using PBS. The diluted sample was subjected to HPLC analysis to quantify the product. For the HPLC analysis, Agilent 1260 Infinity HPLC equipped with Rezex™ ROA-Organic Acid H^+^ (8%) column (Phenomenex Inc.) and a refractive index detector were used. The column and detector were at 50 °C, and 0.6 mL/min of 0.005 N H_2_SO_4_ was used as the mobile phase.

## 3. Results & Discussion

### 3.1. Development of a bioinformatics algorithm to screen genome-wide intergenic sites for iCas9-mediated gene integration

The availability of specific loci that allows stable and efficient heterologous gene integration and expression is an important prerequisite for constructing a landing pad system. Therefore, we designed an *in-silico* screening platform to simplify the selection and prioritization of genomic loci for the development of a CRISPR/Cas9-facilitated landing pad system in *I. orientalis* (**Fig. 1**). First, we obtained the genome sequence of *I. orientalis* (SD108 strain), the gene table, and necessary annotations from the Joint Genome Institute (JGI) (https://genome.jgi.doe.gov/portal/). In this work, we used iCas9 as the Cas protein (Bao et al., 2015); therefore, we scanned for ‘NGG’ (protospacer adjacent motif (PAM) for iCas9) in both strands of the scaffolds. All 20 bp DNA sequences preceding the PAM were acquired to create a genome-wide gRNA library, and gRNA sequences occurring only once in the genome were selected to ensure single gene integration. To find gRNAs with minimal off-target activity, these were screened against the genome-wide gRNA library for more than X mismatches where X = 5, 6, and 7. While no gRNA sequences with more than 6 or 7 mismatches existed in the genome, we found 2,760 gRNAs with more than 5 mismatches. While the traditional DNA integration approach involves gene replacement, we used intergenic loci for gene integration. Without the availability of knockout screens and genetic interaction datasets, multigene disruption can be particularly harmful (Costanzo et al., 2016; Kuzmin et al., 2018). We utilized the location of the genes in the SD108 gene table to determine that 976 of the 2,760 gRNAs were intergenic. Next, 50 bp upstream and downstream genomic sequences were then obtained based on the location of these gRNAs for homology-directed repair. The guide sequences and homology arms containing polyT, polyG, and *Bsa*I sites were excluded (Lian et al., 2019). To reduce the likelihood of disrupting regulatory elements of the neighbouring genes, gRNAs located in the potential promoter regions (425 bp before the gene) or terminator regions (125 bp after the gene) were also excluded. This resulted in a list of 204 gRNAs to serve as intergenic loci for genome-wide targeted single-gene integration (**Table S5**).

**Fig. 1.**
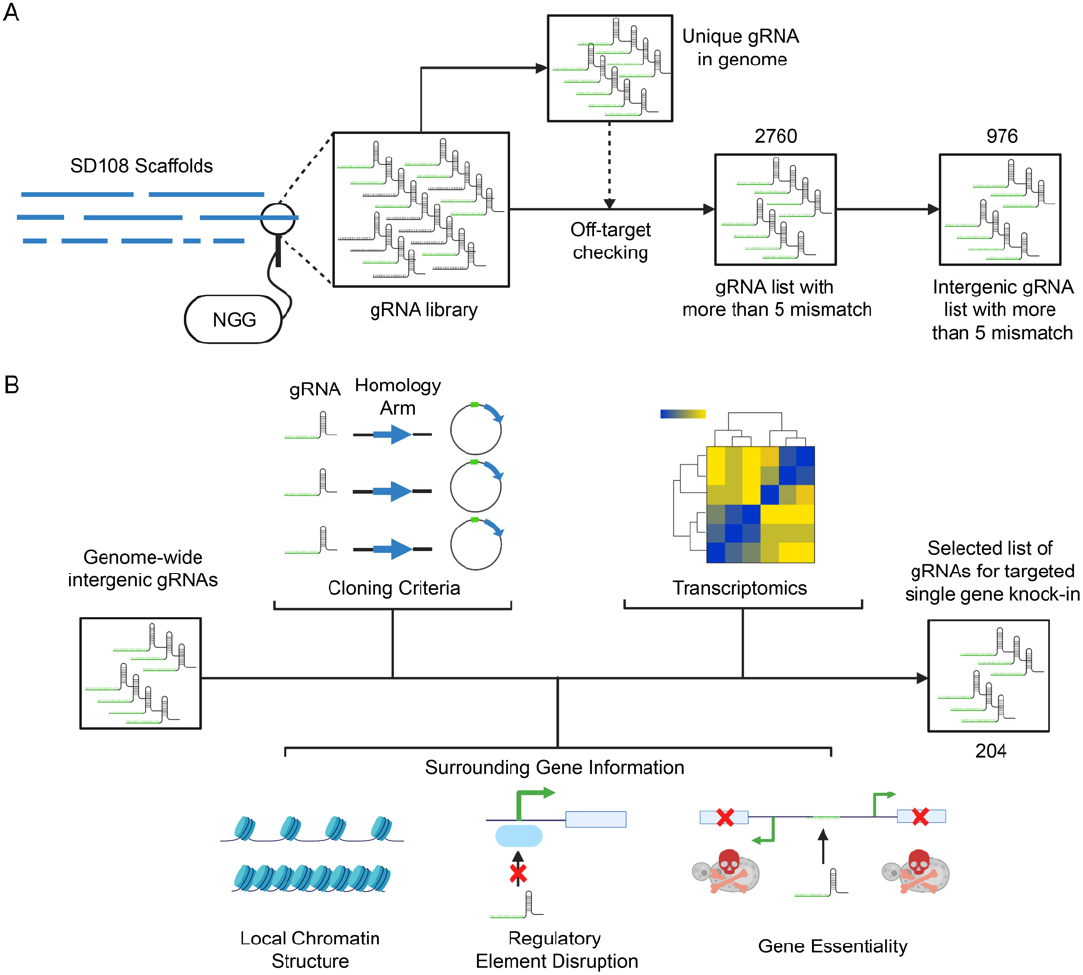
Algorithm for *in silico* screening of genome-wide integration sites in SD108. (A) Selection of gRNAs in intergenic loci for iCas9-mediated integration. The genome is scanned for ‘NGG’ PAM to obtain a guide RNA library. The gRNAs are screened to minimize potential off-targets and filtered based on their genomic location. (B) Incorporation of various factors for prioritization of genomic loci for experimental screening. gRNAs and their corresponding homology arms are refined based on the oligonucleotide synthesis and plasmid cloning criteria. Design rules are implemented to ensure strain stability by avoiding the disruption of regulatory elements and including gene essentiality information, while gene density is added as a proxy for open chromatin. Transcriptomic data is incorporated to select sites close to transcriptionally active genes.

To prioritize the intergenic loci for efficiently constructing the landing pad system, there is a need to incorporate factors that can ensure higher on-target activity. The absence of well-characterized integration sites in non-model organisms and our lack of understanding in selecting which CRISPR/Cas protein to choose can lead to low integration efficiencies. As iCas9 is an improved double mutant of SpCas9 that harbours two mutations, we obtained CRISPROn (Xiang et al., 2021) and Rule Set 2 (Doench et al., 2016) on-target scores to analyze *in vivo* editing efficiency. Another factor that plays an important role in CRISPR-Cas9 genome editing is the local chromatin structure (Verkuijl and Rots, 2019). The integration efficiencies in intergenic loci can be improved by selecting regions with higher chromatin accessibility, i.e., open chromatin structures. Given the unavailability of chromatin data in non-model organisms such as *I. orientalis*, we instead turned to gene density. Gene-rich domains are known to be enriched in open chromatin fibers (Gilbert et al., 2004) and therefore, can serve as a proxy to find intergenic loci in the genome with higher accessibility to iCas9. In this work, we defined gene density as the number of genes located in the 10 kb regions upstream and downstream of the gRNA. We obtained the gene density for each candidate gRNA. We also integrated the available transcriptomic data obtained from a previous study (Cao et al., 2020) to locate transcriptionally active sites for high-level expression of the heterologous genes.

Another aspect of heterologous gene integration for metabolic engineering is stable and reliable expression. Heterologous genes are often equipped with identical promoters and terminators. Over time, the engineered strains can evolve through homologous recombination to lose the integrated genetic information. To overcome this problem, integration sites separated by essential genes are usually utilized (Mikkelsen et al., 2012). To ensure reliable, long-term expression of heterologous enzymes, we identified gRNAs that occur in the intergenic loci adjacent to the putative essential genes of SD108. As *I. orientalis* and *S. cerevisiae* are closely related, we compared all the essential genes of *S. cerevisiae* obtained from the Database of Essential Genes (Luo et al., 2021) to SD108 genes using the BlastP command line (Camacho et al., 2009) with an E-value cutoff of 10^−5^ and identified 933 genes from SD108 as homologs. We selected 34 gRNAs as they are located next to an essential gene (EG) while the remaining 35 gRNAs were selected based on the high gene density (GD) criteria (>8 neighbouring genes) for experimental validation. As the NCBI annotation for SD108 differs from JGI, we further manually discarded 22 gRNAs located within any gene as per NCBI annotation. Overall, we selected 47 gRNAs for characterization and screening. Additional statistics on the integration sites are provided in **Fig. S2**.

### 3.2. Evaluation of guide RNAs for their editing efficiency and intergenic loci for gene expression analysis

To develop a multiplex CRISPR-Cas9 chromosomal integration platform in *I. orientalis*, we sought to design a landing pad system that is often used for precise DNA integration and tuning gene copy numbers in mammalian and other eukaryotic systems (Gaidukov et al., 2018; Baek et al., 2021; Bourgeois et al., 2018). Designing a landing pad system requires a unique gRNA target sequence and flanking regions for recombination that are used for CRISPR-Cas9-mediated gene integration. Using the above-mentioned computational pipeline, we selected 47 distinct guide RNAs present in the intergenic regions of the genome for characterization. These are required for building the landing pad system and additionally could also be used as integration sites for expressing any heterologous genes in *I. orientalis*. It is well known that chromatin structure (heterochromatin or euchromatin) plays a critical role in determining the success of CRISPR-Cas9-mediated genome editing (Chechik et al., 2020; Chen et al., 2022; Daer et al., 2017; Jain et al., 2021). However, we do not have a clear understanding of the chromatin structure of this organism. Therefore, it is possible that some of the selected guide RNAs could be present in a region of poor chromosome accessibility. Therefore, we sought to test the efficacy of each gRNA by introducing it into a plasmid (pVT36b) along with the iCas9 expression cassette and transforming it to *I. orientalis* without providing any repair donor. The efficiency of guide RNA was calculated by comparing the total number of colony-forming units (CFU) resulting from the introduction of a selected gRNA relative to the SD108 strain transformed with a control plasmid (pVT36b, negative control), which does not harbour any guide RNA. As a positive control, we cloned the expression cassette of a previously characterized guide RNA known for its higher editing efficiency (i.e., gRNA residing on JL09_g38).

We found that among 47 gRNAs, 32 gRNAs labeled as EG2, EG3, EG4, EG6, EG7, EG8, EG9, EG10, EG14, EG15, EG16, EG17, GD1, GD5, GD6, GD7, GD9, GD11, GD12, GD14, GD15, GD16, GD17, GD20, GD21, GD22, GD23, GD24, GD26, GD27, GD28, and GD29 could obtain good killing efficiency, which results to lesser or no colonies after transformation, while 15 gRNAs showed poor efficiency (EG1, EG5, EG11, EG12, EG13, EG18, GD2, GD3, GD4, GD8, GD10, GD13, GD18, GD19, GD25) (**Fig. 2A**). The results showed a 68% positive rate of obtaining a good gRNA based on the cell killing efficiency.

**Fig. 2.**
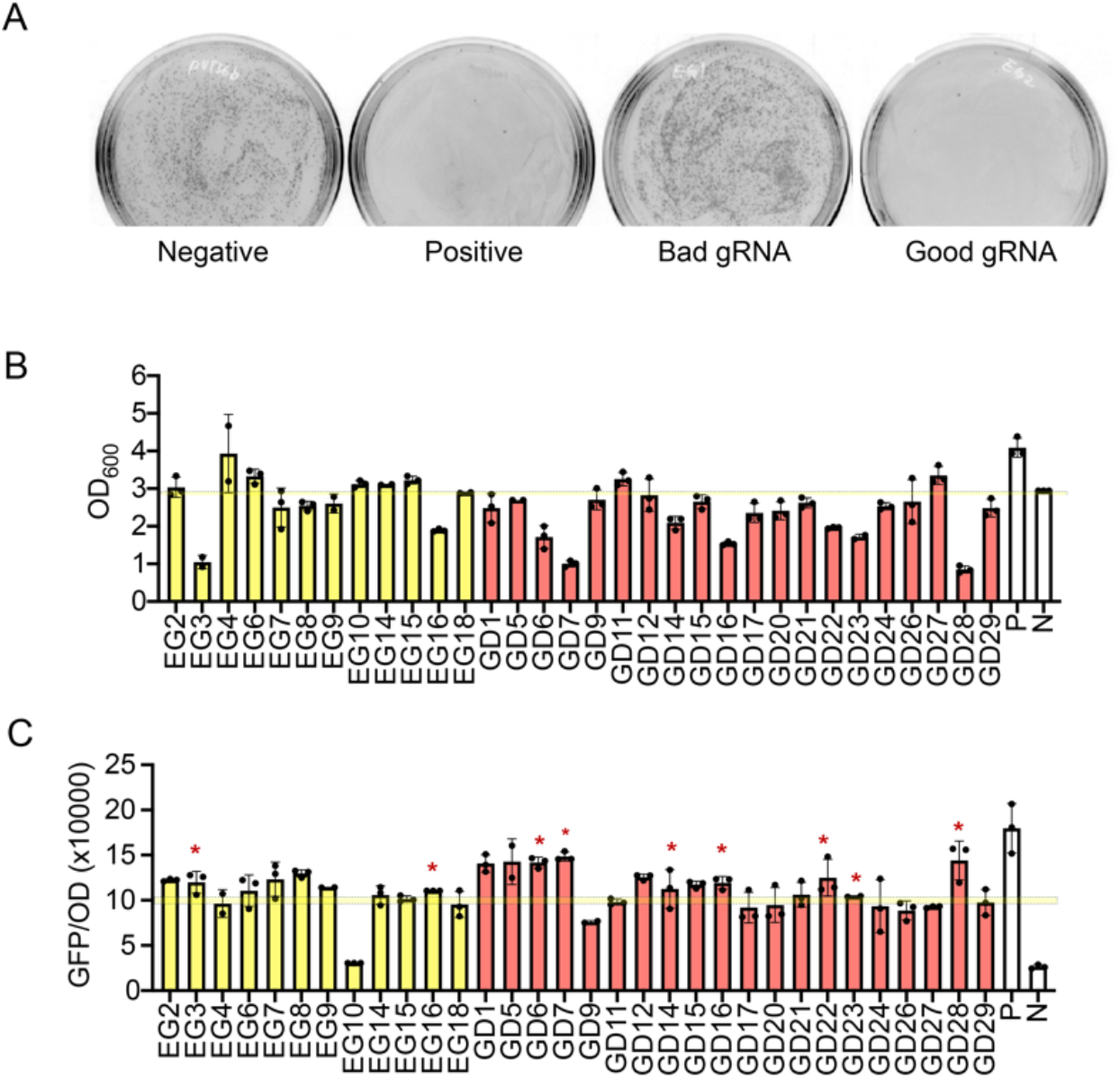
Characterization of gRNAs and integration sites for *in vivo* editing efficiency and GFP expression. (A) gRNA editing efficiency. Negative control (N): pVT36b (backbone). Positive control (P): previously screened gRNA (JL09_g38). Guide RNA with good efficiency: EG2 (shown in the figure), EG3, EG4, EG6, EG7, EG8, EG9, EG10, EG14, EG15, EG16, EG17, GD1, GD5, GD6, GD7, GD9, GD11, GD12, GD14, GD15, GD16, GD17, GD20, GD21, GD22, GD23, GD24, GD26, GD27, GD28, GD29. Guide RNA with poor efficiency: EG1 (shown in the figure), EG5, EG11, EG12, EG13, EG18, GD2, GD3, GD4, GD8, GD10, GD13, GD18, GD19, GD25. (B) Measurement of OD after three days of growth in SC-URA medium, (C) Normalized gene expression after culturing in SC-URA for three days. The bars in colour yellow, red, and white indicate the sites selected by the criteria of having an adjacent essential gene, having high gene density, and controls respectively. The yellow line indicates the cut-off for the selection of good integration sites. Sites with OD higher or similar to the negative control and GFP values higher than or around 10000 a.u. (i.e., half the GFP value obtained from positive control) were selected. Asterisks indicate sites where editing was associated with poor cell growth.

To prioritize gRNA selection in the future, we compared the *in vivo* editing efficiency with the predicted on-target scores obtained from CRISPROn and Rule Set 2 (**Fig. S3**). However, no significant correlation was observed between killing efficiency and the predicted scores or the gene density. These might be due to the different genomic contexts as these models are trained on human cell line datasets (Moreb and Lynch, 2021). Apart from this, as iCas9 harbours two mutations, there might be a different position-dependent nucleotide preference (Kim et al., 2020). Overall, our results indicated that better models are required to predict on-target scores and facilitate higher editing efficiency in *I. orientalis*.

We next assessed the integration efficiency using the gRNAs that exhibited 100% killing efficiency by providing a repair donor. To quantify the integration efficiency of DNA fragments into the intergenic sites, we co-transformed the plasmid carrying the expression cassette for selected gRNA and iCas9 along with a donor DNA comprised of a GFP expression cassette flanked by 50 bp homology arms corresponding to the guide RNA (**Table S3**). The resulting recombinant strains were tested for cellular fitness (by measuring optical density) and GFP signal after three days of growth in the SC-URA medium. The positive control strain (without gRNA) grew to an OD of around 3 within three days in the SC-URA medium. Thus, an OD of 3 was used as the cutoff. Among 32 gRNAs, 23 gRNAs (EG2, EG4, EG6, EG7, EG8, EG9, EG10, EG14, EG15, EG17, GD1, GD5, GD9, GD11, GD12, GD15, GD17, GD20, GD21, GD24, GD26, GD27, GD29) did not show any detrimental effects to the cell growth (**Fig. 2B**). By contrast, 9 gRNAs (EG3, EG16, GD6, GD7, GD14, GD16, GD22, GD23, GD28) showed a poor cell growth. It could be possible that these gRNAs are interfering with some essential or regulatory components of the genome, and the integration of GFP cassettes leads to a loss of function.

We used the plasmid carrying a GFP cassette as a positive control for gene expression analysis. The copy number for the episomal plasmid was previously estimated to be 2 (Cao et al., 2020). Therefore, we expected that the good GFP expression is around or higher than 10000 a.u., which is half the value obtained from the positive control. Higher GFP expression with good cell growth led to the selection of 21 gRNAs which belong to 14 intergenic sites (**Fig. 2C** and **Fig. S4**).

### 3.3. Evaluation of *in vivo* integration efficiency for the long pathway assembly

To implement a multiplex gene integration system in *I. orientalis*, a prerequisite is to test the efficiency of the number of fragments and the size of DNA that can be integrated. Cao *et al*. showed the assembly of a 7 kb plasmid using multiple DNA fragments with an integration efficiency of 100% (Cao et al., 2020). However, multiple fragment integration at one site or multiplex integration of genetic parts in the genome has not been explored so far. Here, we sought to determine the extent of DNA size and the number of fragments that can be integrated into the genome. To demonstrate the integration of large DNA fragments into the chromosomes, we selected site EG14. For donor DNA, we used the holomycin biosynthetic gene cluster from *Bacillus subtilis* (Enghiad et al., 2021) and the 250 bp homology arm surrounding EG14 (**Fig. 3A**). We demonstrated two strategies to integrate the long linear strands of DNA. In the first strategy, the plasmid harbouring the holomycin biosynthetic gene cluster was digested by various restriction enzymes to generate fragments of 9 kb, 13 kb, and 18 kb in size (**Fig. 3B**) and integrated at site EG14. After genotyping five colonies from each transformant, we observed an integration efficiency of 100% (**Fig. 3C**). However, the number of obtained colonies decreased as the size of DNA donor fragments increased (**Fig. 3D**). In the second strategy, the DNA donor was divided into fragments, i.e., four fragments of different sizes to cover 9 kb, five fragments of different sizes to cover 13 kb, and seven fragments of different sizes to cover 18 kb. The fragments were amplified via PCR and then simultaneously transformed to assemble and integrate the donor to the chromosome (**Fig. 3B**). The integration efficiency ranged from 80% to 100% (**Fig. 3E**) and we did not notice any difference in the CFU count in different transformants (**Fig. 3F**). These results showed that the screened site is suitable for integrating long DNA fragments with high efficiency and can be used for developing the landing pad system.

**Fig. 3.**
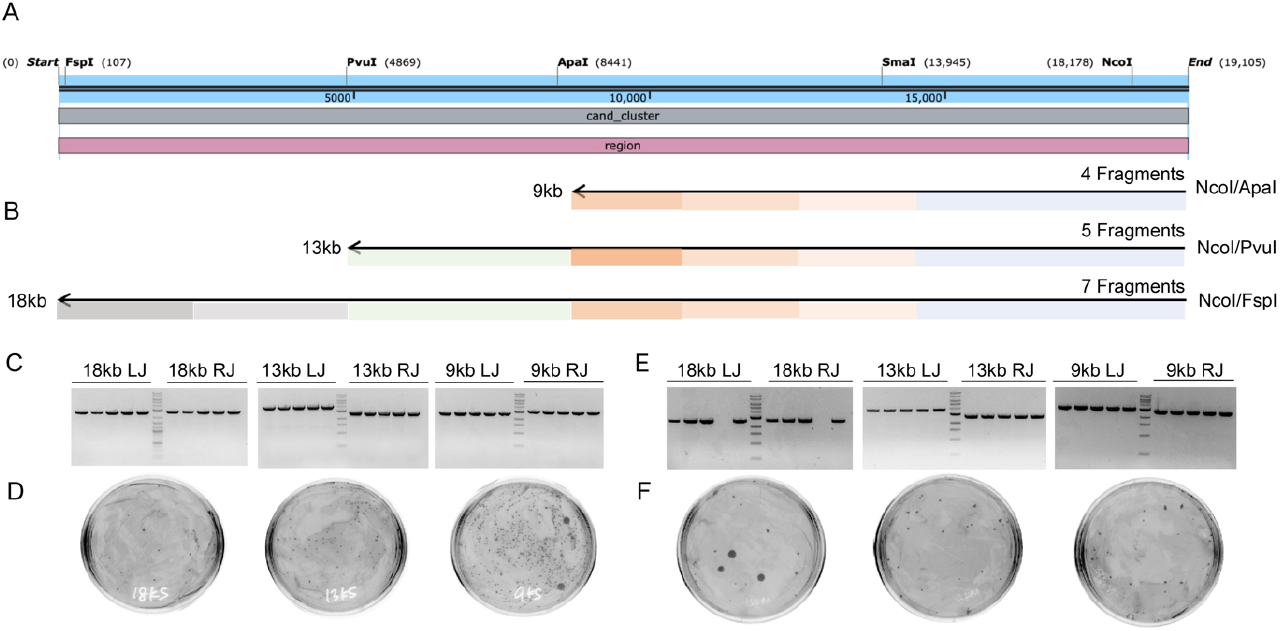
Characterization of site EG14 in terms of donor size and assembly of different numbers of DNA fragments. (A) Linear map of the holomycin biosynthetic gene cluster from *B. subtilis*. (B) A large linear donor is obtained using two methods: restriction digestion or PCR amplification of multiple fragments with overlaps. (C, D) Confirmation of the DNA integration by genotyping and plating using restriction digestion. (E, F) Confirmation of the DNA integration by genotyping and plating using PCR amplification. Successful integration was verified using one primer corresponding to the I. orientalis genome and another corresponding to the PKS4 plasmid. LJ and RJ indicate the left junction and right junction of the integration site.

### 3.4. Design of a synthetic landing pad system for multicopy gene integration in *I. orientalis*

Next, we constructed the landing pad strain using the sites characterized in the previous section. We picked seven sites (EG9, EG4, EG6, GD5, GD11, GD21, and GD27) to be encompassed in the landing pad sequence. These were selected based on *in vivo* editing efficiency, higher cellular fitness, and GFP expression per gram of biomass (**Fig. 4A**). We stitched synthetic components of the seven sites to make an 861 bp fragment. These components contain a 50 bp left homology arm, 20 bp spacer, 3 bp PAM, and 50 bp right homology arm. To construct the landing pad system in *I. orientalis*, we integrated either the complete or partial landing pad sequence to create a different landing pad system into four chromosome loci (i.e., EG14, EG15, EG2, and GD1). To obtain a strain named LPB that can be used to integrate five copies of the gene, the full landing pad sequence was integrated into EG14, and partial landing pad sequences which include guide RNA with homology arm of 5 out of 7 (i.e, EG6, GD5, GD11, GD21, and GD27), 3 out of 7 (i.e, GD11, GD21, and GD27), and 2 out of 7 (i.e, GD21, and GD27) were integrated into EG15, EG2 and GD1 sites, respectively (**Fig. 4A**) and confirmed through genotyping compared to the control strain (**Fig. 4B**). Using this strategy, we have different copies of the loci in the chromosome, thus creating one strain for the integration of one to five copies of genes. For example, as EG9 was only integrated into the EG14 site, there are two EG9 sites in the chromosome and it can be used for two fragments integration (**Fig. 4A**).

**Fig. 4.**
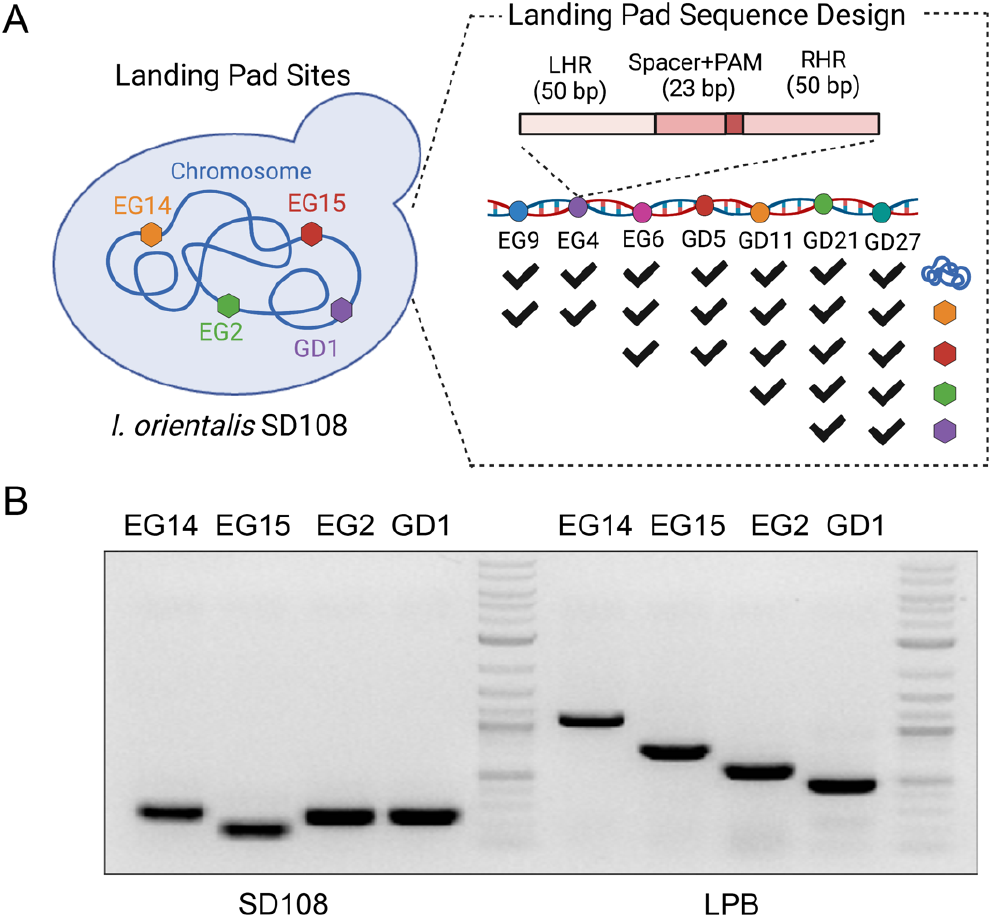
Design and experimental validation of the constructed landing pad strain. (A) The gBlock containing synthetic components of seven sites (20 bp gRNA, 3 bp PAM, and 50 bp for left and right homology arms) were integrated partially or completely in four screened sites (EG14, EG15, EG2, and GD1). Such a strategy created variable repeats of integration sites in the chromosomes. (B) Colony PCR confirmed successful integration of the partial or complete landing pad sequence.

### 3.5. Application of the landing pad strain for multiple gene integration

The primary purpose of constructing a landing pad strain is to enable the chromosomal integration of multiple copies of the same gene expression cassettes or biosynthetic pathways in one step. Following confirmation of the LPB strain by genotyping, we evaluated the targeting efficiency of the introduced landing pad system. Because multiple DSBs could cause genome instability, we started with a proof-of-concept study by integrating varying copy numbers of a GFP cassette. We compared the integration and expression of the GFP cassette in the LPB strain including the control with only one copy of the target site. One to five copies of the GFP expression cassette were integrated into the *I. orientalis* chromosomes by targeting each of the sites in five parallel transformations. As shown in **Fig. 5A**, the GFP signal increased linearly from 41,000 arbitrary units (a.u.) to 200,000 a.u., when the GFP copy number increased from one to five. Besides quantifying GFP expression from multiple copies, integration efficiency is another key factor in evaluating the landing pad system. Thus, three colonies of one to five copies of the GFP strain (i.e., LPBG2, LPBG3, LPBG4, and LPBG5) were randomly selected after transformation and verified by colony PCR. At the specific sites of EG4, GD5, GD11, and GD21, three colonies of each group showed a successful integration band around 2.5 kb (**Fig. 5B**). However, for the other sites (i.e., EG9, EG6, and GD27), the integration efficiency is lower, ranging from 33% to 67% (**Fig. S5**). Overall, this demonstrates that the landing pad system works to efficiently integrate genes with a size of around 3 kb in *I. orientalis*. Moreover, the number of colonies decreases as the number of DSBs in the genome increases (**Fig. S6**).

**Fig. 5.**
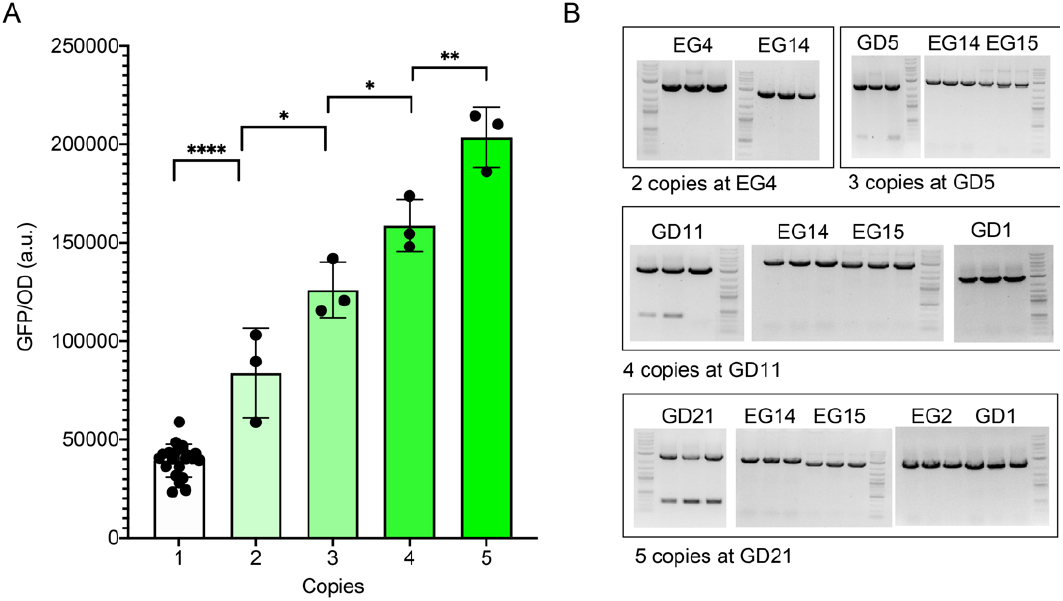
Demonstration of the landing pad strain for integration of multiple copies of the GFP expression cassette. (A) Normalized GFP intensities and (B) Genotyping results for integration of one to five copies at selected sites. *P* value was calculated by two-tailed unpaired *t*-test, \**P* < 0.05, **P < 0.01, \*\*\**P* < 0.001 and \*\*\*\**P* < 0.0001.

To demonstrate the real-world application of the landing pad system, we first focused on the metabolic engineering of *I. orientalis* to produce 5-aminolevulinic acid (5-ALA). 5-ALA is a non-proteinogenic amino acid with many applications in medicine, agriculture, and livestock (Yi et al., 2021). Furthermore, 5-ALA was approved by the U.S. Food and Drug Administration (FDA) as a photodynamic therapy candidate for cancer. The 5-ALA biosynthetic pathway relies on the 5-aminolevulinic acid synthase (ALAS) to convert succinyl-CoA and glycine to 5-ALA (**Fig. 6A**, colored in red). Thus, we hypothesized that more copies of the ALAS gene can lead to higher ALA production. As shown in **Fig. 6B**, the 5-ALA production increased approximately linearly from 25 mg/L to 50 mg/L, 75 mg/L, 120 mg/L, and 150 mg/L as the copy number of the ALAS gene increased from 1 to 2, 3, 4, and 5, respectively. This result indicates there is a clear correlation between the gene copy number and the 5-ALA production titer, which is consistent with a previously reported result (Mao et al., 2020).

**Fig. 6.**
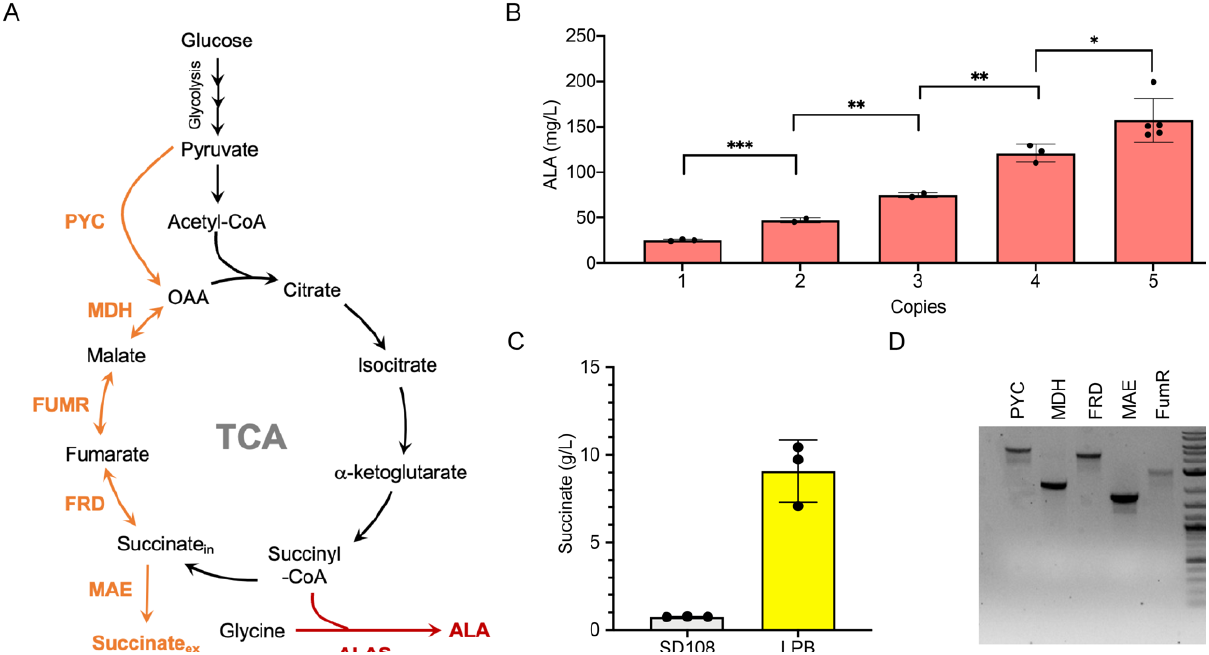
Application of the landing pad strain for metabolic engineering. (A) The metabolic pathway for ALA and succinic acid. (B) Enhanced ALA production using multiple copies of the ALAS gene. (C) Succinic acid production in SD108 and the landing pad strain (i.e., LPB). (D) The genotyping results confirm the integration of PYC, MDH, FRD, MAE, and FUMR. *P* value was calculated by two-tailed unpaired *t*-test, \**P* < 0.05, **P < 0.01, \*\*\**P* < 0.001 and \*\*\*\**P* < 0.0001.

We then focused on the metabolic engineering of *I. orientalis* for the production of succinic acid. Unlike the first application in which a single guide and single donor DNA were used for multiplex integration, this application uses a single guide but different donor DNA for the construction of a complex biosynthetic pathway. Succinic acid is an important four-carbon platform chemical that serves as a precursor for industrially relevant chemicals such as 1,4-butanediol, γ-butyrolactone, tetrahydrofuran, and as a monomer to manufacture various polymers (Kumar et al., 2020). We previously demonstrated that the reductive pathway of the tricarboxylic acid (TCA) cycle in *I. orientalis* could be engineered to produce succinic acid. In this pathway (**Fig. 6A**, colored in orange), pyruvate is first converted to oxaloacetate (OAA) by pyruvate carboxylase (PYC), which is further converted to succinic acid by three enzymes, including malate dehydrogenase (MDH), fumarate reductase (FUMR), and fumarate dehydrogenase (FRD). The fifth enzyme is a dicarboxylate transporter MAE to export the intracellular succinic acid outside the cell. Using this pathway as a proof-of-concept, we demonstrated that our landing pad strain could be used to assemble a five-gene pathway in one round of transformation by using a single gRNA of site GD27. Post transformation, we obtained only three colonies, which indicates that the transformation efficiency was low for the simultaneous integration of five different repair templates. The three colonies were picked and grown in SC-URA with 50 g/L glucose for succinic acid production. As shown in **Fig. S7**, two of the three colonies produced a significant amount of succinic acid and genotyping showed that they successfully integrated all five genes in the succinic acid pathway. The correct strain (i.e., colony 2) was selected and then quantification of the succinic acid production was repeated by batch fermentation. As shown in **Fig. 6C** and **Fig. 6D**, the LPB strain with the reductive TCA pathway for succinic acid production reached 9 g/L succinic acid after two days of culture and the genotyping results showed that all five genes were successfully integrated.

## 4. Conclusion

In this study, we ranked and characterized 47 guide RNAs from 976 candidate intergenic guide RNAs obtained by implementing a bioinformatics pipeline based on various criteria of CRISPR-targeting, gene density, and gene essentiality. *In vivo* screening of these guide RNAs demonstrate that 27 gRNAs are suitable for developing a landing pad system. Moreover, these intergenic sites can also be utilized to express heterologous genes or helper enzymes. We have further shown that we can assemble and integrate up to 18 kb of DNA at these intergenic sites with very high efficiencies. Next, we developed a landing pad system to facilitate multiplex engineering and integrate multiple copies using a single guide and donor DNA (up to five of the same cassette) or a large biosynthetic pathway using a single guide and five different donor DNAs. Implementing this approach, we showed a linear increase in 5-ALA production and 9 g/L of succinic acid production obtained from strains constructed in a single transformation step. We expect that the landing pad system developed in this study will be widely used for efficient multiplex engineering of *I. orientalis* and can be extended to construct longer metabolic pathways in the future.

## Supporting information

Supplementary data to this article can be found online at xxx.

## Author contributions

Z.F., S.T., A.G.B., and H.Z. conceived and designed the study. Z.F., and S.T. performed the experiments. A.G.B. designed the computational pipeline. Z.F., S.T., A.G.B., and H.Z. wrote the manuscript.

## Acknowledgments

This work was supported by U.S. Department of Energy award DE-SC0018420. Any opinions, findings, and conclusions or recommendations expressed in this publication are those of the author(s) and do not necessarily reflect the views of the U.S. Department of Energy. The online tool BioRender (biorender.com) was used to create Fig. 1 and Fig. 4A. The authors acknowledge the use of computing facilities of Biocluster at the Carl R. Woese Institute for Genomic Biology.

