## Supplementary material for "A landing pad system for multicopy gene integration in *Issatchenkia orientalis*": SI

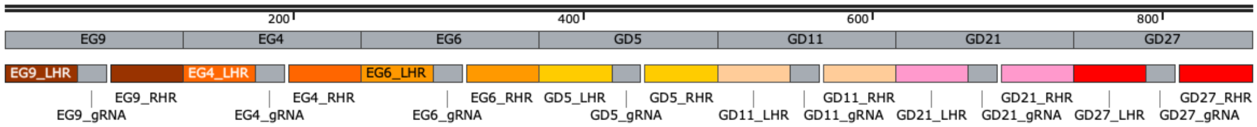


**New landing pad gblock**
861 bp

TTGGAAATTCCATGGCATGTATAAATAACGGATGTGTGAGTTCAAGATGC**TCTTCCGTGATCCAGGTGTT**
TGGGTTTGTTTGTGTCTGTACTTCTATTGTTTACAATAGACATGGTTCACAGAGGGTTTTCGTGAAGTGC
AAGTGGAGTGTGGAAGTTGAGGCGGACACGTGC**GTCGCTGTCACGGGATCACG**TGGTGCGCCACTGTTGG
CATGGCTGCCTGTGGGACTCTTGTTTGTTGCGAGGGCCTCTGTGGTGTCACAGCAAGGCCCGTCTATACC
AACATGTGCAGAGCAC**ACGACAGACGACGCCACGCG**TGGAGGGGTGGCTGCCCCTGTGTGTGGATATAAC
ATACACACACACGGACAGATAAGCGTATAGGTGGAATCCAGTTGATTTCTATCAGGTTAACCACATACA**C**
**TCCGTATGGCTAAGAGTGC**AGGAACCTTTGTGTAACAGTAAGGTAAAACTTTCGACATACAGAGTTTCCG
TGGACGCATCCTTTATCTCGGACTTGACTCGGCACCCTCTTGCCGCGGCACC**TGTAGGTATTTTCCGCGG**
**CA**AGGGCCTCTCGAGCCGCGGAACAACCAATATTCTGTCATTTCCACTCCCGTGAGTCCTGAGATTGAAA
TCTCATAGTTAAGTTGGCTTCCTTTTGCTCTAATT**TCTTTAATACCTGTTGCGTA**TGGTCTGTCAGCAAT
GTTTCTCAATTCTCATTGACAAAGCCAGAAATATGCAACCACGTGTTGCAGCTTACTTACGTATACAGAT
TATCCTCTCTATGGACTC**CCTAGGTACTGGCTGAAGTA**TGGGTACGTGGCACCTTGTGATGCGCCGTAAC
GTCCAGCTACGATGGATGCTG

**Fig. S1.** Design and the underlying DNA sequence of the gBlock used to introduce the landing pad sequence in the *I. orientalis* genome. The color corresponds to the landing pad gBlock map. The brown, dark orange, orange, yellow, peach, pink and red colors correspond to EG9, EG4, EG6, GD5, GD11, GD21 and GD27 sequences, respectively. These sequences include 50 bp right homology arm, 20 bp gRNA targeting sequence (bold), 3 bp PAM sequence (underlined), and 50 bp left homology arm.

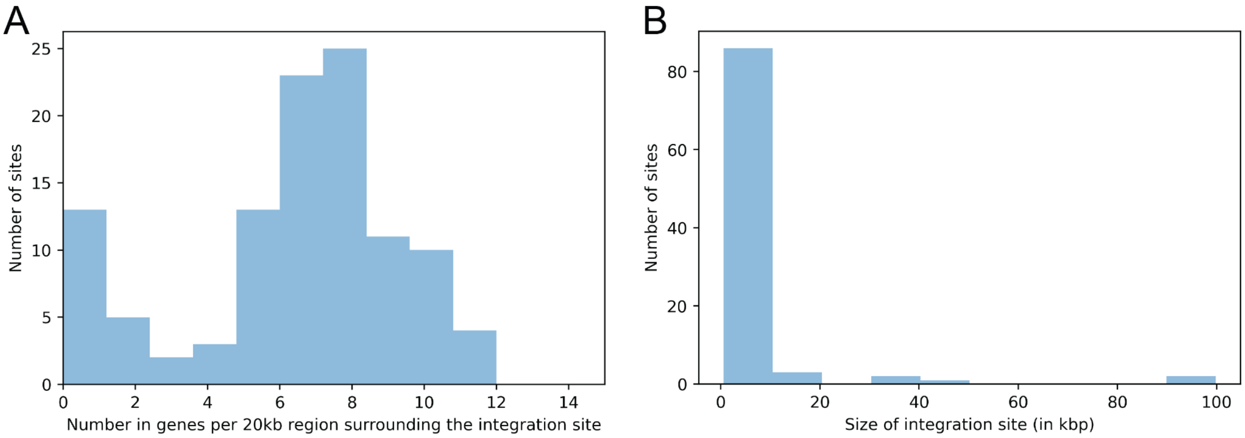


**Fig. S2.** Statistics of the genome-wide sites obtained from the *in silico* platform. (A) Gene density or number of genes in the 20 kb region surrounding the intergenic sites. Majority of sites have more than 4 genes in vicinity. (B) Length distribution of the intergenic sites. Most of the intergenic regions are less than 10 kb in size. Note sites located on scaffolds with no neighboring genes are excluded from the data in the figure.

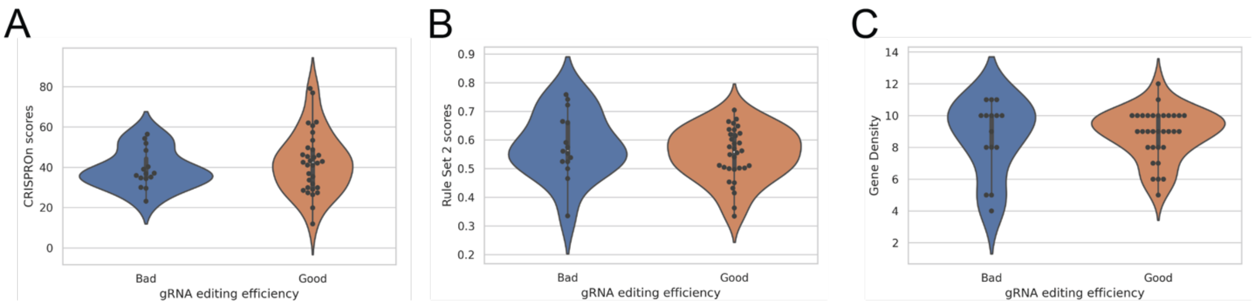


**Fig. S3.** gRNA editing efficiency compared with (A) State-of-the-art machine learning model, CRISPRon, (B) Rule Set 2 scores and (C) Gene Density. A ‘Good’ gRNA is defined as observing no colonies if the plasmid containing sgRNA is transformed into SD108 with no repair cassette. No significant correlation can be observed with any ML model or gene density for gRNA prioritization.

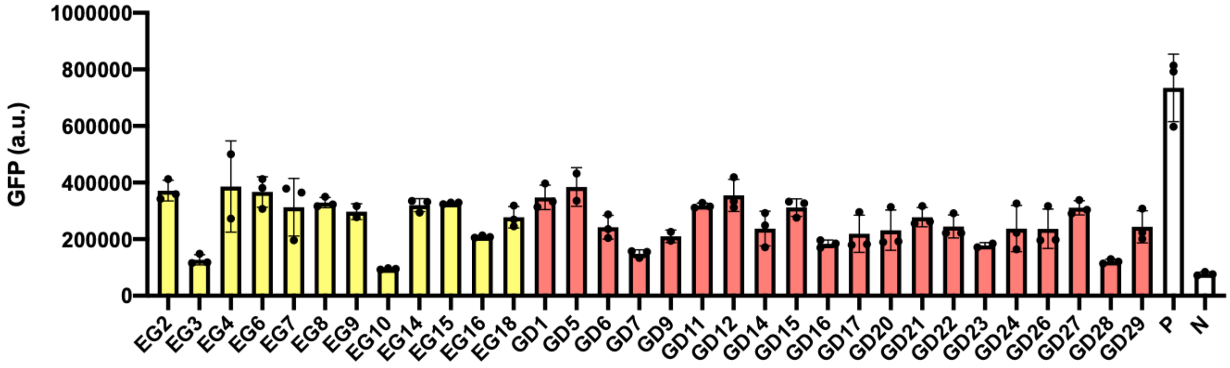


**Fig. S4.** Absolute GFP expression obtained from integrating the GFP cassette into the chromosomal sites. a.u.: arbitrary unit.

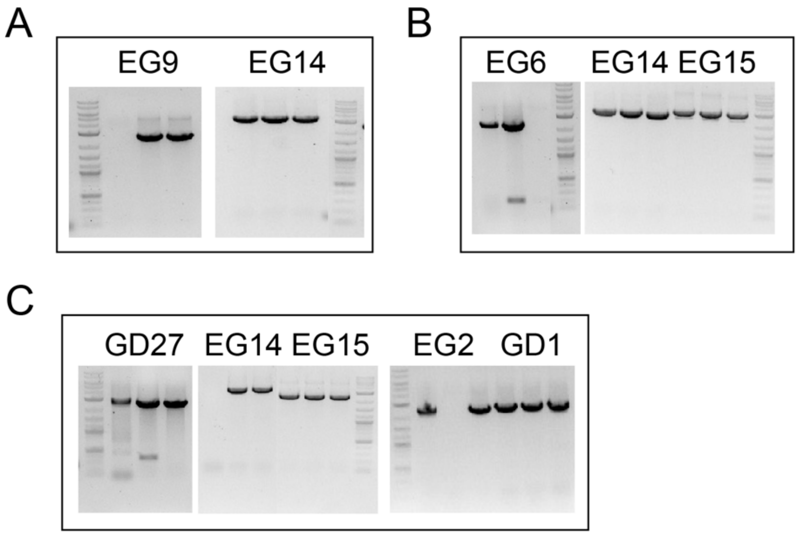


**Fig. S5.** Integration of GFP into the landing pad strain by the other three sites with lower efficiency. (A) two copies at EG9 (B) three copies at EG6 and (C) five copies at GD27.

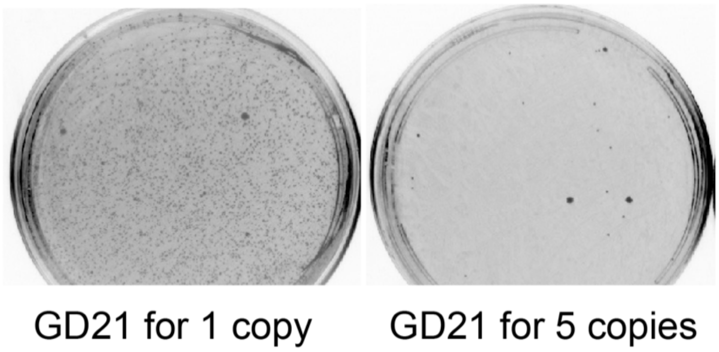


**Fig. S6.** DSB resulted in lower number of colonies when using the same guide RNA corresponding to GD21 in SD108 (i.e., one copy) and LPB (i.e., five copies).

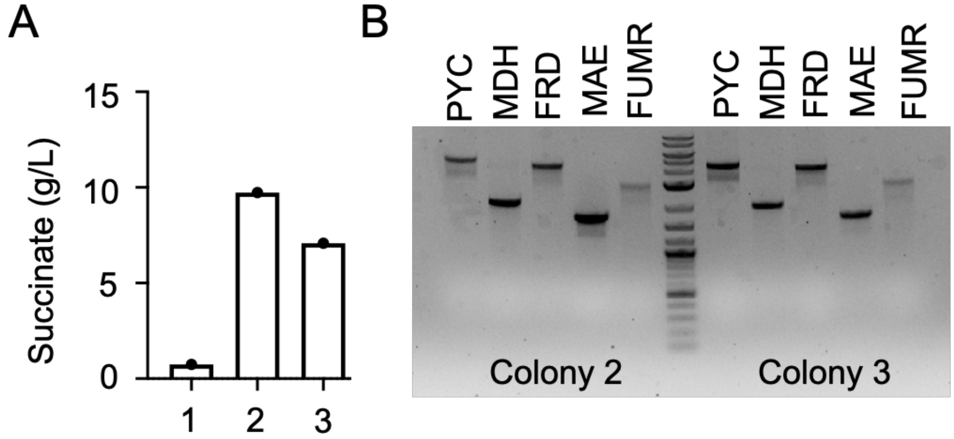


**Fig. S7.** Construction of succinic acid producing strain using the landing pad system. (A) After the transformation of the repair template, three colonies were obtained. All colonies were directly picked to the SC-URA with 50 g/L glucose and cultured for two days. Among the three colonies, colony 2 and colony 3 showed significant succinic acid production with a titer of 9 g/L and 7 g/L, respectively. (B) Genotyping results of the integration in colony 2 and colony 3 shows successful integration of all five genes.

**Table S1.** List of strains used in this study.

| Names | Description | Sources |
|---|---|---|
| *E. coli* DH5a | *F- endA1 glnV44 thi-1 recA1 relA1 gyrA96 deoR nupG Φ80d lacZΔM15 Δ(lacZYA-argF) U169, hsdR17(rK- mK+), λ–* | NEB |
| *I. orientalis* SD108 Δura3 | *ura3Δ*, host for plasmids in this study | (Tran et al., 2019) |
| LPA | *I. orientalis* SD108 Δura3 integrated with landing pad sequence at EG14 and EG15 | This study |
| LPB | LPA integrated with landing pad sequence at EG2 and GD1 | This study |
| LPBG2 | LPB strain integrated with 5025p-GFP at EG4 | This study |
| LPBG3 | LPB strain integrated with 5025p-GFP at GD5 | This study |
| LPBG4 | LPB strain integrated with 5025p-GFP at GD11 | This study |
| LPBG5 | LPB strain integrated with 5025p-GFP at GD21 | This study |

**Table S2.** List of plasmids used in this study.

| Names | Description | Sources |
|---|---|---|
| pZF | AmpR, pUC ori, CEN/ARS from *S. cerevisiae*, *ScURA3*, *IoURA3*, *GPMp*, *ENO2t* | This study |
| pVT36b | AmpR, pUC ori, CEN/ARS from S. cerevisiae, *IoURA3*, *TEF*p, FLAG-SV40-iCas9-SV40, *PGK1*t, *RPR1*, tRNA_*Leu*, gRNA scaffold | (Tran et al., 2019) |
| pVT36z | AmpR, pUC ori, CEN/ARS from S. cerevisiae, *IoURA3,* *TEF*p, FLAG-SV40-iCas9-SV40, *PGK1*t | (Tran et al., 2019) |
| pZF-IoALAS | AmpR, pUC ori, CEN/ARS from S. cerevisiae, *ScURA3*, *IoURA3*, *GPM*p, *IoHEM1*, *ENO2*t | (Tran et al., 2019) |
| pVT36b-LP3 | pVT36b with insertion of protospacer (N20) targeting to LP3 site | This study |
| pVT36b-LP4 | pVT36b with insertion of protospacer (N20) targeting to LP4 site | This study |
| pVT36b-LP5 | pVT36b with insertion of protospacer (N20) targeting to LP5 site | This study |
| pVT36b-LP7 | pVT36b with insertion of protospacer (N20) targeting to LP7 site | This study |
| pVT36b-LP8 | pVT36b with insertion of protospacer (N20) targeting to LP8 site | This study |
| pVT36b-LP9 | pVT36b with insertion of protospacer (N20) targeting to LP9 site | This study |
| pVT36b-LP10 | pVT36b with insertion of protospacer (N20) targeting to LP10 site | This study |
| pVT36b-LP11 | pVT36b with insertion of protospacer (N20) targeting to LP11 site | This study |
| pVT36b-LP15 | pVT36b with insertion of protospacer (N20) targeting to LP15 site | This study |
| pVT36b-LP31 | pVT36b with insertion of protospacer (N20) targeting to LP31 site | This study |
| pVT36b-LP37 | pVT36b with insertion of protospacer (N20) targeting to LP37 site | This study |

| | | |
|---|---|---|
| pVT36b-EG1 | pVT36b with insertion of protospacer (N20) targeting to EG1 site | This study |
| pVT36b-EG2 | pVT36b with insertion of protospacer (N20) targeting to EG2 site | This study |
| pVT36b-EG3 | pVT36b with insertion of protospacer (N20) targeting to EG3 site | This study |
| pVT36b-EG4 | pVT36b with insertion of protospacer (N20) targeting to EG4 site | This study |
| pVT36b-EG5 | pVT36b with insertion of protospacer (N20) targeting to EG5 site | This study |
| pVT36b-EG6 | pVT36b with insertion of protospacer (N20) targeting to EG6 site | This study |
| pVT36b-EG7 | pVT36b with insertion of protospacer (N20) targeting to EG7 site | This study |
| pVT36b-GD1 | pVT36b with insertion of protospacer (N20) targeting to GD1 site | This study |
| pVT36b-GD2 | pVT36b with insertion of protospacer (N20) targeting to GD2 site | This study |
| pVT36b-GD3 | pVT36b with insertion of protospacer (N20) targeting to GD3 site | This study |
| pVT36b-GD4 | pVT36b with insertion of protospacer (N20) targeting to GD4 site | This study |
| pVT36b-GD5 | pVT36b with insertion of protospacer (N20) targeting to GD5 site | This study |
| pVT36b-GD6 | pVT36b with insertion of protospacer (N20) targeting to GD6 site | This study |
| pVT36b-GD7 | pVT36b with insertion of protospacer (N20) targeting to GD7 site | This study |
| pVT36b-GD8 | pVT36b with insertion of protospacer (N20) targeting to GD8 site | This study |
| pVT36b-GD9 | pVT36b with insertion of protospacer (N20) targeting to GD9 site | This study |
| pVT36b-GD10 | pVT36b with insertion of protospacer (N20) targeting to GD10 site | This study |

| | | |
|---|---|---|
| pVT36b-GD11 | pVT36b with insertion of protospacer (N20) targeting to GD11 site | This study |
| pVT36b-GD12 | pVT36b with insertion of protospacer (N20) targeting to GD12 site | This study |
| pVT36b-GD13 | pVT36b with insertion of protospacer (N20) targeting to GD13 site | This study |
| pVT36b-GD14 | pVT36b with insertion of protospacer (N20) targeting to GD14 site | This study |
| pVT36b-GD15 | pVT36b with insertion of protospacer (N20) targeting to GD15 site | This study |
| pVT36b-GD16 | pVT36b with insertion of protospacer (N20) targeting to GD16 site | This study |
| pVT36b-GD17 | pVT36b with insertion of protospacer (N20) targeting to GD17 site | This study |
| pVT36b-GD18 | pVT36b with insertion of protospacer (N20) targeting to GD18 site | This study |
| pVT36b-GD19 | pVT36b with insertion of protospacer (N20) targeting to GD19 site | This study |
| pVT36b-GD20 | pVT36b with insertion of protospacer (N20) targeting to GD20 site | This study |
| pVT36b-GD21 | pVT36b with insertion of protospacer (N20) targeting to GD21 site | This study |
| pVT36b-GD22 | pVT36b with insertion of protospacer (N20) targeting to GD22 site | This study |
| pVT36b-GD23 | pVT36b with insertion of protospacer (N20) targeting to GD23 site | This study |
| pVT36b-GD24 | pVT36b with insertion of protospacer (N20) targeting to GD24 site | This study |
| pVT36b-GD25 | pVT36b with insertion of protospacer (N20) targeting to GD25 site | This study |
| pVT36b-GD26 | pVT36b with insertion of protospacer (N20) targeting to GD26 site | This study |
| pVT36b-GD27 | pVT36b with insertion of protospacer (N20) targeting to GD27 site | This study |

| | | |
|---|---|---|
| pVT36b-GD28 | pVT36b with insertion of protospacer (N20) targeting to GD28 site | This study |
| pVT36b-GD29 | pVT36b with insertion of protospacer (N20) targeting to GD29 site | This study |

**Table S3.** List of selected guide RNAs along with the homology arms.

| Name | gRNA sequence | LHR | RHR |
|---|---|---|---|
| EG1 | ATAAGGCCTGGGTATACATC | TGCCACTCACTAGGTAGACATGCTACTTTTCTCTCTTGGTTTTACCACCT | GTGTATGTAGATTAAATTGTCTACATATAAAATGATATGGAAACCAATTC |
| EG2 | TAAGGCCTGGGTATACATCG | GCCACTCACTAGGTAGACATGCTACTTTTCTCTCTTGGTTTTACCACCTA | TGTATGTAGATTAAATTGTCTACATATAAAATGATATGGAAACCAATTCA |
| EG3 | ATACGCCAAAAGTGGCTAGC | AAGAGGTTCTCAATGTGCCTCAAGATCTCAAGCCATGGTTCAACATCTAT | ATGTTCGCGTTGGGTGTTTCCGGGGCCAACAACTTTTCCGCGTAGTATAT |
| EG4 | GTCGCTGTCACGGGATCACG | GGGTTTTCGTGAAGTGCAAGTGGAGTGTGGAAGTTGAGGCGGACACGTGC | TGCGCCACTGTTGGCATGGCTGCCTGTGGGACTCTTGTTTGTTGCGAGGG |
| EG5 | TGCCCCCTTCCAAATGGGCA | GTCAAAGAATGGGCTTGACAACGTTGATCTTAATTTTCCTACATATAAAA | GACAACAAATACAGTTGTCGCGGCACAACAACTTCTGACCAAATGCGCCA |
| EG6 | ACGACAGACGACGCCACGCG | CCTCTGTGGTGTCACAGCAAGGCCCGTCTATACCAACATGTGCAGAGCAC | AGGGGTGGCTGCCCCTGTGTGTGGATATAACATACACACACACGGACAGA |
| EG7 | ACAGACGACGCCACGCGTGG | CTGTGGTGTCACAGCAAGGCCCGTCTATACCAACATGTGCAGAGCACACG | GGTGGCTGCCCCTGTGTGTGGATATAACATACACACACACGGACAGAGAC |
| GD1 | TCGTCTTATCGCATCTCGTG | TCGGTTAGATTTAAAATTGAAAAGTTGAACGATCAATATTTCTAATAGTA | CTTGAAATAACACGGTTCCGGTTGTTCAATAAATTGAATCAAACGATTTT |
| GD2 | ATCGTCAGCATGCAGGTAAC | GAGTATACATCCGTTGCGTATTATCGTGCATACACAGTTGAGCGGAAATT | GGTGTGATCAAAGTTGGGGAGGAACAACGTGAATTACGGCATCTATGGAT |
| GD3 | AGAGCGTTTCGGGCGATTGT | TATGGATGCGTTTCACATTGGTATTCATTCGGATATGGTGAAGCAAGAAC | CTATCGGTGTAGCGTAATTTGACGGCCAGTCATTGCAAAGTGTGAAGACG |
| GD4 | GTTACACTTCGCTAGACTGT | AATTTGACGGCCAGTCATTGCAAAGTGTGAAGACGGCATCGACTAAACGA | GCTTTGCAATAGCATTTCCCCACACAAATCCAATTGTTTTGTATAGCACG |

| | | | |
|---|---|---|---|
| GD5 | CTCCGTATGGCTAAGAGTGC | TAAGCGTATAGGTGGAATCCAGTTGATTTCTATCAGGTTAACCACATACA | AACCTTTGTGTAACAGTAAGGTAAAACTTTCGACATACAGAGTTTCCGTG |
| GD6 | TAGTCAGTAAATCGACTCAA | TTGTGGGCCAGTTTGGTGTTTACTACTCTTCATTTTAGAATATTCCGATT | TGGGCTATCCAGAACGTGTCATCCTGTTCAGATTCATTCCTTTGATAGAA |
| GD7 | TTCCTGCACTCTTAGCCATA | TTCACGGAAACTCTGTATGTCGAAAGTTTTACCTTACTGTTACACAAAGG | AGTGTATGTGGTTAACCTGATAGAAATCAACTGGATTCCACCTATACGCT |
| GD8 | AATACGGCGGCACGCCTAGA | TACAAGTGTATATGAGGGTTAACAGGCTTGCTGTGCGGTGTGCGGGTGGG | GGCGCAGAGGCTTTGTCAGCCCTCTTTCTGTGCACGTGATACCATGTTTT |
| GD9 | AGTCCGCCCTAGGCCGTGCT | TGTTCCGTCGGAATTGTCCTGTTTCTGCTACCGGTATTCTTTCCTCGGCT | CCGCGTTTAGCACACTTCCCGCGGCCAGCCGGGGCGCGACCTCTTTCCTG |
| GD10 | GACTCGGCACCCTCTTGCCG | ATGCCGCGGCAATGTCTGCAGGACTATGACGCATCCTTTATCTCGGACTT | CACCTGTAGGTATTTTCCGCGGCAAGGGCCTCTCGAGCCGCGGAACAACC |
| GD11 | TGTAGGTATTTTCCGCGGCA | GACGCATCCTTTATCTCGGACTTGACTCGGCACCCTCTTGCCGCGGCACC | GCCTCTCGAGCCGCGGAACAACCAATATTCTGTCATTTCCACTCCCGTGA |
| GD12 | GGCAAGGGCCTCTCGAGCCG | CGGACTTGACTCGGCACCCTCTTGCCGCGGCACCTGTAGGTATTTTCCGC | AACAACCAATATTCTGTCATTTCCACTCCCGTGAACAATAGCTGCAGATT |
| GD13 | CCTCCGGTTTGCTCCAGTTC | TCCAATGCTTACATCCATACATAATAAAATTGTCAGTTATTGGCATAAGG | CTAGGTGATGTGGAATACCCAGTATATCAAGGCTCTAAAAAGATACAATA |
| GD14 | ATCGTGTTTAGGTCTAAAAC | GGAATAATTACCATTAATAAAGCCAAAGCATTATCCGAAAAACAAGCAAA | ACGGCTGGCCAAACAATCCGGAAAAGTGTAGCCAGCATGACAGAACTAGC |
| GD15 | GGACGGCTGGCCAAACAATC | GCCAAAGCATTATCCGAAAAACAAGCAAAATCGTGTTTAGGTCTAAAACC | AAAAGTGTAGCCAGCATGACAGAACTAGCAAGAAGTGCTAGAATAGTTAA |
| GD16 | TGGTTGTTCCGCGGCTCGAG | ACTTATCTAATCTGCAGCTATTGTTCACGGGAGTGGAAATGACAGAATAT | CCCTTGCCGCGGAAAATACCTACAGGTGCCGCGGCAAGAGGGTGCCGAGT |

| | | | |
|---|---|---|---|
| GD17 | ACGCGGCCTAGCACGGCCTA | CTCTTTCAGGAAAGAGGTCGCGCCCCGGCTGGCCGCGGGAAGTGTGCTAA | CGGACTAGCCGAGGAAAGAATACCGGTAGCAGAAACAGGACAATTCCGAC |
| GD18 | CGGCCTAGCACGGCCTAGGG | TTTCAGGAAAGAGGTCGCGCCCCGGCTGGCCGCGGGAAGTGTGCTAAACG | ACTAGCCGAGGAAAGAATACCGGTAGCAGAAACAGGACAATTCCGACGGA |
| GD19 | GGCCTAGGGCGGACTAGCCG | AGGTCGCGCCCCGGCTGGCCGCGGGAAGTGTGCTAAACGCGGCCTAGCAC | AAAGAATACCGGTAGCAGAAACAGGACAATTCCGACGGAACAAAACACGG |
| GD20 | TAAGGTCGCATGCCCGCAAC | CGCAGATAAGAAATGTAAGGTTTTAACCAATCACGTCCCGCTTCTGTAGA | ATTGTATTAAATAAAATAAAATAAAATAAAATAAAATAAAATAAAATAAA |
| GD21 | TCTTTAATACCTGTTGCGTA | GTCCTGAGATTGAAATCTCATAGTTAAGTTGGCTTCCTTTTGCTCTAATT | TCTGTCAGCAATGTTTCTCAATTCTCATTGACAAAGCCAGAAATATGCAA |
| GD22 | CAGCAGCCATATCCGGACCC | ATGCAAGCACACTGTGAATGACTAGGGCCAAGAAAAATGACTGCCTGCAA | GCGGCTGTATGCCATGTCGTCATAGTCGAGTAGGAACTGGAAAGAAGTAA |
| GD23 | CCATGTCGTCATAGTCGAGT | AAATGACTGCCTGCAACAGCAGCCATATCCGGACCCAGGGCGGCTGTATG | AACTGGAAAGAAGTAACTCAATTATCATCACTTCAACAGTATAAACAACT |
| GD24 | CAGGACAGAACAGTATTGCA | TAAAGAAATTAGAGCAAAAGGAAGCCAACTTAACTATGAGATTTCAATCT | AATTTCGGTTTTGCTTTCTTTCCTTGTTTTCCTGGTGAAGTAATGCCTAC |
| GD25 | CACAGCCCGAGCGCGTCGTG | GGGCGCGTGCCTACCCCATTGGGAAAAACATCAACACGCGGCGCCGATCA | CAAATGCATTAACACGTGACTGTGGCGCATTGAAACTCGGTGGGAAAAAC |
| GD26 | ATCCTCTCTATGGACTCCCT | AGCTGTGGGCATTCCACCCACGTGTTGCAGCTTACTTACGTATACAGATT | TACTGGCTGAAGTATGGGTACGTGGCACCTTGTGATGCGCCGTAACGTCC |
| GD27 | CCTAGGTACTGGCTGAAGTA | CCACGTGTTGCAGCTTACTTACGTATACAGATTATCCTCTCTATGGACTC | GTACGTGGCACCTTGTGATGCGCCGTAACGTCCAGCTACGATGGATGCTG |
| GD28 | TACTTCCCAAAGGGGAGGGA | CAAGTGGAGATTTAGAAACAATGGAAAATGTAAATGATGCCGCTCTCGGA | AATATTGTCTAGCTCCCTCTCAGTAAAGGGAAACGTCGTTATCTGCCAAA |

| | | | |
|---|---|---|---|
| GD29 | CTAGAGTGACTCGTGTACCC | AATATTCCCTCAAAGTGTACATCAACAATACGTTCAAATGGACAGCTGTT | ATAAGCAGGTATATCTGACTTGGTCTGGCCGTGTCCTATCAGACGATAGT |
| EG8 | AGGCTCAGCACGGTGTAATT | ACTAATAGTATAACTAAATTAGACTGTATTTCAAATGAGCCGCCTTTAGA | AAATTCCATGGCATGTATAAATAACGGATGTGTGAGTTCAAGATGCTCTT |
| EG9 | TCTTCCGTGATCCAGGTGTT | TTGGAAATTCCATGGCATGTATAAATAACGGATGTGTGAGTTCAAGATGC | GTTTGTTTGTGTCTGTACTTCTATTGTTTACAATAGACATGGTTCACAGA |
| EG10 | CGCGGAAAAGTTGTTGGCCC | TTATTATATGCTGAGGCGGAAAATTCAATCATTTCTTACATAATATACTA | AAACACCCAACGCGAACATCCAGCTAGCCACTTTTGGCGTATATAGATGT |
| EG11 | TTCTAGTGGCACAGTGCCAT | TCACAGGTTTATGCTGTTTGCCCGTCTCTGTAAAATATCTCGGAACAGTG | GCTAATTTTGCTGACAATTTGGCCCGTTCGGCGTGTATGCTTGTCAGTAA |
| EG12 | ACCTGCGCAGCGGAGCGGGC | AGAAGCCCATTTTCGGGCGAGAAGGGGTAACGGACGCCGGGGTTCTCGCC | GGAAAAAGAAACGAGGCGCGGGCGGTCACGTGTGCAGTCAAGTGTTAGAC |
| EG13 | ACTGTGGTCACGTGTGCATG | GTGTGCAGTCAAGTGTTAGACTGTGGTCACGTGTGCAGTCACGTGTTAGA | CGGTCACGTGGTTTTAGTCAGAGACTTTGTTATTTCAAAATACCTCACTG |
| EG14 | GTAGCGTGTAGTGGTTAGAC | TTTCCAAAATGATACGCGGTCGTGCTATTATGGTTTCCAAAAGTTTAATG | TAGGTCCATCTATACATGTAGAAAATAATCATTGTGAATATAAGTTATAA |
| EG15 | TACGTCAGACCGGGCTTACC | CTGGCATTGAAGAAGTACCTAAGACGGAAACCTGAATAATGGAGTGAGTT | ACTATTCCACAAGCCAATAGAAGTCACGAGCATCTCCTTTTGTCTTCCCA |
| EG16 | GTCGTGTGCTCTGCACATGT | TGTTATATCCACACACAGGGGCAGCCACCCCTCCACGCGTGGCGTCGTCT | TATAGACGGGCCTTGCTGTGACACCACAGAGGGGGACATTGACTTTAGTG |
| EG17 | TATAAACGCAGCACCCCGGC | TTGTGCAACTGTGGCCTAAACGTTACTTATGTACTCAGAGATACGTATAC | TTGATGTCCGAAATAGAAGGTTCATACTATAAGTGAGTTATCGGACTTGG |
| EG18 | CTAACCTCAAGTGAGGGGAA | TGTTTGTTCTCGAGTAAATGAGAAACCCAAAAGGTAACAACTTAGAGTAA | GTCAATATAAATAGTGCAAAATAGCTGCAAACCCATGACAGTAAACAGTG |

**Table S4.** List of primers used in the study.

| Names | Sequence (5' to 3') |
|---|---|
| EG1-F | TGCAATAAGGCCTGGGTATACATC |
| EG1-R | AAACGATGTATACCCAGGCCTTAT |
| EG2-F | TGCATAAGGCCTGGGTATACATCG |
| EG2-R | AAACCGATGTATACCCAGGCCTTA |
| EG3-F | TGCAATACGCCAAAAGTGGCTAGC |
| EG3-R | AAACGCTAGCCACTTTTGGCGTAT |
| EG4-F | TGCAGTCGCTGTCACGGGATCACG |
| EG4-R | AAACCGTGATCCCGTGACAGCGAC |
| EG5-F | TGCATGCCCCCTTCCAAATGGGCA |
| EG5-R | AAACTGCCCATTTGGAAGGGGGCA |
| EG6-F | TGCAACGACAGACGACGCCACGCG |
| EG6-R | AAACCGCGTGGCGTCGTCTGTCGT |
| EG7-F | TGCAACAGACGACGCCACGCGTGG |
| EG7-R | AAACCCACGCGTGGCGTCGTCTGT |
| GD1-F | TGCATCGTCTTATCGCATCTCGTG |
| GD1-R | AAACCACGAGATGCGATAAGACGA |
| GD2-F | TGCAATCGTCAGCATGCAGGTAAC |
| GD2-R | AAACGTTACCTGCATGCTGACGAT |
| GD3-F | TGCAAGAGCGTTTCGGGCGATTGT |
| GD3-R | AAACACAATCGCCCGAAACGCTCT |
| GD4-F | TGCAGTTACACTTCGCTAGACTGT |
| GD4-R | AAACACAGTCTAGCGAAGTGTAAC |
| GD5-F | TGCACTCCGTATGGCTAAGAGTGC |

| | |
|---|---|
| GD5-R | AAACGCACTCTTAGCCATACGGAG |
| GD6-F | TGCATAGTCAGTAAATCGACTCAA |
| GD6-R | AAACTTGAGTCGATTTACTGACTA |
| GD7-F | TGCATTCCTGCACTCTTAGCCATA |
| GD7-R | AAACTATGGCTAAGAGTGCAGGAA |
| GD8-F | TGCAAATACGGCGGCACGCCTAGA |
| GD8-R | AAACTCTAGGCGTGCCGCCGTATT |
| GD9-F | TGCAAGTCCGCCCTAGGCCGTGCT |
| GD9-R | AAACAGCACGGCCTAGGGCGGACT |
| GD10-F | TGCAGACTCGGCACCCTCTTGCCG |
| GD10-R | AAACCGGCAAGAGGGTGCCGAGTC |
| GD11-F | TGCATGTAGGTATTTTCCGCGGCA |
| GD11-R | AAACTGCCGCGGAAAATACCTACA |
| GD12-F | TGCAGGCAAGGGCCTCTCGAGCCG |
| GD12-R | AAACCGGCTCGAGAGGCCCTTGCC |
| GD13-F | TGCACCTCCGGTTTGCTCCAGTTC |
| GD13-R | AAACGAACTGGAGCAAACCGGAGG |
| GD14-F | TGCAATCGTGTTTAGGTCTAAAAC |
| GD14-R | AAACGTTTTAGACCTAAACACGAT |
| GD15-F | TGCAGGACGGCTGGCCAAACAATC |
| GD15-R | AAACGATTGTTTGGCCAGCCGTCC |
| GD16-F | TGCATGGTTGTTCCGCGGCTCGAG |
| GD16-R | AAACCTCGAGCCGCGGAACAACCA |
| GD17-F | TGCAACGCGGCCTAGCACGGCCTA |
| GD17-R | AAACTAGGCCGTGCTAGGCCGCGT |
| GD18-F | TGCACGGCCTAGCACGGCCTAGGG |
| GD18-R | AAACCCCTAGGCCGTGCTAGGCCG |

| | |
|---|---|
| GD19-F | TGCAGGCCTAGGGCGGACTAGCCG |
| GD19-R | AAACCGGCTAGTCCGCCCTAGGCC |
| GD20-F | TGCATAAGGTCGCATGCCCGCAAC |
| GD20-R | AAACGTTGCGGGCATGCGACCTTA |
| GD21-F | TGCATCTTTAATACCTGTTGCGTA |
| GD21-R | AAACTACGCAACAGGTATTAAAGA |
| GD22-F | TGCACAGCAGCCATATCCGGACCC |
| GD22-R | AAACGGGTCCGGATATGGCTGCTG |
| GD23-F | TGCACCATGTCGTCATAGTCGAGT |
| GD23-R | AAACACTCGACTATGACGACATGG |
| GD24-F | TGCACAGGACAGAACAGTATTGCA |
| GD24-R | AAACTGCAATACTGTTCTGTCCTG |
| GD25-F | TGCACACAGCCCGAGCGCGTCGTG |
| GD25-R | AAACCACGACGCGCTCGGGCTGTG |
| GD26-F | TGCAATCCTCTCTATGGACTCCCT |
| GD26-R | AAACAGGGAGTCCATAGAGAGGAT |
| GD27-F | TGCACCTAGGTACTGGCTGAAGTA |
| GD27-R | AAACTACTTCAGCCAGTACCTAGG |
| GD28-F | TGCATACTTCCCAAAGGGGAGGGA |
| GD28-R | AAACTCCCTCCCCTTTGGGAAGTA |
| GD29-F | TGCACTAGAGTGACTCGTGTACCC |
| GD29-R | AAACGGGTACACGAGTCACTCTAG |
| EG8-F | TGCAAGGCTCAGCACGGTGTAATT |
| EG8-R | AAACAATTACACCGTGCTGAGCCT |
| EG9-F | TGCATCTTCCGTGATCCAGGTGTT |
| EG9-R | AAACAACACCTGGATCACGGAAGA |
| EG10-F | TGCACGCGGAAAAGTTGTTGGCCC |

| | |
|---|---|
| EG10-R | AAACGGGCCAACAACTTTTCCGCG |
| EG11-F | TGCATTCTAGTGGCACAGTGCCAT |
| EG11-R | AAACATGGCACTGTGCCACTAGAA |
| EG12-F | TGCAACCTGCGCAGCGGAGCGGGC |
| EG12-R | AAACGCCCGCTCCGCTGCGCAGGT |
| EG13-F | TGCAACTGTGGTCACGTGTGCATG |
| EG13-R | AAACCATGCACACGTGACCACAGT |
| EG14-F | TGCAGTAGCGTGTAGTGGTTAGAC |
| EG14-R | AAACGTCTAACCACTACACGCTAC |
| EG15-F | TGCATACGTCAGACCGGGCTTACC |
| EG15-R | AAACGGTAAGCCCGGTCTGACGTA |
| EG16-F | TGCAGTCGTGTGCTCTGCACATGT |
| EG16-R | AAACACATGTGCAGAGCACACGAC |
| EG17-F | TGCATATAAACGCAGCACCCCGGC |
| EG17-R | AAACGCCGGGGTGCTGCGTTTATA |
| EG18-F | TGCACTAACCTCAAGTGAGGGGAA |
| EG18-R | AAACTTCCCCTCACTTGAGGTTAG |
| EG2-LHR-F | TAGGTAGACATGCTACTTTTCTCTCTTGGTTTTACCACCTAacttggctaaag aataaga |
| EG2-RHR-R | TTTCCATATCATTTTATATGTAGACAATTTAATCTACATACAgacccaggtg gctctaga |
| EG3-LHR-F | TCAATGTGCCTCAAGATCTCAAGCCATGGTTCAACATCTATacttggctaaa gaataaga |
| EG3-RHR-R | CGCGGAAAAGTTGTTGGCCCCGGAAACACCCAACGCGAACATgacccag gtggctctaga |
| EG4-LHR-F | TGAAGTGCAAGTGGAGTGTGGAAGTTGAGGCGGACACGTGCacttggctaa agaataaga |
| EG4-RHR-R | ACAAACAAGAGTCCCACAGGCAGCCATGCCAACAGTGGCGCAgacccag gtggctctaga |

| | |
|---|---|
| EG6-LHR-F | TGTCACAGCAAGGCCCGTCTATACCAACATGTGCAGAGCACacttggctaaagaataaga |
| EG6-RHR-R | TGTGTGTGTATGTTATATCCACACACAGGGGCAGCCACCCCTgacccaggtggctctaga |
| EG7-LHR-F | CACAGCAAGGCCCGTCTATACCAACATGTGCAGAGCACACGacttggctaaagaataaga |
| EG7-RHR-R | CCGTGTGTGTGTATGTTATATCCACACACAGGGGCAGCCACCgacccaggtggctctaga |
| GD1-LHR-F | TTTAAAATTGAAAAGTTGAACGATCAATATTTCTAATAGTAacttggctaaagaataaga |
| GD1-RHR-R | TTGATTCAATTTATTGAACAACCGGAACCGTGTTATTTCAAGgacccaggtggctctaga |
| GD5-LHR-F | AGGTGGAATCCAGTTGATTTCTATCAGGTTAACCACATACAacttggctaaagaataaga |
| GD5-RHR-R | CTCTGTATGTCGAAAGTTTTACCTTACTGTTACACAAAGGTTgacccaggtggctctaga |
| GD6-LHR-F | AGTTTGGTGTTTACTACTCTTCATTTTAGAATATTCCGATTacttggctaaagaataaga |
| GD6-RHR-R | AAGGAATGAATCTGAACAGGATGACACGTTCTGGATAGCCCAgacccaggtggctctaga |
| GD7-LHR-F | ACTCTGTATGTCGAAAGTTTTACCTTACTGTTACACAAAGGacttggctaaagaataaga |
| GD7-RHR-R | GGTGGAATCCAGTTGATTTCTATCAGGTTAACCACATACACTgacccaggtggctctaga |
| GD9-LHR-F | GGAATTGTCCTGTTTCTGCTACCGGTATTCTTTCCTCGGCTacttggctaaagaataaga |
| GD9-RHR-R | AGGTCGCGCCCCGGCTGGCCGCGGGAAGTGTGCTAAACGCGGgacccaggtggctctaga |
| GD11-LHR-F | TTTATCTCGGACTTGACTCGGCACCCTCTTGCCGCGGCACCacttggctaaagaataaga |
| GD11-RHR-R | GTGGAAATGACAGAATATTGGTTGTTCCGCGGCTCGAGAGGCgacccaggtggctctaga |
| GD12-LHR-F | CTCGGCACCCTCTTGCCGCGGCACCTGTAGGTATTTTCCGCacttggctaaagaataaga |

| | |
|---|---|
| GD12-RHR-R | GCTATTGTTCACGGGAGTGGAAATGACAGAATATTGGTTGTTgacccaggtggctctaga |
| GD14-LHR-F | ACCATTAATAAAGCCAAAGCATTATCCGAAAAACAAGCAAAacttggctaaagaataaga |
| GD14-RHR-R | TGTCATGCTGGCTACACTTTTCCGGATTGTTTGGCCAGCCGTgacccaggtggctctaga |
| GD15-LHR-F | TTATCCGAAAAACAAGCAAAATCGTGTTTAGGTCTAAAACCacttggctaaagaataaga |
| GD15-RHR-R | TCTAGCACTTCTTGCTAGTTCTGTCATGCTGGCTACACTTTTgacccaggtggctctaga |
| GD16-LHR-F | ATCTGCAGCTATTGTTCACGGGAGTGGAAATGACAGAATATacttggctaaagaataaga |
| GD16-RHR-R | CCCTCTTGCCGCGGCACCTGTAGGTATTTTCCGCGGCAAGGGgacccaggtggctctaga |
| GD17-LHR-F | GAAAGAGGTCGCGCCCCGGCTGGCCGCGGGAAGTGTGCTAAacttggctaaagaataaga |
| GD17-RHR-R | TGTCCTGTTTCTGCTACCGGTATTCTTTCCTCGGCTAGTCCGgacccaggtggctctaga |
| GD20-LHR-F | GAAATGTAAGGTTTTAACCAATCACGTCCCGCTTCTGTAGAacttggctaaagaataaga |
| GD20-RHR-R | ATTTTATTTTATTTTATTTTATTTTATTTTATTTAATACAATgacccaggtggctctaga |
| GD21-LHR-F | TTGAAATCTCATAGTTAAGTTGGCTTCCTTTTGCTCTAATTacttggctaaagaataaga |
| GD21-RHR-R | TTCTGGCTTTGTCAATGAGAATTGAGAAACATTGCTGACAGAgacccaggtggctctaga |
| GD22-LHR-F | CACTGTGAATGACTAGGGCCAAGAAAAATGACTGCCTGCAAacttggctaaagaataaga |
| GD22-RHR-R | TTCCAGTTCCTACTCGACTATGACGACATGGCATACAGCCGCgacccaggtggctctaga |
| GD23-LHR-F | CCTGCAACAGCAGCCATATCCGGACCCAGGGCGGCTGTATGacttggctaaagaataaga |
| GD23-RHR-R | ATACTGTTGAAGTGATGATAATTGAGTTACTTCTTTCCAGTTgacccaggtggctctaga |

| | |
|---|---|
| GD24-LHR-F | TAGAGCAAAAGGAAGCCAACTTAACTATGAGATTTCAATCTacttggctaaagaataaga |
| GD24-RHR-R | TACTTCACCAGGAAAACAAGGAAAGAAAGCAAAACCGAAATTgacccaggtggctctaga |
| GD26-LHR-F | CATTCCACCCACGTGTTGCAGCTTACTTACGTATACAGATTacttggctaaagaataaga |
| GD26-RHR-R | CGGCGCATCACAAGGTGCCACGTACCCATACTTCAGCCAGTAgacccaggtggctctaga |
| GD27-LHR-F | GCAGCTTACTTACGTATACAGATTATCCTCTCTATGGACTCacttggctaaagaataaga |
| GD27-RHR-R | ATCGTAGCTGGACGTTACGGCGCATCACAAGGTGCCACGTACgacccaggtggctctaga |
| GD28-LHR-F | ATTTAGAAACAATGGAAAATGTAAATGATGCCGCTCTCGGAacttggctaaagaataaga |
| GD28-RHR-R | ATAACGACGTTTCCCTTTACTGAGAGGGAGCTAGACAATATTgacccaggtggctctaga |
| GD29-LHR-F | TCAAAGTGTACATCAACAATACGTTCAAATGGACAGCTGTTacttggctaaagaataaga |
| GD29-RHR-R | CTGATAGGACACGGCCAGACCAAGTCAGATATACCTGCTTATgacccaggtggctctaga |
| EG8-LHR-F | ATAACTAAATTAGACTGTATTTCAAATGAGCCGCCTTTAGAacttggctaaagaataaga |
| EG8-RHR-R | CTTGAACTCACACATCCGTTATTTATACATGCCATGGAATTTgacccaggtggctctaga |
| EG9-LHR-F | CCATGGCATGTATAAATAACGGATGTGTGAGTTCAAGATGCacttggctaaagaataaga |
| EG9-RHR-R | CCATGTCTATTGTAAACAATAGAAGTACAGACACAAACAAACgacccaggtggctctaga |
| EG10-LHR-F | GCTGAGGCGGAAAATTCAATCATTTCTTACATAATATACTAacttggctaaagaataaga |
| EG10-RHR-R | ATACGCCAAAAGTGGCTAGCTGGATGTTCGCGTTGGGTGTTTgacccaggtggctctaga |
| EG14-LHR-F | TGATACGCGGTCGTGCTATTATGGTTTCCAAAAGTTTAATGacttggctaaagaataaga |

| | |
|---|---|
| EG14-RHR-R | TATATTCACAATGATTATTTTCTACATGTATAGATGGACCTAgacccaggtg gctctaga |
| EG15-LHR-F | AAGAAGTACCTAAGACGGAAACCTGAATAATGGAGTGAGTTacttggctaa agaataaga |
| EG15-RHR-R | CAAAAGGAGATGCTCGTGACTTCTATTGGCTTGTGGAATAGTgacccaggt ggctctaga |
| EG16-LHR-F | CACACACAGGGGCAGCCACCCCTCCACGCGTGGCGTCGTCTacttggctaaa gaataaga |
| EG16-RHR-R | TCAATGTCCCCCTCTGTGGTGTCACAGCAAGGCCCGTCTATAgacccaggt ggctctaga |
| EG17-LHR-F | TGTGGCCTAAACGTTACTTATGTACTCAGAGATACGTATACacttggctaaa gaataaga |
| EG17-RHR-R | GATAACTCACTTATAGTATGAACCTTCTATTTCGGACATCAAgacccaggt ggctctaga |
| EG14-18kb-Frg1-F | tacgcggtcgtgctattatggtttccaaaagtttaatggcataaggtagttttttctct |
| 18kb-Frg1-R | ttctccttaagactgtaccaa |
| 18kb-Frg2-F | tagaaggaggaatagtttgtg |
| 18kb-Frg2-R | ttgagtaggaaatctatcattt |
| 18kb-Frg3-F | cgttcgtaattaaattttaaactt |
| EG14-13kb-Frg1-F | tacgcggtcgtgctattatggtttccaaaagtttaatgcgttcgtaattaaattttaaac |
| 13kb-Frg1-R | cttacggcaaacgttatcca |
| 13kb-Frg2-F | aggtatcagacgcgataaaa |
| EG14-9kb-Frg1-F | aaatgatacgcggtcgtgctattatggtttccaaaagtttaatgccccaggtatcagacg |
| 9kb-Frg1-R | aagtagggtatattcaacatacctt |
| 9kb-Frg2-F | atacttccattaattcccggt |
| 9kb-Frg2-R | caaacattgggaatagaagtg |
| 9kb-Frg3-F | tctacctccctttctagttg |
| 9kb-Frg3-R | ccacaagcttccttcattac |
| 9kb-Frg4-F | ttagatagaccaaccaccat |

| | |
|---|---|
| LP10-4kb-R | tcacaatgattattttctacatgtatagatggacctaggacttattccagtttcttatgg |
| EG14-common-UF | caacgaggtgttattagatg |
| 18kb-LP10-UR (new) | atctctaaaatgccaattagagaaaaaaactaccttatgccattaaacttttggaaacca |
| 13kb-LP10-UR | tttttattgttcgtttaagtttaaaatttaattacgaacgcattaaacttttggaaacca |
| 9kb-LP10-UR (new) | **aaaatttgaaaaatatttttatcgcgtctgatacctggggcattaaacttttggaaacca** |
| 4kb-LP10-UR | ttgagtctttccaaccagacttagtccttgagataggccccattaaacttttggaaacca |
| EG14-common-LF (new) | gaagagtggaaactaacccataagaaactggaataagtcctaggtccatctatacatgta |
| EG14-common-LR | gtaaaaaatacaaacggtaactg |
| EG2-Check-F | agtgatgccctttgagatg |
| EG2-Check-R | agacaaggcaaggattgtg |
| GD1-Check-F | cataatgcgctagttggtg |
| GD1-Check-R | catatggatatggtgccaaag |
| LP4-Check-F | ggtgagggtttgtaacaaag |
| LP4-Check-R | tccgcctcagcatataataac |
| EG4-Check-F | ccacaattccaaacaaggtg |
| EG4-Check-R | gcttgtatccggagaaactc |
| EG6-Check-F | gttagtggctcactatcgtg |
| EG6-Check-R | gtttggacgtgctagacag |
| GD5-Check-F | ggatcgacttgtcgttatc |
| GD5-Check-R | gcatatcatgcaaacagcg |
| GD11-Check-F | ccattttcggtcgaatgc |
| GD11-Check-R | caatgcaggaggaaaacg |
| GD21-Check-F | ccagcttttgatgtcgttc |

| | |
|---|---|
| GD21-Check-R | aaagagtgcctcacactac |
| GD27-Check-F | ttgcttatacagcagctgtg |
| GD27-Check-R | gccaaactaacactgaggtc |
| EG14-Check-F | tggactctctacgtgaatgt |
| EG14-Check-R | tttccttctttctacccccttt |
| EG15-Check-F | GTTTCCCTGTTTTCCTTAGAT |
| EG15-Check-R | TGTGGCCTCGGAGTCCTT |
| EG14-LP-F | TGATACGCGGTCGTGCTATTATGGTTTCCAAAAGTTTAATGTTGGAAATTCCATGGCATG |
| EG14-LP-R | TATATTCACAATGATTATTTTCTACATGTATAGATGGACCTACAGCATCCATCGTAGCTG |
| EG15-LP-F | TGAAGAAGTACCTAAGACGGAAACCTGAATAATGGAGTGAGTTCCTCTGTGGTGTCACAG |
| EG15-LP-R | CAAAAGGAGATGCTCGTGACTTCTATTGGCTTGTGGAATAGTCAGCATCCATCGTAGCTG |
| EG2-LP-F | TAGGTAGACATGCTACTTTTCTCTCTTGGTTTTACCACCTAGACGCATCCTTTATCTCGG |
| EG2-LP-R | TTTCCATATCATTTTATATGTAGACAATTTAATCTACATACACAGCATCCATCGTAGCTG |
| GD1-LP-F | TTTAAAATTGAAAAGTTGAACGATCAATATTTCTAATAGTACCCGTGAGTCCTGAGATTG |
| GD1-LP-R | TTGATTCAATTTATTGAACAACCGGAACCGTGTTATTTCAAGCAGCATCCATCGTAGCTG |
| GD27-FBA1p-IoPYC-F | CAGCTTACTTACGTATACAGATTATCCTCTCTATGGACTCgatcgattgtatgtgtattg |
| GD27'-TEF1t-IoPYC-R | GCCAAAAAGTTCTAAATCCTCTTTAACTTGTCTTCAACAggctaaagaataagatgaacg |
| GD27-TEF1ap-IoMDH-F | GCAGCTTACTTACGTATACAGATTATCCTCTCTATGGACTCgatcgtttgaaacatcatg |

| | |
|---|---|
| LP10-Trp3t-IoMDH-R | TATATTCACAATGATTATTTTCTACATGTATAGATGGACCTAtgttcaaatgcttgtctg |
| GD27-PGK1p-IoFUMR-F | GCAGCTTACTTACGTATACAGATTATCCTCTCTATGGACTCcgctacttagcttctatag |
| LP11-ENO2t-IoFUMR-R | CAAAAGGAGATGCTCGTGACTTCTATTGGCTTGTGGAATAGTcgtggaacatgtttgtac |
| GD27-TDH3p-FRDg'-F | GCAGCTTACTTACGTATACAGATTATCCTCTCTATGGACTCtcgttgatttaacctgatc |
| EG2-PGK1t-FRDg'-R | TTTCCATATCATTTTATATGTAGACAATTTAATCTACATACAGGTCctttgtgtaagaag |
| GD27-g853-MAE-F | TTGCAGCTTACTTACGTATACAGATTATCCTCTCTATGGACTCcgaaaaatgcaccacac |
| GD1-g3767-MAE-R | TCAATTTATTGAACAACCGGAACCGTGTTATTTCAAGggaaacttaagaattcaattaac |

Table S5

| grna_seq | scaffold | loc | strand | LHR | RHR | grna_with_PAM | gene_density |
|---|---|---|---|---|---|---|---|
| **TTAACACATGCCCTCGGCCG** | scaffold_1 | 585 | + | CTCCGGACTACCGAGACTGCCCGCACCGGCTTTAATCCCAATAAGGAAAG | TCCTTATTTGACTGGGAAGATCCTTATTATTTGATTCTTAGGCTATGCTG | TTAACACATGCCCTCGGCCGAGG | 2 |
| **TAAGGTCGCATGCCCGCAAC** | scaffold_1 | 31358 | + | CGCAGATAAGAAATGTAAGGTTTTAACCAATCACGTCCCGCTTCTGTAGA | ATTGTATTAAATAAAATAAAATAAAATAAAATAAAATAAAATAAAATAAA | TAAGGTCGCATGCCCGCAACTGG | 9 |
| **TCTTTAATACCTGTTGCGTA** | scaffold_1 | 66501 | + | GTCCTGAGATTGAAATCTCATAGTTAAGTTGGCTTCCTTTTGCTCTAATT | TCTGTCAGCAATGTTTCTCAATTCTCATTGACAAAGCCAGAAATATGCAA | TCTTTAATACCTGTTGCGTATGG | 9 |
| **ATGATGACCGAAACCTGCCG** | scaffold_1 | 221958 | + | TGTACAGCTTGAAGGAATTGCTCTTTGCCTTTATCTTCCTCTCGTTTTAT | AATTAGTGTTGATTTCTTGCTTACCATATGAAACCCCTCGCAAATCTTTA | ATGATGACCGAAACCTGCCGAGG | 10 |
| **TCTACATAGCAAGGTATGGC** | scaffold_1 | 291160 | + | TAGAGTCTAATGAAGTCTGGTGGAGTTGTACTTTACCAATTTTCTTTGTT | CATGTGCTACTATTATAGGCTATTTACAAACACTGCAGCCCCTTACTTCT | TCTACATAGCAAGGTATGGCTGG | 7 |
| **TAACACACCAGATTGTGTGT** | scaffold_1 | 1449658 | + | GATAATGACCCATTAGGCATATCTGCTTGAGTTCAAGAATTGAAGAGAGG | GATTTGAATAAACACTGGAGTTGCTGCTAGAACAATCTTGCCAAGAGAAT | TAACACACCAGATTGTGTGTTGG | 5 |
| **AGGGCGGGATAAACGTTGCT** | scaffold_1 | 1450248 | + | GTGAGGTTGGTGCTGTCACTCTTGAGCCCATTAGTTTCTTGTTGTTCAAT | GAAAACTTCATCCTTCCGAGGAGTCCCACTGTGTACAATGAGTTTTAGTG | AGGGCGGGATAAACGTTGCTAGG | 5 |
| **AGGAGGTCTTCACTGCCGTC** | scaffold_1 | 2269642 | + | CGTCCTCTCCATCTTCTCCCTCAATATGGGCTTCTGAATCAAGTTCCAAG | TTCACCATTTACCTCAGCCGTGGTAGACAATCGACTACCATTGGCTTCTT | AGGAGGTCTTCACTGCCGTCAGG | 6 |
| **CAAATTTGCCTCGACGATGC** | scaffold_1 | 2595134 | + | CGACTTTAAAAATTTCGGCAAAAGGTAATCCATTGTCTCTTGCAATGCCC | TGACTAACGATGCTCCATTAGCCATTGGCATACCGCCATTAACGACATGA | CAAATTTGCCTCGACGATGCGGG | 6 |
| **ATATGGATCAATACGCCGAA** | scaffold_1 | 2610747 | + | TGGTTCTGATAACCCATGAAAATTTGCTGATAAAATTCCCGATTAGAACA | CATTATAA | ATATGGATCAATACGCCGAATGG | 6 |
| **CGGCAGTGAAGACCTCCTCT** | scaffold_1 | 341118 | - | GCCAATGGTAGTCGATTGTCTACCACGGCTGAGGTAAATGGTGAACCTGA | AACTTGATTCAGAAGCCCATATTGAGGGAGAAGATGGAGAGGACGAAGAT | CGGCAGTGAAGACCTCCTCTTGG | 6 |
| **GCCTCTACCTAAATCTCTCG** | scaffold_1 | 1182499 | - | TCTTTCCTGTCAGCAATAATATCTATTCTTCACGACTTTGAGAAAGTTCA | ATTACAGATAAACTGGAAGGTTTATTGTATGAATATTGAAAACACATAGA | GCCTCTACCTAAATCTCTCGTGG | 6 |
| **TCTCAAGGATCCATCGCGTG** | scaffold_1 | 2509883 | - | TATAAACTCTAAGTACTAGCCCTATGATTATCAAGTTTCTTTCCAATTGG | GACTGGAGTGTCAAACTAAGTTAATCCAATAGCTCCGCAATTATCTTCGC | TCTCAAGGATCCATCGCGTGTGG | 5 |
| **CATACTCGATACTTTCGTAC** | scaffold_1 | 2510218 | - | CCCTCCCTAATGTAAGACATAAACCCTCATTAAACATGCATACGCAAGCC | ACATATCAAACCCGTTCATCAATTGAGCAATATTGCCAGGGATTTGCATC | CATACTCGATACTTTCGTACCGG | 5 |
| **CAGCAGCCATATCCGGACCC** | scaffold_1 | 2543412 | - | ATGCAAGCACACTGTGAATGACTAGGGCCAAGAAAAATGACTGCCTGCAA | GCGGCTGTATGCCATGTCGTCATAGTCGAGTAGGAACTGGAAAGAAGTAA | CAGCAGCCATATCCGGACCCAGG | 9 |
| **CCATGTCGTCATAGTCGAGT** | scaffold_1 | 2543446 | - | AAATGACTGCCTGCAACAGCAGCCATATCCGGACCCAGGGCGGCTGTATG | AACTGGAAAGAAGTAACTCAATTATCATCACTTCAACAGTATAAACAACT | CCATGTCGTCATAGTCGAGTAGG | 9 |
| **CAGGACAGAACAGTATTGCA** | scaffold_1 | 2544321 | - | TAAAGAAATTAGAGCAAAAGGAAGCCAACTTAACTATGAGATTTCAATCT | AATTTCGGTTTTGCTTTCTTTCCTTGTTTTCCTGGTGAAGTAATGCCTAC | CAGGACAGAACAGTATTGCATGG | 9 |
| **TGCGGGCAGTCTCGGTAGTC** | scaffold_1 | 2610218 | - | GACCTCGGCCGAGGGCATGTGTTAACTTTCCTTATTGGGATTAAAGCCGG | AGTTTGCTCTACCAACAAAATTCAATTACGGGAGGTGCTTACTAAGCTGG | TGCGGGCAGTCTCGGTAGTCCGG | 2 |
| **CGCAATAGAATCCTCTTGTG** | scaffold_2 | 209 | + | GCACTGAGGTTTAGAGACACTTTTGAAAGACTCTCTTCAATAGTCAAAGC | TCAGAGGTTTATCCTCACCCAGAACTAATGATATATCAAGTTCAAGAGTC | CGCAATAGAATCCTCTTGTGAGG | 5 |
| **CCCTCCCGCTACAGGTGCAT** | scaffold_2 | 780131 | + | CATGAAAGAATAGGAAGAAAGAGACGTACACAGGAAAGAGAGTACATTTA | AGCTACAGCGGCGTGCATCGTATACATAGTTGGTGATGAAGGTGGTGAAA | CCCTCCCGCTACAGGTGCATAGG | 8 |
| **CCCACTCGCTGCTATCTCCG** | scaffold_2 | 1986656 | + | TGCCTCTTTGTACTTTCTGAACCAAACTGTAGTTTCATTAACTAAACCAT | AGATTGCTTTTCAACCACTGCTGGTTATGTGTGGTGTCCCGTGTACTATG | CCCACTCGCTGCTATCTCCGCGG | 8 |
| **AGGCTCAGCACGGTGTAATT** | scaffold_2 | 2003926 | + | ACTAATAGTATAACTAAATTAGACTGTATTTCAAATGAGCCGCCTTTAGA | AAATTCCATGGCATGTATAAATAACGGATGTGTGAGTTCAAGATGCTCTT | AGGCTCAGCACGGTGTAATTTGG | 6 |
| **TCTTCCGTGATCCAGGTGTT** | scaffold_2 | 2003995 | + | TTGGAAATTCCATGGCATGTATAAATAACGGATGTGTGAGTTCAAGATGC | GTTTGTTTGTGTCTGTACTTCTATTGTTTACAATAGACATGGTTCACAGA | TCTTCCGTGATCCAGGTGTTTGG | 6 |
| **CGCGGAAAAGTTGTTGGCCC** | scaffold_2 | 2004177 | + | TTATTATATGCTGAGGCGGAAAATTCAATCATTTCTTACATAATATACTA | AAACACCCAACGCGAACATCCAGCTAGCCACTTTTGGCGTATATAGATGT | CGCGGAAAAGTTGTTGGCCCCGG | 6 |
| **ACCCCTGAAAGGACCCTCAG** | scaffold_2 | 2413394 | + | ATTTTAAAAGTACGTGATCTCGCTCCTCTGCGAGGTGTCCGGGGTAGTCG | TGGCGCAGTGCAGCTGAGAGGTGGGAGTGGCGCATGCGTTTTGAGGGCAT | ACCCCTGAAAGGACCCTCAGAGG | 5 |
| **ATACGCCAAAAGTGGCTAGC** | scaffold_2 | 457782 | - | AAGAGGTTCTCAATGTGCCTCAAGATCTCAAGCCATGGTTCAACATCTAT | ATGTTCGCGTTGGGTGTTTCCGGGGCCAACAACTTTTCCGCGTAGTATAT | ATACGCCAAAAGTGGCTAGCTGG | 7 |
| **TGGCACCTGTAGGGGACTTA** | scaffold_2 | 856637 | - | AGATCTGCGTTGTTCTTTCGCATATCGCGTCGTATTTGAGGGCCGCATTT | ACAAATAATATACGTGAGGTATGAATGGAAAAATTAGTAGCACAAAATGT | TGGCACCTGTAGGGGACTTAAGG | 8 |
| **CACCGTGCTCCCCTTTCGCA** | scaffold_2 | 2438921 | - | TTGACATGTTACCTCGATATCTTCTTGTCCTGTCATGAACACAAGAATGT | GGATTCGATAAATGTATCTTAAGCGCCTGTTTGACAGCAGAGTCAATGTA | CACCGTGCTCCCCTTTCGCATGG | 10 |
| **GCCTAGTGGTCAGCACCGAA** | scaffold_2 | 2439815 | - | CAACTTGTACAAGTCACCAGACTTGTCTCGGACAATCTCAGCTATATTCT | GAATCGAGAAATGGTGGCGTGAGTCTGTGTACAAAAATATCATTAACTCC | GCCTAGTGGTCAGCACCGAATGG | 8 |
| **TTCGTGCAGGTTGTCTTACT** | scaffold_2 | 2439919 | - | CAAAAATATCATTAACTCCAGTAGTGTCGATATTCTTACTGAGTATCTGC | ATCTACCTCATAATCATCAAACTGTTCAAAATATGACTCTTCAGCGTCGT | TTCGTGCAGGTTGTCTTACTTGG | 8 |
| **TTCTAGTGGCACAGTGCCAT** | scaffold_3 | 1848573 | + | TCACAGGTTTATGCTGTTTGCCCGTCTCTGTAAAATATCTCGGAACAGTG | GCTAATTTTGCTGACAATTTGGCCCGTTCGGCGTGTATGCTTGTCAGTAA | TTCTAGTGGCACAGTGCCATTGG | 5 |
| **TGCCCCCTTCCAAATGGGCA** | scaffold_3 | 590335 | - | GTCAAAGAATGGGCTTGACAACGTTGATCTTAATTTTCCTACATATAAAA | GACAACAAATACAGTTGTCGCGGCACAACAACTTCTGACCAAATGCGCCA | TGCCCCCTTCCAAATGGGCAAGG | 8 |
| **CACAGCCCGAGCGCGTCGTG** | scaffold_3 | 1829640 | - | GGGCGCGTGCCTACCCCATTGGGAAAAACATCAACACGCGGCGCCGATCA | CAAATGCATTAACACGTGACTGTGGCGCATTGAAACTCGGTGGGAAAAAC | CACAGCCCGAGCGCGTCGTGTGG | 9 |
| **TCCGCTTTCACGGGCAGACC** | scaffold_4 | 39454 | + | CAACACTTTCATTTCATTTGTTTCCGATATAGTTGTAACGGCTATCACGG | GTTCGACTCCCGGTATCGGAACATTTTAAGCCTGGTTAGCTCAATCGGTA | TCCGCTTTCACGGGCAGACCCGG | 5 |
| **CCGCTTTCACGGGCAGACCC** | scaffold_4 | 39455 | + | AACACTTTCATTTCATTTGTTTCCGATATAGTTGTAACGGCTATCACGGT | TTCGACTCCCGGTATCGGAACATTTTAAGCCTGGTTAGCTCAATCGGTAG | CCGCTTTCACGGGCAGACCCGGG | 5 |
| **ATCCTCTCTATGGACTCCCT** | scaffold_4 | 57149 | + | AGCTGTGGGCATTCCACCCACGTGTTGCAGCTTACTTACGTATACAGATT | TACTGGCTGAAGTATGGGTACGTGGCACCTTGTGATGCGCCGTAACGTCC | ATCCTCTCTATGGACTCCCTAGG | 9 |
| **CCTAGGTACTGGCTGAAGTA** | scaffold_4 | 57166 | + | CCACGTGTTGCAGCTTACTTACGTATACAGATTATCCTCTCTATGGACTC | GTACGTGGCACCTTGTGATGCGCCGTAACGTCCAGCTACGATGGATGCTG | CCTAGGTACTGGCTGAAGTATGG | 9 |
| **ACCTGCGCAGCGGAGCGGGC** | scaffold_4 | 353163 | + | AGAAGCCCATTTTCGGGCGAGAAGGGGTAACGGACGCCGGGGTTCTCGCC | GGAAAAAGAAACGAGGCGCGGGCGGTCACGTGTGCAGTCAAGTGTTAGAC | ACCTGCGCAGCGGAGCGGGCCGG | 8 |
| **ACTGTGGTCACGTGTGCATG** | scaffold_4 | 353265 | + | GTGTGCAGTCAAGTGTTAGACTGTGGTCACGTGTGCAGTCACGTGTTAGA | CGGTCACGTGGTTTTAGTCAGAGACTTTGTTATTTCAAAATACCTCACTG | ACTGTGGTCACGTGTGCATGCGG | 8 |
| **TACTAGTGTCAGAAGCCACC** | scaffold_4 | 824796 | + | GCCAATAGTTCAGTCTATAACATTCAAAAACGCTAATCTATATTATAGTT | TAATAAACTTAATCATTATTATAATCCTTATTATAGATGCACGATTAAAC | TACTAGTGTCAGAAGCCACCAGG | 6 |
| **GTAGCGTGTAGTGGTTAGAC** | scaffold_4 | 865276 | + | TTTCCAAAATGATACGCGGTCGTGCTATTATGGTTTCCAAAAGTTTAATG | TAGGTCCATCTATACATGTAGAAAATAATCATTGTGAATATAAGTTATAA | GTAGCGTGTAGTGGTTAGACAGG | 10 |
| **TACTTCCCAAAGGGGAGGGA** | scaffold_4 | 867457 | + | CAAGTGGAGATTTAGAAACAATGGAAAATGTAAATGATGCCGCTCTCGGA | AATATTGTCTAGCTCCCTCTCAGTAAAGGGAAACGTCGTTATCTGCCAAA | TACTTCCCAAAGGGGAGGGACGG | 9 |
| **TATTAGTTCACACTGTACCC** | scaffold_4 | 881941 | + | TTCTGACTAGTGACCGTGATATTACCACCAGTGTTATTATTAGGGCCATT | GGAATGCAGTACCAAAAAGTTGTTTATCATTCAACAATTGGAAAACCTGG | TATTAGTTCACACTGTACCCTGG | 8 |
| **GGACAAATTCAGATCGTCCG** | scaffold_4 | 882379 | + | TTATTTGTATTTGTCTTACTCAAGTTGGTATTTATTGTTAGATTCTTCAA | AGTTATTGGATGCATAGAATGACATGATTGGAGATGATGCTGGAACAAAA | GGACAAATTCAGATCGTCCGAGG | 8 |
| **TACGTCAGACCGGGCTTACC** | scaffold_4 | 905171 | + | CTGGCATTGAAGAAGTACCTAAGACGGAAACCTGAATAATGGAGTGAGTT | ACTATTCCACAAGCCAATAGAAGTCACGAGCATCTCCTTTTGTCTTCCCA | TACGTCAGACCGGGCTTACCAGG | 11 |
| **AGTCGAGATCACGACCTAAC** | scaffold_4 | 911509 | + | TCCTGATAAAAATTGGAGTGAATTGGACTCTCTCTCTAGATTGATCACAA | GAATTCTTTCTCAATCACAAAGTATCGGCAATTGCTGACTCCAGCGTTGA | AGTCGAGATCACGACCTAACTGG | 11 |
| **GAGAGACAACGCGTGTCCTC** | scaffold_4 | 911897 | + | GAAAGTAGAAAGAGAATTGAGTTAGATACAGTTGTAGAATCTTTGTTGAT | TTGTGATTAGCCACAGTGATAAGAAGTGTATATACACATATGATCTGGGT | GAGAGACAACGCGTGTCCTCTGG | 10 |
| **CTCCGTATGGCTAAGAGTGC** | scaffold_4 | 916024 | + | TAAGCGTATAGGTGGAATCCAGTTGATTTCTATCAGGTTAACCACATACA | AACCTTTGTGTAACAGTAAGGTAAAACTTTCGACATACAGAGTTTCCGTG | CTCCGTATGGCTAAGAGTGCAGG | 10 |
| **GGGGCGGATTGACCTAATAC** | scaffold_4 | 980623 | + | AGCTCAACGAGAAGGTGGATGTACTCTCGAGGAGATGGAAACGTATAGAC | AGGTGGATCAAGTACGCTGTGCATGGTCCCGCAGTATTTCGCAAACTATG | GGGGCGGATTGACCTAATACAGG | 6 |
| **GGTCCTGTAGAGGCATTGTT** | scaffold_4 | 1056953 | + | GGATCCCAGACTAGTGTTGGCCAGACCGTCCGCCATATGGTCCACCTCGG | TATACTCTTGGAAAGAAGGAAAGGACAGAGGAAGGAACTGGATTGGCAAT | GGTCCTGTAGAGGCATTGTTGGG | 8 |
| **GTCGTGTGCTCTGCACATGT** | scaffold_4 | 1060226 | + | TGTTATATCCACACACAGGGGCAGCCACCCCTCCACGCGTGGCGTCGTCT | TATAGACGGGCCTTGCTGTGACACCACAGAGGGGGACATTGACTTTAGTG | GTCGTGTGCTCTGCACATGTTGG | 8 |
| **GTGATGACGCCTGTGCTGGC** | scaffold_4 | 1098563 | + | ATCAGCCCTCGAGACACCTATAACATTGGGGTTCTGAACAATGGGTGATG | AGAACTTGAGATTGTATATGTAAAGATTGTTGTTGATGGAGATGCTGTTG | GTGATGACGCCTGTGCTGGCAGG | 9 |
| **CACTATTCGTCCCAGCTGAT** | scaffold_4 | 1128671 | + | CGGTGTTTTGTTGTGGAAGCATTGAAAGAAATCACTGAATATCCTGAAAT | CTCTAAGTTATACAGAATTTGGGAACATGGAAATCTTCAGAGACGAAGTG | CACTATTCGTCCCAGCTGATCGG | 3 |
| **TGCAGGAGGCCACACCACTC** | scaffold_4 | 1129658 | + | AAGCAATTTCACCTTCTAGACAACTCCTCCACAAATAAATTGGTTACATA | CACAGATTTTATGCACATTGAGAGCACACTATTTGACTTTTGACGCGTCT | TGCAGGAGGCCACACCACTCTGG | 2 |
| **AACGGTGCCCGACAGGAAAC** | scaffold_4 | 2669 | - | TAAGCAAATGTTCTTTAAAAACGCCTAGAGATGAACGCTTGACTGTATTC | AGAATGTCTACAAGTCGTTGAATGAATCATGAAAATATGACCTATCTTAG | AACGGTGCCCGACAGGAAACAGG | 2 |
| **CGCGTGGCGTTCTTTTGATA** | scaffold_4 | 4253 | - | AAAGCCTTGGTTATGTTTAGTTAGCTATCATAATGTAAATCATCTATGAG | CATATACATAGTCTTCAGTATTTGCTTAATTAACCGATTCTACACCATGC | CGCGTGGCGTTCTTTTGATATGG | 2 |
| **CTGGGACGAATAGTGATTTC** | scaffold_4 | 5743 | - | CTCTGAAGATTTCCATGTTCCCAAATTCTGTATAACTTAGAGCCGATCAG | ATATTCAGTGATTTCTTTCAATGCTTCCACAACAAAACACCGAATCAATA | CTGGGACGAATAGTGATTTCAGG | 3 |
| **TATCCATCGTATAAGCGTAT** | scaffold_4 | 6784 | - | TGAGGCTGTTATCAATCCTAAGTGAGGGTGTGTTGTCACATCAAATGCAG | TACTGTGGCCCCAAGAGCGATTCTTGAGACACTGTAACCAATACAGGCTT | TATCCATCGTATAAGCGTATTGG | 4 |
| **TTCAATTTCGAACACTGCCG** | scaffold_4 | 7568 | - | AAATATAGCAAAAGCTTCCCCCGGATTCCACTAAATCACTGTGAACATTT | CATACCTATGAGGAGCTCTCCTGAATAACCTGCCATACTTCCCTAGGATT | TTCAATTTCGAACACTGCCGGGG | 4 |
| **TTCTCCTCAGCCCGGATCAT** | scaffold_4 | 35960 | - | GCTGATCCAAATAATACCAGGGTCAATTCATCTTCTATTACTACTAGGTT | GAAACAAAATGTTAAACTACTACCACAAGAATAGTGGAGTGAACAATGCC | TTCTCCTCAGCCCGGATCATCGG | 9 |
| **TCTCCTCAGCCCGGATCATC** | scaffold_4 | 35961 | - | CTGATCCAAATAATACCAGGGTCAATTCATCTTCTATTACTACTAGGTTT | AAACAAAATGTTAAACTACTACCACAAGAATAGTGGAGTGAACAATGCCA | TCTCCTCAGCCCGGATCATCGGG | 9 |
| **CCTATAGGGTCGAAGTCCGG** | scaffold_4 | 52950 | - | TGATGTCTGACAATGAAAACCAGAAAAACAGTACATCACCGGAATTCACA | TGTTGCCGAAAACACGATGCATACATTCTCAAACATGACTCCAATTTTAA | CCTATAGGGTCGAAGTCCGGTGG | 8 |
| **GTCGGCCGATAGTGCTAACA** | scaffold_4 | 53294 | - | AACGAACGGGAAAGGGCAGATAACCCTAACAATGATCCTTTTAGTAGAAG | AGGTCCCTATATTAAATGGGAAGGAAAAGATAAGAGATGATATGATTGGC | GTCGGCCGATAGTGCTAACATGG | 7 |
| **ACGACAGACGACGCCACGCG** | scaffold_4 | 74198 | - | CCTCTGTGGTGTCACAGCAAGGCCCGTCTATACCAACATGTGCAGAGCAC | AGGGGTGGCTGCCCCTGTGTGTGGATATAACATACACACACACGGACAGA | ACGACAGACGACGCCACGCGTGG | 8 |
| **ACAGACGACGCCACGCGTGG** | scaffold_4 | 74201 | - | CTGTGGTGTCACAGCAAGGCCCGTCTATACCAACATGTGCAGAGCACACG | GGTGGCTGCCCCTGTGTGTGGATATAACATACACACACACGGACAGAGAC | ACAGACGACGCCACGCGTGGAGG | 8 |
| **GGACCATATGGCGGACGGTC** | scaffold_4 | 77483 | - | TCCTTCTTTCCAAGAGTATACCCAACAATGCCTCTACAGGACCCCGAGGT | CCAACACTAGTCTGGGATCCGGCAATTCAAACGACTCAGATGACAGATCT | GGACCATATGGCGGACGGTCTGG | 8 |
| **GGGCAAAGCACACTTAGCCA** | scaffold_4 | 161048 | - | TGAGGAATCGTATGCTAGGTTTATCGAGAAGTATGATATATTACCGGCAC | CCGGACATAGAATTGCAGAATTCAAAGGTGAAGATGAAACGCTACTTCCA | GGGCAAAGCACACTTAGCCATGG | 7 |
| **TAGTCAGTAAATCGACTCAA** | scaffold_4 | 217652 | - | TTGTGGGCCAGTTTGGTGTTTACTACTCTTCATTTTAGAATATTCCGATT | TGGGCTATCCAGAACGTGTCATCCTGTTCAGATTCATTCCTTTGATAGAA | TAGTCAGTAAATCGACTCAATGG | 10 |
| **TTCCTGCACTCTTAGCCATA** | scaffold_4 | 218380 | - | TTCACGGAAACTCTGTATGTCGAAAGTTTTACCTTACTGTTACACAAAGG | AGTGTATGTGGTTAACCTGATAGAAATCAACTGGATTCCACCTATACGCT | TTCCTGCACTCTTAGCCATACGG | 10 |
| **CTAGAGTGACTCGTGTACCC** | scaffold_4 | 266741 | - | AATATTCCCTCAAAGTGTACATCAACAATACGTTCAAATGGACAGCTGTT | ATAAGCAGGTATATCTGACTTGGTCTGGCCGTGTCCTATCAGACGATAGT | CTAGAGTGACTCGTGTACCCTGG | 9 |
| **TACATGTGACTGTTCGCGTG** | scaffold_4 | 307070 | - | CCAAATCATTAATAAAAGGAACCAATTATAATACAATATGGATAGCTCAA | CGATTTTGTGGAGTGTTGTTCCTGCTCCAAAAGATGTTCCTGGTGCTAAT | TACATGTGACTGTTCGCGTGCGG | 6 |
| **GGACGTTACGGCGCATCACA** | scaffold_4 | 1077207 | - | TTTAACTTGTCTTCAACACCAGCACAATGGAAACAGCATCCATCGTAGCT | TGCCACGTACCCATACTTCAGCCAGTACCTAGGGAGTCCATAGAGAGGAT | GGACGTTACGGCGCATCACAAGG | 8 |
| **TACCTAGGGAGTCCATAGAG** | scaffold_4 | 1077255 | - | CTGGACGTTACGGCGCATCACAAGGTGCCACGTACCCATACTTCAGCCAG | ATAATCTGTATACGTAAGTAAGCTGCAACACGTGGGTGGAATGCCCACAG | TACCTAGGGAGTCCATAGAGAGG | 8 |
| **CCCGGGTCTGCCCGTGAAAG** | scaffold_4 | 1094951 | - | CTACCGATTGAGCTAACCAGGCTTAAAATGTTCCGATACCGGGAGTCGAA | ACCGTGATAGCCGTTACAACTATATCGGAAACAAATGAAATGAAAGTGTT | CCCGGGTCTGCCCGTGAAAGCGG | 5 |
| **GTGGAGAACGCTCCCTACCA** | scaffold_5 | 513648 | + | CGATTAATGTAACGATATAGGTTCAACTGTACAACAGGAGAGTCACCTCT | GATCGCCTTCCTCTTTGTCGTGTTTCCATGTAGCAGGACCAGTGCAATAC | GTGGAGAACGCTCCCTACCAAGG | 9 |
| **GAACGCATCTTAACCGAATG** | scaffold_5 | 586594 | + | GAGAAGGATTGGAATCTATCGGGTGTATCGACGAAGTGGACGAACCAGAA | CGCTTCCAGTTTGGTAGAGGCACTATCAGTCTTGAGGGTTGGTGACGAGC | GAACGCATCTTAACCGAATGAGG | 8 |
| **GCACTATCAGTCTTGAGGGT** | scaffold_5 | 586636 | + | AACCAGAAGAACGCATCTTAACCGAATGAGGCGCTTCCAGTTTGGTAGAG | TGACGAGCCAGGATAAGATGAGTGGGCATCGTTTGGATGTCCTGAAGTAG | GCACTATCAGTCTTGAGGGTTGG | 8 |
| **ACCGGACCCCAGAAATGGGG** | scaffold_5 | 600180 | + | GTTTGTTTCTCAAGTCCAAGACAGCAGCTACGGGGAGAACGAAATTAGAG | TGAACACATACTTGAGGGTTTGTTTTATTAAAGATTTTCTGTGAGAGTTT | ACCGGACCCCAGAAATGGGGTGG | 8 |
| **CACACGATCAGGGCCTCTGC** | scaffold_5 | 709050 | + | CAACAACAGCAACAACAGCAGCAGCAGCAACAACAGTGTGTACCACCACA | CACAAACCGAAAAACGATCTCAAGACCAACCTTCTCTTGCGAAGCGAAAT | CACACGATCAGGGCCTCTGCAGG | 8 |
| **TAATCTTTCGAGCGTGAGGC** | scaffold_5 | 990375 | + | TATGCATAGCTGATATATTTTCTTATTCAAATACAAGCTGGCCGGTTTCT | CATTAGTAGGACTAACCATCTAAAATATTTTCCACTTACTATTACTATAA | TAATCTTTCGAGCGTGAGGCCGG | 3 |
| **GGGAATGCCCGGCCAGGCTT** | scaffold_5 | 579919 | - | AGTGGCCGGCACTGTATGTTATGGCACTCGCACACAACGTGTTCTCCACA | ATTATGCAGAATGGTCTTTCCCACCAAATCCACTGACGTCTTCTATGCAA | GGGAATGCCCGGCCAGGCTTTGG | 7 |
| **AATACGGCGGCACGCCTAGA** | scaffold_5 | 941914 | - | TACAAGTGTATATGAGGGTTAACAGGCTTGCTGTGCGGTGTGCGGGTGGG | GGCGCAGAGGCTTTGTCAGCCCTCTTTCTGTGCACGTGATACCATGTTTT | AATACGGCGGCACGCCTAGAGGG | 10 |
| **GGCAGATGTTCTGAGCATGC** | scaffold_5 | 1089182 | - | GTTGGGCTGCCGGAGAGAGTCCTTGCTCTGAATGCGGTTCTGGAGAGTCT | TAGTGTCGAGACGTGGCAATGTATGTATATATCGTATATATATATGTATC | GGCAGATGTTCTGAGCATGCTGG | 8 |
| **GGCACCTTTTCGTCGCCACG** | scaffold_6 | 155117 | + | TAATACTTGGTTGGAATGTAGATTGGCTATTTACAAACGGTGGAACCTGG | GTTGGTAGGACTGTTCAGTAGATTCATAAGATTCAGGCGTCCGTCTGCTG | GGCACCTTTTCGTCGCCACGGGG | 0 |
| **GCACCTTTTCGTCGCCACGG** | scaffold_6 | 155118 | + | AATACTTGGTTGGAATGTAGATTGGCTATTTACAAACGGTGGAACCTGGG | TTGGTAGGACTGTTCAGTAGATTCATAAGATTCAGGCGTCCGTCTGCTGC | GCACCTTTTCGTCGCCACGGGGG | 0 |
| **CGGATTCCAGTGGTGAATGG** | scaffold_6 | 188953 | + | CTTACATTGAGGAGGAACTTCGGCCAAAATGATGGACACCACCCGTCCTT | GTGCAAGTTACCAGAATGACAGGGACATTAGTTGGAACATGATTAAGAAG | CGGATTCCAGTGGTGAATGGCGG | 0 |
| **ATACAACGTGGTATCCACAC** | scaffold_6 | 343479 | + | AACACAAGTACCATAAGGCCTCCCAAATCAGATACATCAGGCACCAAACT | CACTATACCCTTTACTTTCTTTGATAAATCAAAAAGGCATTATTGATTAC | ATACAACGTGGTATCCACACAGG | 0 |
| **TTGACACCCCGTCTCTGTTG** | scaffold_6 | 437382 | + | GTTGTTGGTGTTTTCAATTTCGGTTTCACCCTTGTTGTAGTTGCTACTAC | CCAATATACGGGAGTTTCGAATTGGAGTAGAAGCTAAAGAGAGAAACAAG | TTGACACCCCGTCTCTGTTGTGG | 0 |
| **GTAAGTTGAACCGACCCAAC** | scaffold_6 | 553808 | + | GTCGCCTCTGAAATCATTAATGTATCTTTATTAACTACACCACCCAAATT | CAATACCTGTTAAAGTACCCAATGGCGCACCTATGAGTAAAGACAACTTT | GTAAGTTGAACCGACCCAACAGG | 0 |
| **GAGACACATAGCTGCACTAC** | scaffold_6 | 620880 | + | ATAGGAATAGAGTCATTACCAACAAGATCTGAGCTATTGAATTTGATCTG | TTAGTGAAGAAACAAAAGAATCTTTTGTCGATAATGTTGCAACTGAATTA | GAGACACATAGCTGCACTACCGG | 0 |
| **AACCCAGGCGGTTTGCTGAA** | scaffold_6 | 727186 | + | AACCATGATGTTTCGACAAGTAAGTGCAGCCGTCACCTTGGTTCTTCTGC | AAACTTACTCAAATTCCCTCGTGAACTGTATGAAAAACAACACCGACATT | AACCCAGGCGGTTTGCTGAAAGG | 0 |
| **CTTTGGTAACTCGGCTCGCT** | scaffold_6 | 727693 | + | CGGCTTGTTATTGCAGGCTTGGGGTTGCTTGTGTCGGCCTTGCTTTCTAT | CGTCAAGAATGTCTGCTGTTGCAATGTCTCTCTTCACATACTTTCAATCC | CTTTGGTAACTCGGCTCGCTCGG | 0 |
| **CCCTGGTTCGCAGTGTACGC** | scaffold_6 | 740736 | + | CATCCATCTTCCTAGCCCTTACCAACGGGGTATAAGCTTCCACTAGTAGA | TCAAATCATTCCTTGTCAACCATGGATGAAGTTCGAGTTGGTTCACTGCT | CCCTGGTTCGCAGTGTACGCGGG | 0 |
| **AGCGAGGACGGTGACGTGAT** | scaffold_6 | 948465 | + | GAAGAGGGCGACCGGGGTGCATTTACATGTGATGGATAAGCGATTCCACG | TGATAATAGGGAGCGTCCGTGTGAAGACAAGGCACTCCAACGGGTCCATG | AGCGAGGACGGTGACGTGATTGG | 0 |
| **GCACGAAGTTGAGCCCCACA** | scaffold_6 | 2430 | - | TGTAGCGTTCAAATGTACAGCGGTCCTTGTCAGGTAATATACTCTTGGTA | CTTCTCGCCATGTGCACTTTTCAATTTCCAATAGAAAGGCACGCTCAAAT | GCACGAAGTTGAGCCCCACAAGG | 0 |
| **ATTCTGTGCAGTCCGTGTAG** | scaffold_6 | 2746 | - | AGATAGAGTTTGTCGGCTTTCTCTGTATGTGCCATTGATTTGATTTCTCT | CCACTTGATAGAACTAACAATACCAGTACCAGTACCAATGTCAATACTCG | ATTCTGTGCAGTCCGTGTAGAGG | 0 |
| **GTTCTTCTGTTTACGGATGT** | scaffold_6 | 5746 | - | GAGAGCCGATGAAATTAACCAAATTTTCGATGCAATTTCGTACGAGAAGG | CCAATTGGTTAGGTGAAGAGACTTTCATTAAGGGTGTCTCTCACTATCTA | GTTCTTCTGTTTACGGATGTTGG | 0 |
| **GTTGCCTTATGAGAATACCC** | scaffold_6 | 23221 | - | CTAGCTTTAATAAGGTTTAAATTCTGACTCTCTGGAAAATAAGAATGGTA | TATGATTGATGTCTTGAGTATACGTGTATCGTTTACACTTAAACTTGTCT | GTTGCCTTATGAGAATACCCCGG | 0 |
| **GTAACTCTTGATCGAGCGTT** | scaffold_6 | 147212 | - | CTGCAATGACATCTGGCAAGCTAGATGACTTCTCTTGCTCCTCAGAAATG | AGCAGGTGCATTGTCAACAACCTCTGTGCTTAATGGCACAACCTCATCTT | GTAACTCTTGATCGAGCGTTGGG | 0 |
| **TATCTGATGCCTCCTGCAGC** | scaffold_6 | 159402 | - | TTGAAGCGTATTACTGAGAGACAGAGAAACCTGTCTAAAGAATGAAAGAA | CCAAGAAATACCTCGGGGGTGGGACAAAATGGAGCTAGTTATCATATAAG | TATCTGATGCCTCCTGCAGCCGG | 0 |
| **GACGCATCACCGCCACCAGC** | scaffold_6 | 171757 | - | AAGATAAACACGTCGGACTAAAAGAGAGCGGGGTTCAGCAGATTTATGAT | CAGCCAAGCGAACACATCCACGACTGAAAACGTTCTTATAGAAAACGTCA | GACGCATCACCGCCACCAGCGGG | 0 |
| **TAGCCACAGGCGTCCAAGTC** | scaffold_6 | 225348 | - | GAGGCTTGCAAACAGCCCAAGATTCGTTTTCTTGGTGAGAATAATCAACC | GAAAGTGTAAGCAAATACCTGGAATTTTGTCCACGTATGACTTGTCGATG | TAGCCACAGGCGTCCAAGTCTGG | 0 |
| **ATCGCCAGCACCACGATTAC** | scaffold_6 | 550627 | - | AAGAATTCATTACGCGGGAAATCTTGAGTATCATCGTCTTCTTCGCCAGA | GTTCGTCATCCATGTAGGAATCCGTGGTATCGAGTCCGTCTGGAGATCTG | ATCGCCAGCACCACGATTACCGG | 0 |
| **CAAGTTTGGCTGTGTCCTAG** | scaffold_7 | 392267 | + | GCGTGGAAAGATGTAGATGACGATGATGAGGCCAGAGAGGCTGAGGACGG | AGAAGATGGAGGAAATGGAGGAGAAGGACGAATTTCTCGAGGTAGGTCTA | CAAGTTTGGCTGTGTCCTAGAGG | 5 |
| **TATCTCGAAAAGCAACGGGA** | scaffold_7 | 392591 | + | ATTCTTGCATATTGAGGTTTGAGGTTGTAGAAATTGATCTGGGTTGAGTT | GACTCCCCCGAGAGGGCATGTAATCATTGTGTGTTTCGTCGACAAGATAT | TATCTCGAAAAGCAACGGGAGGG | 5 |
| **ACCGAGGGGGTGTACTCCGG** | scaffold_7 | 392682 | + | TGTAATCATTGTGTGTTTCGTCGACAAGATATACAAGGGAGAGATGGCGA | TTGGTGAAGGTATTCTGAGACGCTATAACTGGGAGGTGGGGTAGAGGTAT | ACCGAGGGGGTGTACTCCGGCGG | 5 |
| **TCTGAGACGCTATAACTGGG** | scaffold_7 | 392718 | + | GGGAGAGATGGCGAACCGAGGGGGTGTACTCCGGCGGTTGGTGAAGGTAT | TGGGGTAGAGGTATCCATTTCTCTGGGTAGATTCTAGATGGGGGTGCAGT | TCTGAGACGCTATAACTGGGAGG | 5 |
| **GTACGTAATACAAGTTAATG** | scaffold_7 | 435902 | + | ACGAATACATCCCCAGCACCAAAGTAAGTATAATGTTTAACTTCCCCAGT | TTGACGCCATCTTAGCACATAGTACGCGCCCTAAGTATAGTATTACGTAA | GTACGTAATACAAGTTAATGGGG | 4 |
| **AGCTTGTACACAGCCTAGAA** | scaffold_7 | 469134 | + | CAAATCACCCAGTATTTGGCTGAAGGCGGCTATGCTAATGATGGCATCAT | TTGCCGCCATTAGCGTTGCAAAAAGGGTTGCTGAGGAAGTCAACTGTAAA | AGCTTGTACACAGCCTAGAAGGG | 8 |
| **GCCCAAGCCGACCAAAGATC** | scaffold_7 | 469951 | + | ATGATCCGAGGCGAAGTATGGACTCACTCGTTGTTCAGCCGATTTCAAGA | TAGAGCTGGTAGAACGGGACCGGGTGTGTGCTATAGATTATACACCAAAA | GCCCAAGCCGACCAAAGATCGGG | 8 |
| **TATAAACGCAGCACCCCGGC** | scaffold_7 | 532412 | + | TTGTGCAACTGTGGCCTAAACGTTACTTATGTACTCAGAGATACGTATAC | TTGATGTCCGAAATAGAAGGTTCATACTATAAGTGAGTTATCGGACTTGG | TATAAACGCAGCACCCCGGCTGG | 8 |
| **CTTTGCGTTGTGGTGGAAGC** | scaffold_7 | 562181 | + | GAGTGCCTAAAATAGATACAGACCCCGTAGATACCTGCAGAAACAGGATC | TTTTGTAGTAACTTGGACTAGAGGAAGGTGAAATATAGCTATCTGTCGAA | CTTTGCGTTGTGGTGGAAGCAGG | 6 |
| **ACCTTACTTCCTAACGGTTT** | scaffold_7 | 583079 | + | TCTTAACATCTGGGTTCTGGAAAAGCGTTTCCACTAAACCTGTTTCTAAC | TATACACTTTTATAGATCAGTACACACATAGGTATTTATGTCTACTGCAA | ACCTTACTTCCTAACGGTTTGGG | 5 |
| **CAGCGTTAGATGGCTGCTCA** | scaffold_7 | 583336 | + | CATTATTTCAAGGAAGCTAAATAGAAATGCCTTTACTAATACCCTACATA | TTATCAAAAATATTCACATCTAGTTTGCAAGTTGAAATCTAGCAGTTAAT | CAGCGTTAGATGGCTGCTCAGGG | 5 |
| **GACAGTAGTAGTGCAACCTC** | scaffold_7 | 647719 | + | CCTTGACAGATGCGTTGGCGTTGATCACTGCTATACCGAATTCTACTGGT | CATACCAAATGATGAAACAGTCACTGTTTATACATACAACTCTCAGAATT | GACAGTAGTAGTGCAACCTCGGG | 7 |
| **GGCCCAAAAGCGTTTCGTCA** | scaffold_7 | 647969 | + | TCCATCAATTTCTGCACAATTTACAAACAGCACGAAATCGGGATACAATA | ATAACACGCTCCAGTTTAAACATGGTACCTTCAGAATATCCAAAAGTCTC | GGCCCAAAAGCGTTTCGTCATGG | 7 |
| **ATCGTCAGCATGCAGGTAAC** | scaffold_7 | 658073 | + | GAGTATACATCCGTTGCGTATTATCGTGCATACACAGTTGAGCGGAAATT | GGTGTGATCAAAGTTGGGGAGGAACAACGTGAATTACGGCATCTATGGAT | ATCGTCAGCATGCAGGTAACAGG | 11 |
| **AGAGCGTTTCGGGCGATTGT** | scaffold_7 | 658189 | + | TATGGATGCGTTTCACATTGGTATTCATTCGGATATGGTGAAGCAAGAAC | CTATCGGTGTAGCGTAATTTGACGGCCAGTCATTGCAAAGTGTGAAGACG | AGAGCGTTTCGGGCGATTGTTGG | 11 |
| **GTTACACTTCGCTAGACTGT** | scaffold_7 | 658277 | + | AATTTGACGGCCAGTCATTGCAAAGTGTGAAGACGGCATCGACTAAACGA | GCTTTGCAATAGCATTTCCCCACACAAATCCAATTGTTTTGTATAGCACG | GTTACACTTCGCTAGACTGTTGG | 11 |
| **TGACTTGGGGGACAAACTGC** | scaffold_7 | 685118 | + | TGTATCAGGGAAAACAGGAATTGTAGGCAACTTCTCCAATCCGCCAGCTG | GCCTGTTGTTTTGGTGTCTCGATGATTGGTTCGTCAATCTTAAATGGATT | TGACTTGGGGGACAAACTGCTGG | 10 |
| **GACCAACCGTACCACGCTAG** | scaffold_7 | 707882 | + | CTATTTCCATCTGCATTCCAAGTGTACTGCGTTGCAATGACCGAGAAAAA | AGAAGATCATGATCATTGACTACTTTAGGAGTTCGAATCGGCCGCAAAAC | GACCAACCGTACCACGCTAGAGG | 8 |
| **TCGGCCGCAAAACGAAACCG** | scaffold_7 | 707942 | + | ACCACGCTAGAGGAGAAGATCATGATCATTGACTACTTTAGGAGTTCGAA | AGCATTTCAAGGACAAGTACTCGATTTCCACATCGAGTTTGAGTGCGTGG | TCGGCCGCAAAACGAAACCGTGG | 8 |
| **CCACATCGAGTTTGAGTGCG** | scaffold_7 | 707992 | + | TCGGCCGCAAAACGAAACCGTGGAGCATTTCAAGGACAAGTACTCGATTT | ATTAGACAGGAAGACGAACTACGGAAGAGATACAACGATTCCATGAAGGA | CCACATCGAGTTTGAGTGCGTGG | 8 |
| **AGTCCGCCCTAGGCCGTGCT** | scaffold_7 | 745219 | + | TGTTCCGTCGGAATTGTCCTGTTTCTGCTACCGGTATTCTTTCCTCGGCT | CCGCGTTTAGCACACTTCCCGCGGCCAGCCGGGGCGCGACCTCTTTCCTG | AGTCCGCCCTAGGCCGTGCTAGG | 10 |
| **GACTCGGCACCCTCTTGCCG** | scaffold_7 | 745447 | + | ATGCCGCGGCAATGTCTGCAGGACTATGACGCATCCTTTATCTCGGACTT | CACCTGTAGGTATTTTCCGCGGCAAGGGCCTCTCGAGCCGCGGAACAACC | GACTCGGCACCCTCTTGCCGCGG | 10 |
| **TGTAGGTATTTTCCGCGGCA** | scaffold_7 | 745474 | + | GACGCATCCTTTATCTCGGACTTGACTCGGCACCCTCTTGCCGCGGCACC | GCCTCTCGAGCCGCGGAACAACCAATATTCTGTCATTTCCACTCCCGTGA | TGTAGGTATTTTCCGCGGCAAGG | 10 |

| | | | | | | | |
|---|---|---|---|---|---|---|---|
| **GGCAAGGGCCTCTCGAGCCG** | scaffold_7 | 745490 | + | CGGACTTGACTCGGCACCCTCTTGCCGCGGCACCTGTAGGTATTTTCCGC | AACAACCAATATTCTGTCATTTCCACTCCCGTGAACAATAGCTGCAGATT | GGCAAGGGCCTCTCGAGCCGCGG | 10 |
| **TCGTCTTATCGCATCTCGTG** | scaffold_7 | 759781 | + | TCGGTTAGATTTAAAATTGAAAAGTTGAACGATCAATATTTCTAATAGTA | CTTGAAATAACACGGTTCCGGTTGTTCAATAAATTGAATCAAACGATTTT | TCGTCTTATCGCATCTCGTGAGG | 12 |
| **AATGCCATCACGACGTAAGC** | scaffold_7 | 857327 | + | AAATGTGATTTGTCTGCACAGTACCTGGATCTCTGTGCCCCAATGCCCAA | GTCTCTCCTCCACGAAGATAAGATTAGCCAAGTTCAGCTGTAGTGTTACA | AATGCCATCACGACGTAAGCAGG | 7 |
| **GACCTCACCCCGTCCGAAAA** | scaffold_7 | 857470 | + | ACTGGAAAAACTAAACCCGTAAGGCATTTACCCAATGTGGCAACTCACCA | AAAGTGCCCTTGCCGTACAACAAGAGGGCGACAAGAGGGAGAAAAGATAA | GACCTCACCCCGTCCGAAAAAGG | 7 |
| **TCGAGTTGACTAGGGCTGGT** | scaffold_7 | 10393 | - | TACGCGCCGGTACTGCATGGCAACTCACCATCACTGTGCCTTGCATATTT | TCTTTGGGTCTTTGTGTGAAGTTTTGTCTCTATGTTTGATGATGCGGTTG | TCGAGTTGACTAGGGCTGGTGGG | 2 |
| **AAATAGCCCGTTTATCACTG** | scaffold_7 | 46294 | - | CGATCATGCAGATGTCAGCGCCACTTGGCTATATGGTGAAGCTGACAGTA | TAGATATTGTCATATGCAATTGAATTTGGTGTCCGATTTAAGGAAAATTC | AAATAGCCCGTTTATCACTGAGG | 9 |
| **GTGTCCGAGATTAAGATACG** | scaffold_7 | 65952 | - | TTCCAAGACTGGCGTCCCAAAACTTCCTTTGAAACCTTCAGTGAGAAAAA | AGTCGGTTTCACAAGGAGGGGAAGAGGGCAAACACAGATTTCTGTTATCC | GTGTCCGAGATTAAGATACGTGG | 7 |
| **AGACCCTGCTTACGTCGTGA** | scaffold_7 | 70736 | - | CTATTGTAACACTACAGCTGAACTTGGCTAATCTTATCTTCGTGGAGGAG | CATTTTGGGCATTGGGGCACAGAGATCCAGGTACTGTGCAGACAAATCAC | AGACCCTGCTTACGTCGTGATGG | 7 |
| **CCTCCGGTTTGCTCCAGTTC** | scaffold_7 | 169927 | - | TCCAATGCTTACATCCATACATAATAAAATTGTCAGTTATTGGCATAAGG | CTAGGTGATGTGGAATACCCAGTATATCAAGGCTCTAAAAAGATACAATA | CCTCCGGTTTGCTCCAGTTCGGG | 10 |
| **ATCGTGTTTAGGTCTAAAAC** | scaffold_7 | 170266 | - | GGAATAATTACCATTAATAAAGCCAAAGCATTATCCGAAAAACAAGCAAA | ACGGCTGGCCAAACAATCCGGAAAAGTGTAGCCAGCATGACAGAACTAGC | ATCGTGTTTAGGTCTAAAACCGG | 10 |
| **GGACGGCTGGCCAAACAATC** | scaffold_7 | 170287 | - | GCCAAAGCATTATCCGAAAAACAAGCAAAATCGTGTTTAGGTCTAAAACC | AAAAGTGTAGCCAGCATGACAGAACTAGCAAGAAGTGCTAGAATAGTTAA | GGACGGCTGGCCAAACAATCCGG | 10 |
| **TGGTTGTTCCGCGGCTCGAG** | scaffold_7 | 182569 | - | ACTTATCTAATCTGCAGCTATTGTTCACGGGAGTGGAAATGACAGAATAT | CCCTTGCCGCGGAAAATACCTACAGGTGCCGCGGCAAGAGGGTGCCGAGT | TGGTTGTTCCGCGGCTCGAGAGG | 10 |
| **ACGCGGCCTAGCACGGCCTA** | scaffold_7 | 182842 | - | CTCTTTCAGGAAAGAGGTCGCGCCCCGGCTGGCCGCGGGAAGTGTGCTAA | CGGACTAGCCGAGGAAAGAATACCGGTAGCAGAAACAGGACAATTCCGAC | ACGCGGCCTAGCACGGCCTAGGG | 10 |
| **CGGCCTAGCACGGCCTAGGG** | scaffold_7 | 182845 | - | TTTCAGGAAAGAGGTCGCGCCCCGGCTGGCCGCGGGAAGTGTGCTAAACG | ACTAGCCGAGGAAAGAATACCGGTAGCAGAAACAGGACAATTCCGACGGA | CGGCCTAGCACGGCCTAGGGCGG | 10 |
| **GGCCTAGGGCGGACTAGCCG** | scaffold_7 | 182856 | - | AGGTCGCGCCCCGGCTGGCCGCGGGAAGTGTGCTAAACGCGGCCTAGCAC | AAAGAATACCGGTAGCAGAAACAGGACAATTCCGACGGAACAAAACACGG | GGCCTAGGGCGGACTAGCCGAGG | 10 |
| **CCACGCACTCAAACTCGATG** | scaffold_7 | 220075 | - | TCCTTCATGGAATCGTTGTATCTCTTCCGTAGTTCGTCTTCCTGTCTAAT | AAATCGAGTACTTGTCCTTGAAATGCTCCACGGTTTCGTTTTGCGGCCGA | CCACGCACTCAAACTCGATGTGG | 8 |
| **GACCTCTCACCATCGAGTCG** | scaffold_7 | 277358 | - | TTCTTCGTTACAGACACATAGATATGGGAAGGGTTTGCGTAGACTACTTT | GGACATTACGTCGGGGTCCAGTCCCAAGTCCAACGACAGGGCCAATATTA | GACCTCTCACCATCGAGTCGGGG | 8 |
| **ACCTCTCACCATCGAGTCGG** | scaffold_7 | 277359 | - | TCTTCGTTACAGACACATAGATATGGGAAGGGTTTGCGTAGACTACTTTG | GACATTACGTCGGGGTCCAGTCCCAAGTCCAACGACAGGGCCAATATTAC | ACCTCTCACCATCGAGTCGGGGG | 8 |
| **GTGGCGAGTCTAACACACGC** | scaffold_7 | 277491 | - | CAAGTATAGGCGGCCAAGTGCAACGAGATCATCAATCTCGTCGATTTATA | TCTTCCAGTCATTCACAGTACTCATTTGGGTCTGTCTTTTCTGGGGTCAT | GTGGCGAGTCTAACACACGCAGG | 8 |
| **ACACTCATGGTTCGGTCGAT** | scaffold_7 | 279019 | - | CACTTTCAAATTCATCCCAGATACATTGCTCGTTTGGAGATTCATACAAC | TAGTCTGTCTATCGGGGTGGTTGGGGTATCATCAACTTGGTAGTTCATTT | ACACTCATGGTTCGGTCGATTGG | 7 |
| **GATTGGTAGTCTGTCTATCG** | scaffold_7 | 279036 | - | CAGATACATTGCTCGTTTGGAGATTCATACAACACACTCATGGTTCGGTC | TGGTTGGGGTATCATCAACTTGGTAGTTCATTTTATCGATAGAATTTATT | GATTGGTAGTCTGTCTATCGGGG | 7 |
| **GAAGCAGGAAGTAATCCCGT** | scaffold_7 | 280222 | - | TTGTAGAATTTAATGGCTGAGTAAAAATTGATGAATTAGTATAGTTTGAC | AAGGGTATTCGTAGTGATTGTTGGAACTATCAGCTCAATTGAAGGTTTTA | GAAGCAGGAAGTAATCCCGTCGG | 7 |
| **CAGGAAGTAATCCCGTCGGA** | scaffold_7 | 280226 | - | AGAATTTAATGGCTGAGTAAAAATTGATGAATTAGTATAGTTTGACGAAG | GTATTCGTAGTGATTGTTGGAACTATCAGCTCAATTGAAGGTTTTAATGA | CAGGAAGTAATCCCGTCGGAAGG | 7 |
| **CACGAAAATTTCCCGCGTAC** | scaffold_7 | 287781 | - | CACTTCCTAGTTAGACAATTGCAAGTTTGCCGTTCCATATATTCGCTTTG | ATTGCCGCCAAAACTCGTTTGTGTTAATCCCAAATGCTCATGAAATAGGG | CACGAAAATTTCCCGCGTACTGG | 8 |
| **ATAAGGCCTGGGTATACATC** | scaffold_7 | 361274 | - | TGCCACTCACTAGGTAGACATGCTACTTTTCTCTCTTGGTTTTACCACCT | GTGTATGTAGATTAAATTGTCTACATATAAAATGATATGGAAACCAATTC | ATAAGGCCTGGGTATACATCGGG | 5 |
| **TAAGGCCTGGGTATACATCG** | scaffold_7 | 361275 | - | GCCACTCACTAGGTAGACATGCTACTTTTCTCTCTTGGTTTTACCACCTA | TGTATGTAGATTAAATTGTCTACATATAAAATGATATGGAAACCAATTCA | TAAGGCCTGGGTATACATCGGGG | 5 |
| **GCGCCTCGCTGAGAAAAGTG** | scaffold_7 | 366022 | - | TTGTCTTCACAATGTGCATATTGGGTTGATCCTTTAGAAAGAATAAAGTT | CGCCATTGTTTGCTTTTCCCCGCCGTGACTTTCAGCCTCGTTTCGGTTTG | GCGCCTCGCTGAGAAAAGTGTGG | 6 |
| **AGGGGATGCACTGTGAGGTT** | scaffold_7 | 366179 | - | AATTTGTCAAACACGTGACACTCGGTGGAATCATTCCATTGTTCTTTAAA | GAAGTAATTGTGAGCAGAATGTTCTTATCTTTTCTTATCGGAGAAAGGTG | AGGGGATGCACTGTGAGGTTGGG | 6 |
| **GTCGCTGTCACGGGATCACG** | scaffold_7 | 373169 | - | GGGTTTTCGTGAAGTGCAAGTGGAGTGTGGAAGTTGAGGCGGACACGTGC | TGCGCCACTGTTGGCATGGCTGCCTGTGGGACTCTTGTTTGTTGCGAGGG | GTCGCTGTCACGGGATCACGTGG | 7 |
| **TGCGTGTCCCATTTCCGGAT** | scaffold_7 | 406422 | - | AGATTTTGATGTGCTTTCAAATGGTACCGTTAATGATAGTGAAACTATTG | CAAATGTTTGCCGAGTGTATATCAGCAGGCGACCTCTGCGATCTAGATGG | TGCGTGTCCCATTTCCGGATAGG | 8 |
| **AGAGTTCTGCAGTTACGCTA** | scaffold_7 | 453543 | - | TGAATACGGTCCCCCTCTTTTGCCAATGACCTTGACAGAAAAACCAAAGG | AGAATGTACTCCAAGGCCAAGGCAGGTCTTCCGTACAAATCAAGAATGGA | AGAGTTCTGCAGTTACGCTATGG | 7 |
| **AGCCGCCTTCAGCCAAATAC** | scaffold_7 | 458975 | - | GGCGGCAACCCTTCTAGGCTGTGTACAAGCTATGATGCCATCATTAGCAT | GTGATTTGTGTAGTTTTACCAGAACCTGTTTCACCGACAATAACTACAAA | AGCCGCCTTCAGCCAAATACTGG | 8 |
| **GCACTTAGTGCGGGCTTAGC** | scaffold_7 | 492603 | - | GCCCTTAGGGCGCGTACAGTTTCCAACGCCTTTTGTTGAGAAGCCAACGG | ATGCTCAGCACAGTCTGCCAACACATTCTCTCAACCAACTGTTTTCCAAT | GCACTTAGTGCGGGCTTAGCTGG | 5 |
| **ACGTGATCGACGGCCAGGTT** | scaffold_7 | 502780 | - | TTATCAGAGTCTAGATCCAAATGCTGTAACGTTGGATTTAAAAATTTCTT | TGACTTTTGAATTTCCAACATTTTCTATAATTGTATTATGTTTACTTTCA | ACGTGATCGACGGCCAGGTTTGG | 6 |
| **GAACCTCGTCCGTGACATTG** | scaffold_7 | 503543 | - | GCGGGGTTGGAATGACAGGAACAAATTGCGAGAAGCTATTACTCATTGGT | CTTAGACAATGTGTTTGAAGGGTCAATAGCTGACTGTAATCCACTTAGTA | GAACCTCGTCCGTGACATTGAGG | 6 |
| **ATGATTACATGCCCTCTCGG** | scaffold_7 | 535449 | - | CCCCTCGGTTCGCCATCTCTCCCTTGTATATCTTGTCGACGAAACACACA | AGTCCCCTCCCGTTGCTTTTCGAGATAAACTCAACCCAGATCAATTTCTA | ATGATTACATGCCCTCTCGGGGG | 5 |
| **CAACGACCTAAACTCGTCCC** | scaffold_7 | 535564 | - | AATTTCTACAACCTCAAACCTCAATATGCAAGAATCGTGGCATACCTCTT | TGGATCCAAATTTAAATGTAAAGTCCTCAAAACCTTACTCTAAATTGAAC | CAACGACCTAAACTCGTCCCTGG | 5 |
| **CCCCTATACCATAATACATA** | scaffold_19 | 218981 | + | TATTAGATGAAGGGGAATGAGCCTCCAGGATAGTAATGTTACACCAGAAC | AGAATGATTCTCATCCAGTTTGAAAAACAACAGCAACAATAGTAACAACA | CCCCTATACCATAATACATACGG | 0 |
| **CCGTTATAGATGGTAGGTGT** | scaffold_19 | 224847 | + | TTTTATAAAAGACGTCCAAGTAGCTGCACTCATGTCTGTGGATGTGTAGG | TATATGGGGACGAGAACAGAAACGAGGATGCACGGAGGGGAGGTTTGGCT | CCGTTATAGATGGTAGGTGTAGG | 0 |
| **TCATGCTATGCTGTGGACCG** | scaffold_19 | 255703 | + | ATTTATTAACATGTATAGATTTTCATGCAGTCTTAAAGCTTACTGCATTG | GGTGCCTTCTCATCACTCTTCAATGCAGATGCATCTTCAACACGATTATC | TCATGCTATGCTGTGGACCGAGG | 1 |
| **ATCCTCTTCTTAGGTGCACA** | scaffold_19 | 264887 | + | TAATCTCCCTGTGGAACGTTATGGCAATTGTATCACCGTTATTCGTAAGT | AATCACTGTCTTATCCTTGCAAGCAAAAACAACGTATTTAGAGAGATTCT | ATCCTCTTCTTAGGTGCACAAGG | 0 |
| **TCGTAACGAGTCGCTTGGCG** | scaffold_19 | 94910 | - | ATTGGTGTATTACGGTGGTTTATATCCACCTTGAAATCACAATATTGTTC | AGAAAAGAAACCATTGAACCCCTTCCTTGGAGAAATCTTTGTTGGTAAAT | TCGTAACGAGTCGCTTGGCGGGG | 0 |
| **ATCGTATAAACGCTGATCCG** | scaffold_19 | 101492 | - | TTCGATTGAAACTATATCATTATTAATAGTGCTTAAGTTGAAATACCCAA | GAACCATGAATCGAGGCAAATTACAACAAACTATGGATTCTATACAGAAT | ATCGTATAAACGCTGATCCGAGG | 0 |
| **GGTGCAGTTTGCGTCTGTAA** | scaffold_19 | 122974 | - | TTCCTGGCGGTGGAGGCGGCGGTGGTGCAGCAGAGGAGCCAGCAGCGGGT | AGGTGGTGGTGGCGGTGGAGGAGCCAGAGATTGTTGCTGTTGATGTGGCG | GGTGCAGTTTGCGTCTGTAAAGG | 0 |
| **TTGGACCCGGGTGGCCGTTC** | scaffold_19 | 151895 | - | GTCGATCAATTCCACAACAATAGGTTCCCTCTCCTTGGGAGACACCGTTG | TGGGCCATTATGTCCTTGCGTTTCAATTTGTCAATCTGCTTCCTTTCCAG | TTGGACCCGGGTGGCCGTTCGGG | 0 |
| **GAGGGGACCAATTTCGTGTT** | scaffold_19 | 188875 | - | CCGTAATACGCCCTTTCCGGGCTTATTATATTAAGACGCGTATTTATATG | TGAAATGTAGATGCCAAACTTCAAGTATAGGCCAACGAGGTACACCACCT | GAGGGGACCAATTTCGTGTTTGG | 0 |
| **CTTCAGTCGGCTGTCTAGTG** | scaffold_19 | 271131 | - | ACAATCTCTTTCCATCTCTCCCAATCTTAAACCACCATCTCTAGACCTAC | ACGGCTCTAGGACCTCGTGCTCTAGCATGCATTTTATCCAAAACCATATG | CTTCAGTCGGCTGTCTAGTGAGG | 0 |
| **ACGTGAAAAGTCCCTCCCCC** | scaffold_19 | 278688 | - | TTTCAGCTATTGAAGAAATGTTTTATTTCTCAAAATGCTGAAATTTTGAA | TGAATGGGGAACTCGCTGACGTGTTGTTGAGCGAAGTGCCAATTTAAGAA | ACGTGAAAAGTCCCTCCCCCCGG | 0 |
| **ACGGTCGACAACTCCACGAG** | scaffold_20 | 253322 | + | ATACAATGGCATTCCAGACCATCTAAAGAAATATGCTAGCTTGGAATAGG | TCTCCAATTGGCGCCGGCTCTGTAATGATGTATGTAATGTTTCAGATTAA | ACGGTCGACAACTCCACGAGTGG | 0 |
| **TGAGCATTCTTAAGCGTGAA** | scaffold_20 | 191688 | - | CGAAAAACACCTCCTAAATCTAAAAGAATGCTTAATTGATTGGTTATATC | TATTTCGATACCCAAAAGTGACAATACAACACCAGGAGAAAGAGAGATAA | TGAGCATTCTTAAGCGTGAATGG | 0 |
| **TTCACCGAGTATTCGGTATG** | scaffold_20 | 284828 | - | TCCTTTAAACTCTAATTTTATTGAGTGTAATAATAGTGTGTTTGTGTAAC | TTGTCTGTCTTGTTCGACGTTTTATAAGAAGGAAAGACCCTACTCTTTAC | TTCACCGAGTATTCGGTATGAGG | 0 |
| **CTAACCTCAAGTGAGGGGAA** | scaffold_24 | 170 | + | TGTTTGTTCTCGAGTAAATGAGAAACCCAAAAGGTAACAACTTAGAGTAA | GTCAATATAAATAGTGCAAAATAGCTGCAAACCCATGACAGTAAACAGTG | CTAACCTCAAGTGAGGGGAATGG | 4 |
| **TACGACCTGAATCGTCCGTT** | scaffold_24 | 182120 | - | TCTGCCTGTACTTGCCAGCATATATATGACCGACCAATCTAAGATCCCTC | TTATCCTTCAATGAGCGCAATTTAAACCAATATGCGACCCCCACTAATAA | TACGACCTGAATCGTCCGTTAGG | 8 |
| **CCCATTATGGTCAGCGGCAA** | scaffold_26 | 10724 | + | CTATAACGAGTGGTGTATATCAATGGTTAATGTTTTCAGATCTCTCCCCC | AAGGTCGATTTGGGAAACAAGAGACGCAATCAATACAGACACCGAGAATA | CCCATTATGGTCAGCGGCAAGGG | 0 |
| **TCGGATAGACTGCGCAAGCC** | scaffold_26 | 43376 | + | GAACCCGGGTTTCCCGCGTGACAAGCGGAAATTCTAGCCACTAAACTATC | AATCGAACCAGGGGCTCAACGATGGCAACGTTGAATTTTACCACTAAACC | TCGGATAGACTGCGCAAGCCCGG | 0 |
| **AGCAGATGGGTGGCGAGTGT** | scaffold_26 | 79418 | + | GCCATTGTATGTCTACATGTAGTATATACCACTAGGAGACACTGCGTTCC | ATGCGTCTCTTCCAACTGCATAATAGGACACTCATTTGACCCTCTCAACA | AGCAGATGGGTGGCGAGTGTCGG | 0 |
| **TGGCACATACGTCGACTCAT** | scaffold_26 | 83458 | + | TGCTACATCCACAACTACACGTACGTTGGCACGTACGTTGGCACATACGT | TCGACTCATATAGCCACTCCTACGGCCAAGTCTATAGTTACATCCACACT | TGGCACATACGTCGACTCATAGG | 0 |
| **CGAGTCTTACTTTAGTACAC** | scaffold_26 | 161501 | + | ACAGATGACAGTGAGAAATCACCAACTGAATTGATAACACAGTATCTACC | AATCTGCATTGAATGCCATAAAGCAGATGTTCTCACAGCAGTTGGTTTAT | CGAGTCTTACTTTAGTACACCGG | 0 |
| **AACTACTGTCGGTGGTCCGG** | scaffold_26 | 3423 | - | ACTGTTTACCAGAATGCGGACGGTAATGCAGTCACCTCTTCTGTCGAACA | TGCAACTACTTTAGAAACTCAAATGGTAGCAACCCCCGCTGCTTCTAACG | AACTACTGTCGGTGGTCCGGTGG | 1 |
| **AGCCTAATAGCCTTTTGGGA** | scaffold_26 | 88841 | - | TGTGGATTTCTGTAAACTCTAGTATAGTTTTCTTTGGATATTGTCTCTTT | GCCTTTTCTCAGTTTTATACATATAATACACACACACACATATATATATA | AGCCTAATAGCCTTTTGGGAAGG | 0 |
| **CCCCGAAGTCGACAAATTTA** | scaffold_26 | 192446 | - | CCATCATCATCATCATCATCATCAGAATCCGAAATTACCGGAGTATTAGT | ATTTCAACTGTTGGAGGGTCAATCCATGAGAGAATTTAGAATCTTTAGTG | CCCCGAAGTCGACAAATTTAAGG | 0 |
| **GCATAGAAATCGGTCCAAGC** | scaffold_29 | 14191 | - | CCAGACTAGAAAGGCCCCGGATTTCCATTACTGGATAGAAGGGCTATGAG | ACCATTGGAAGAGAAATTTGCTGCTACATATGGGCTACCAGATGTCAGAA | GCATAGAAATCGGTCCAAGCAGG | 2 |
| **CTCAGGCTGCCTTTAAATGA** | scaffold_32 | 126470 | + | TTGATCTCGTTGCATATCTTAGGTCCCGAGTTATAGAGAAAGGTCTTGAA | CTGTTGATATGTATTTCAAAAATATAGCAGTACCGAAGTTAATAACTCCA | CTCAGGCTGCCTTTAAATGAAGG | 5 |
| **CTCGGCCACTTATGCTGTTA** | scaffold_32 | 130683 | + | ACCTGCGCACTCTTACGAGTAACAGAACTTGAGTCTGTCGCCTTAGACCG | ACTTGTAAGAGGTCAAATGAATCACTATCTCTATATTTCTTAACCGGGTA | CTCGGCCACTTATGCTGTTACGG | 3 |
| **GGTGGTACGGGCCAGATGGA** | scaffold_32 | 4138 | - | AAGGAACAGAAGCACGAGAAGTTCCTGGTGTTGAATAATAGTTAGAACTG | AATTGAATGAAAAGAAGGTCCCTAAATAATCAATAGTAACGAAGAAATTA | GGTGGTACGGGCCAGATGGAAGG | 4 |
| **TGAAGAGCCTATCCGATGGG** | scaffold_32 | 6685 | - | TTGCACATTCACTAAAGCAAATTTGGATTGTTCGGTTACTCATATTTAGG | ACCACTGATATGTAAAACTAATGTTAGATTCAGCTGAATGGTTTGTCATT | TGAAGAGCCTATCCGATGGGTGG | 5 |
| **CCCACCTGAGATCTAGACGG** | scaffold_35 | 80195 | + | AAGTTGACGAGTTCCAAGATTCCAGTCAAAATACTGCCACCGCATAATGG | ATGGCGGTTATGTAGCAACCAAAAGTGTGTGGTGATATCGCCTTCCAAAC | CCCACCTGAGATCTAGACGGGGG | 1 |
| **GTTGATAAAGCCAGGGTCCC** | scaffold_35 | 80658 | + | TGAGAGCGGCCAGTCGGGCAGCCGGCGGGGCGGTTACTCATGCAACTTCG | AGGGAGCGGAGGGGCGTTTTCAGGACGCTGCTCAGCCAGCCGGGGGACG/ | GTTGATAAAGCCAGGGTCCCCGG | 1 |
| **GAGACGCTGTGACGTGTGGC** | scaffold_35 | 80862 | + | AGCCAGCGGGAAACTAGCCGAGGGGGAACGGGGGCTGCGCGTAGCCGGC( | ACGCTCTCCACAGGACGCGGTGCCGGGTGCTGCGTGCGAGACGTGCGGA( | GAGACGCTGTGACGTGTGGCCGG | 1 |
| **CTCTCCACAGGACGCGGTGC** | scaffold_35 | 80888 | + | AACGGGGGCTGCGCGTAGCCGGCGGAGACGCTGTGACGTGTGGCCGGACG | GTGCTGCGTGCGAGACGTGCGGAGCGTGCGGGCCGTGGAATGAAAGTCGG | CTCTCCACAGGACGCGGTGCCGG | 1 |
| **TCTCCACAGGACGCGGTGCC** | scaffold_35 | 80889 | + | ACGGGGGCTGCGCGTAGCCGGCGGAGACGCTGTGACGTGTGGCCGGACG( | TGCTGCGTGCGAGACGTGCGGAGCGTGCGGGCCGTGGAATGAAAGTCGGA | TCTCCACAGGACGCGGTGCCGGG | 1 |
| **GGGTGAGCGGGGGAACGGCT** | scaffold_35 | 86822 | + | AGGACTATGTTGTGTGGAGGGGGTGGGGCTGTAGAGAGGATGGATTTGGA | AGGCCGCCCATGTCGTATGCCCCGGGGTAGGGTATAGGCCGGGAGGGGCC | GGGTGAGCGGGGGAACGGCTTGG | 1 |
| **CACGTCTCGCACGCAGCACC** | scaffold_35 | 9049 | - | CTTCCACATAGCAACCTTTCCGACTTTCATTCCACGGCCCGCACGCTCCG | CACCGCGTCCTGTGGAGAGCGTCCGGCCACACGTCACAGCGTCTCCGCCG | CACGTCTCGCACGCAGCACCCGG | 1 |
| **GCACCCGGCACCGCGTCCTG** | scaffold_35 | 9064 | - | CTTTCCGACTTTCATTCCACGGCCCGCACGCTCCGCACGTCTCGCACGCA | AGAGCGTCCGGCCACACGTCACAGCGTCTCCGCCGGCTACGCGCAGCCC( | GCACCCGGCACCGCGTCCTGTGG | 1 |
| **CGCGTCCTGTGGAGAGCGTC** | scaffold_35 | 9075 | - | TCATTCCACGGCCCGCACGCTCCGCACGTCTCGCACGCAGCACCCGGCAC | CCACACGTCACAGCGTCTCCGCCGGCTACGCGCAGCCCCCGTTCCCCCTC | CGCGTCCTGTGGAGAGCGTCCGG | 1 |
| **AAACGCCCCTCCGCTCCCTC** | scaffold_35 | 9279 | - | GTGCAAGCCCAAAGGCTCCTCGTCCCCCGGCTGGCTGAGCAGCGTCCTGA | GGACCCTGGCTTTATCAACCGAAGTTGCATGAGTAACCGCCCCGCCGGCT | AAACGCCCCTCCGCTCCCTCCGG | 1 |
| **TCCGCTCCCTCCGGGGACCC** | scaffold_35 | 9288 | - | CAAAGGCTCCTCGTCCCCCGGCTGGCTGAGCAGCGTCCTGAAAACGCCCC | CTTTATCAACCGAAGTTGCATGAGTAACCGCCCCGCCGGCTGCCCGACTG | TCCGCTCCCTCCGGGGACCCTGG | 1 |
| **CATCAAAAGCCGGCGGGTTG** | scaffold_39 | 35480 | - | TGTTGCCTCCTCTGGAAAGTATCCACTGCATTACCAACGGGTATTCCACA | TACTTTTCAACAAACCCCGGAGATGGGAATAGTTTGCAAGAGACGACATG | CATCAAAAGCCGGCGGGTTGAGG | 1 |

| genes_in_vicinity | intergenic_region_size | 3'_gene_dist | 3'_T_or_P | 3'_gene | 5'_gene_dist | 5'_T_or_P | 5'_gene | adjacent_gene_ess | avg_expr_vic_2 | on_target_activity (Rule Set 2 scores) |
|---|---|---|---|---|---|---|---|---|---|---|
| mRNA_2, mRNA_1 | - | 1398 | T | mRNA_1 | - | - | - | NA | NA | 0.4429 |
| mRNA_18, mRNA_15, mRNA_12, mRNA_17, mRNA_13, mRNA_14, mRNA_19, mRNA_11, mRNA_16 | 1066 | 191 | T | mRNA_15 | 875 | P | mRNA_14 | NA | NA | 0.5116 |
| mRNA_40, mRNA_38, mRNA_32, mRNA_37, mRNA_39, mRNA_36, mRNA_35, mRNA_34, mRNA_33 | 1945 | 1329 | T | mRNA_37 | 616 | T | mRNA_36 | NA | 0.2 | 0.5105 |
| mRNA_114, mRNA_113, mRNA_108, mRNA_111, mRNA_110, mRNA_106, mRNA_105, mRNA_109, mRNA_112, mRNA_107 | 657 | 420 | T | mRNA_110 | 237 | T | mRNA_109 | 1 | NA | 0.3406 |
| mRNA_139, mRNA_141, mRNA_142, mRNA_140, mRNA_144, mRNA_145, mRNA_143 | 1443 | 991 | P | mRNA_142 | 452 | T | mRNA_141 | NA | NA | 0.537 |
| mRNA_664, mRNA_663, mRNA_661, mRNA_660, mRNA_662 | 2057 | 1704 | P | mRNA_663 | 353 | T | mRNA_662 | NA | NA | 0.5437 |
| mRNA_664, mRNA_663, mRNA_661, mRNA_660, mRNA_662 | 2057 | 1114 | P | mRNA_663 | 943 | T | mRNA_662 | NA | NA | 0.2659 |
| mRNA_1047, mRNA_1045, mRNA_1046, mRNA_1048, mRNA_1049, mRNA_1044 | 1313 | 344 | T | mRNA_1045 | 969 | P | mRNA_1044 | NA | NA | 0.3872 |
| mRNA_1210, mRNA_1212, mRNA_1211, mRNA_1209, mRNA_1207, mRNA_1208 | 3122 | 1726 | P | mRNA_1211 | 1396 | P | mRNA_1210 | NA | NA | 0.4687 |
| mRNA_1213, mRNA_1215, mRNA_1214, mRNA_1217, mRNA_1216, mRNA_1212 | - | - | - | - | 490 | P | mRNA_1217 | NA | 0.89 | 0.6291 |
| mRNA_1047, mRNA_1045, mRNA_1046, mRNA_1048, mRNA_1049, mRNA_1044 | 1313 | 987 | P | mRNA_1044 | 326 | T | mRNA_1045 | NA | NA | 0.3781 |
| mRNA_655, mRNA_654, mRNA_652, mRNA_653, mRNA_656, mRNA_657 | 1341 | 797 | T | mRNA_653 | 544 | T | mRNA_654 | NA | NA | 0.5483 |
| mRNA_54, mRNA_53, mRNA_51, mRNA_52, mRNA_49 | 6463 | 1780 | P | mRNA_52 | 4683 | T | mRNA_53 | NA | NA | 0.6979 |
| mRNA_54, mRNA_53, mRNA_51, mRNA_52, mRNA_49 | 6463 | 1445 | P | mRNA_52 | 5018 | T | mRNA_53 | NA | NA | 0.5992 |
| mRNA_40, mRNA_38, mRNA_37, mRNA_39, mRNA_36, mRNA_35, mRNA_41, mRNA_34, mRNA_33 | 1945 | 1481 | T | mRNA_36 | 464 | T | mRNA_37 | NA | 0.2 | 0.4976 |
| mRNA_40, mRNA_38, mRNA_37, mRNA_39, mRNA_36, mRNA_35, mRNA_41, mRNA_34, mRNA_33 | 1945 | 1447 | T | mRNA_36 | 498 | T | mRNA_37 | NA | 0.2 | 0.7047 |
| mRNA_40, mRNA_38, mRNA_32, mRNA_37, mRNA_39, mRNA_36, mRNA_35, mRNA_34, mRNA_33 | 1945 | 572 | T | mRNA_36 | 1373 | T | mRNA_37 | NA | 0.2 | 0.5001 |
| mRNA_2, mRNA_1 | - | - | - | - | 1423 | T | mRNA_1 | NA | NA | 0.3369 |
| mRNA_1220, mRNA_1223, mRNA_1222, mRNA_1221, mRNA_1219 | - | 364 | T | mRNA_1219 | - | - | - | NA | NA | 0.5163 |
| mRNA_1582, mRNA_1584, mRNA_1583, mRNA_1580, mRNA_1579, mRNA_1578, mRNA_1577, mRNA_1576 | 1745 | 245 | T | mRNA_1580 | 1500 | P | mRNA_1579 | 1 | NA | 0.5943 |
| mRNA_2145, mRNA_2144, mRNA_2147, mRNA_2146, mRNA_2141, mRNA_2148, mRNA_2142, mRNA_2143 | 2317 | 1552 | P | mRNA_2145 | 765 | P | mRNA_2144 | NA | NA | 0.5653 |
| mRNA_2151, mRNA_2153, mRNA_2154, mRNA_2149, mRNA_2150, mRNA_2152 | 2682 | 1639 | P | mRNA_2154 | 1043 | P | mRNA_2153 | 1 | NA | 0.4155 |
| mRNA_2151, mRNA_2153, mRNA_2154, mRNA_2149, mRNA_2150, mRNA_2152 | 2682 | 1570 | P | mRNA_2154 | 1112 | P | mRNA_2153 | 1 | NA | 0.6234 |
| mRNA_2151, mRNA_2153, mRNA_2154, mRNA_2149, mRNA_2150, mRNA_2152 | 2682 | 1388 | P | mRNA_2154 | 1294 | P | mRNA_2153 | 1 | NA | 0.5701 |
| mRNA_2346, mRNA_2349, mRNA_2348, mRNA_2345, mRNA_2347 | 3523 | 1454 | P | mRNA_2348 | 2069 | P | mRNA_2347 | NA | NA | 0.3033 |
| mRNA_2155, mRNA_2151, mRNA_2153, mRNA_2154, mRNA_2149, mRNA_2150, mRNA_2152 | 2682 | 1359 | P | mRNA_2153 | 1323 | P | mRNA_2154 | 1 | NA | 0.5492 |
| mRNA_1961, mRNA_1963, mRNA_1958, mRNA_1965, mRNA_1960, mRNA_1964, mRNA_1962, mRNA_1959 | 765 | 466 | T | mRNA_1962 | 299 | T | mRNA_1963 | NA | NA | 0.6501 |
| mRNA_1232, mRNA_1230, mRNA_1228, mRNA_1233, mRNA_1231, mRNA_1227, mRNA_1234, mRNA_1229, mRNA_1225, mRNA_1226 | 3122 | 2077 | T | mRNA_1228 | 1045 | T | mRNA_1229 | NA | NA | 0.6153 |
| mRNA_1232, mRNA_1230, mRNA_1228, mRNA_1231, mRNA_1227, mRNA_1229, mRNA_1225, mRNA_1226 | 3122 | 1183 | T | mRNA_1228 | 1939 | T | mRNA_1229 | NA | NA | 0.4202 |
| mRNA_1232, mRNA_1230, mRNA_1228, mRNA_1231, mRNA_1227, mRNA_1229, mRNA_1225, mRNA_1226 | 3122 | 1079 | T | mRNA_1228 | 2043 | T | mRNA_1229 | NA | NA | 0.5294 |
| mRNA_3183, mRNA_3182, mRNA_3180, mRNA_3181, mRNA_3184 | 2450 | 1386 | T | mRNA_3182 | 1064 | P | mRNA_3181 | 1 | 101.77 | 0.5915 |
| mRNA_2973, mRNA_2972, mRNA_2970, mRNA_2968, mRNA_2969, mRNA_2971, mRNA_2975, mRNA_2974 | 2362 | 126 | T | mRNA_2972 | 2236 | P | mRNA_2973 | 1 | NA | 0.525 |
| mRNA_2406, mRNA_2401, mRNA_2402, mRNA_2405, mRNA_2403, mRNA_2407, mRNA_2404, mRNA_2408, mRNA_2409 | 2135 | 1320 | P | mRNA_2404 | 815 | P | mRNA_2405 | NA | NA | 0.5244 |
| mRNA_3748, mRNA_3747, mRNA_3749, mRNA_3746, mRNA_3745 | 39112 | 760 | P | mRNA_3745 | 38352 | P | mRNA_3744 | NA | NA | 0.6329 |
| mRNA_3748, mRNA_3747, mRNA_3749, mRNA_3746, mRNA_3745 | 39112 | 759 | P | mRNA_3745 | 38353 | P | mRNA_3744 | NA | NA | 0.5569 |
| mRNA_3753, mRNA_3756, mRNA_3749, mRNA_3754, mRNA_3758, mRNA_3751, mRNA_3755, mRNA_3757, mRNA_3750 | 1081 | 522 | T | mRNA_3754 | 559 | T | mRNA_3753 | NA | NA | 0.673 |
| mRNA_3753, mRNA_3756, mRNA_3749, mRNA_3754, mRNA_3758, mRNA_3751, mRNA_3755, mRNA_3757, mRNA_3750 | 1081 | 505 | T | mRNA_3754 | 576 | T | mRNA_3753 | NA | NA | 0.6027 |
| mRNA_3895, mRNA_3894, mRNA_3893, mRNA_3896, mRNA_3892, mRNA_3890, mRNA_3891, mRNA_3897 | 3012 | 1021 | P | mRNA_3894 | 1991 | P | mRNA_3893 | 1 | NA | 0.5611 |
| mRNA_3895, mRNA_3894, mRNA_3893, mRNA_3896, mRNA_3892, mRNA_3890, mRNA_3891, mRNA_3897 | 3012 | 919 | P | mRNA_3894 | 2093 | P | mRNA_3893 | 1 | NA | 0.742 |
| mRNA_4111, mRNA_4109, mRNA_4108, mRNA_4106, mRNA_4107, mRNA_4110 | 2287 | 1288 | T | mRNA_4111 | 999 | P | mRNA_4110 | NA | NA | 0.665 |
| mRNA_4133, mRNA_4131, mRNA_4134, mRNA_4130, mRNA_4132, mRNA_4125, mRNA_4126, mRNA_4129, mRNA_4128, mRNA_4127 | 1444 | 212 | T | mRNA_4131 | 1232 | P | mRNA_4130 | 1 | NA | 0.5763 |
| mRNA_4133, mRNA_4131, mRNA_4134, mRNA_4130, mRNA_4132, mRNA_4135, mRNA_4129, mRNA_4128, mRNA_4127 | 1831 | 831 | P | mRNA_4132 | 1000 | P | mRNA_4131 | NA | NA | 0.3345 |
| mRNA_4136, mRNA_4141, mRNA_4138, mRNA_4134, mRNA_4137, mRNA_4139, mRNA_4135, mRNA_4140 | 2704 | 2164 | P | mRNA_4139 | 540 | P | mRNA_4138 | NA | NA | 0.5639 |
| mRNA_4136, mRNA_4141, mRNA_4142, mRNA_4138, mRNA_4137, mRNA_4139, mRNA_4135, mRNA_4140 | 2704 | 1726 | P | mRNA_4139 | 978 | P | mRNA_4138 | NA | NA | 0.5188 |
| mRNA_4154, mRNA_4152, mRNA_4151, mRNA_4150, mRNA_4149, mRNA_4148, mRNA_4145, mRNA_4146, mRNA_4147, mRNA_4153, mRNA_4144 | 795 | 341 | T | mRNA_4150 | 454 | P | mRNA_4149 | 1 | NA | 0.6489 |
| mRNA_4155, mRNA_4154, mRNA_4152, mRNA_4151, mRNA_4150, mRNA_4157, mRNA_4149, mRNA_4158, mRNA_4156, mRNA_4148, mRNA_4153 | 1704 | 747 | T | mRNA_4153 | 957 | T | mRNA_4152 | 1 | NA | 0.1408 |
| mRNA_4155, mRNA_4154, mRNA_4152, mRNA_4151, mRNA_4150, mRNA_4157, mRNA_4149, mRNA_4158, mRNA_4156, mRNA_4153 | 1704 | 359 | T | mRNA_4153 | 1345 | T | mRNA_4152 | 1 | NA | 0.6269 |
| mRNA_4159, mRNA_4155, mRNA_4154, mRNA_4152, mRNA_4151, mRNA_4157, mRNA_4158, mRNA_4160, mRNA_4156, mRNA_4153 | 2198 | 1689 | T | mRNA_4156 | 509 | P | mRNA_4155 | NA | NA | 0.5877 |
| mRNA_4184, mRNA_4183, mRNA_4187, mRNA_4188, mRNA_4186, mRNA_4185 | 1694 | 991 | P | mRNA_4187 | 703 | P | mRNA_4186 | NA | NA | 0.6163 |
| mRNA_4217, mRNA_4216, mRNA_4221, mRNA_4220, mRNA_4223, mRNA_4222, mRNA_4218, mRNA_4219 | 1352 | 522 | T | mRNA_4221 | 830 | P | mRNA_4220 | 1 | NA | 0.5878 |
| mRNA_4221, mRNA_4224, mRNA_4220, mRNA_4223, mRNA_4222, mRNA_4218, mRNA_4219, mRNA_4225 | 1994 | 974 | P | mRNA_4222 | 1020 | P | mRNA_4221 | 1 | NA | 0.5405 |
| mRNA_4242, mRNA_4236, mRNA_4235, mRNA_4240, mRNA_4239, mRNA_4241, mRNA_4243, mRNA_4237, mRNA_4238 | 544 | 289 | T | mRNA_4240 | 255 | T | mRNA_4239 | NA | NA | 0.261 |
| mRNA_4252, mRNA_4250, mRNA_4251 | 8607 | 4549 | P | mRNA_4252 | 4058 | T | mRNA_4251 | NA | 0.85 | 0.5808 |
| mRNA_4252, mRNA_4251 | 8607 | 3562 | P | mRNA_4252 | 5045 | T | mRNA_4251 | NA | 0.85 | 0.5744 |
| mRNA_4252, mRNA_4251 | 8607 | 7147 | T | mRNA_4251 | 1460 | P | mRNA_4252 | NA | 0.85 | 0.5823 |
| mRNA_4252, mRNA_4251 | 8607 | 5563 | T | mRNA_4251 | 3044 | P | mRNA_4252 | NA | 0.85 | 0.5644 |
| mRNA_4252, mRNA_4250, mRNA_4251 | 8607 | 4073 | T | mRNA_4251 | 4534 | P | mRNA_4252 | NA | 0.85 | 0.4653 |
| mRNA_4249, mRNA_4252, mRNA_4250, mRNA_4251 | 8607 | 3032 | T | mRNA_4251 | 5575 | P | mRNA_4252 | NA | 0.85 | 0.6385 |
| mRNA_4249, mRNA_4252, mRNA_4250, mRNA_4251 | 8607 | 2248 | T | mRNA_4251 | 6359 | P | mRNA_4252 | NA | 0.85 | 0.5574 |
| mRNA_4242, mRNA_4236, mRNA_4235, mRNA_4240, mRNA_4239, mRNA_4241, mRNA_4243, mRNA_4237, mRNA_4238 | 544 | 161 | T | mRNA_4239 | 383 | T | mRNA_4240 | NA | NA | 0.5112 |
| mRNA_4242, mRNA_4236, mRNA_4235, mRNA_4240, mRNA_4239, mRNA_4241, mRNA_4243, mRNA_4237, mRNA_4238 | 544 | 160 | T | mRNA_4239 | 384 | T | mRNA_4240 | NA | NA | 0.4119 |
| mRNA_4233, mRNA_4229, mRNA_4228, mRNA_4234, mRNA_4235, mRNA_4231, mRNA_4232, mRNA_4230 | 3493 | 3247 | T | mRNA_4230 | 246 | T | mRNA_4231 | NA | NA | 0.5538 |
| mRNA_4233, mRNA_4229, mRNA_4228, mRNA_4234, mRNA_4231, mRNA_4232, mRNA_4230 | 3493 | 2903 | T | mRNA_4230 | 590 | T | mRNA_4231 | NA | NA | 0.3329 |
| mRNA_4221, mRNA_4224, mRNA_4220, mRNA_4223, mRNA_4222, mRNA_4218, mRNA_4219, mRNA_4225 | 1994 | 1025 | P | mRNA_4221 | 969 | P | mRNA_4222 | 1 | NA | 0.6586 |
| mRNA_4221, mRNA_4224, mRNA_4220, mRNA_4223, mRNA_4222, mRNA_4218, mRNA_4219, mRNA_4225 | 1994 | 1022 | P | mRNA_4221 | 972 | P | mRNA_4222 | 1 | NA | 0.6147 |
| mRNA_4217, mRNA_4216, mRNA_4221, mRNA_4220, mRNA_4223, mRNA_4222, mRNA_4218, mRNA_4219 | 1352 | 823 | P | mRNA_4220 | 529 | T | mRNA_4221 | 1 | NA | 0.5466 |
| mRNA_4184, mRNA_4183, mRNA_4180, mRNA_4186, mRNA_4182, mRNA_4185, mRNA_4181 | 1843 | 1330 | T | mRNA_4182 | 513 | T | mRNA_4183 | NA | NA | 0.6187 |
| mRNA_4159, mRNA_4155, mRNA_4154, mRNA_4152, mRNA_4151, mRNA_4157, mRNA_4158, mRNA_4160, mRNA_4156, mRNA_4153 | 2198 | 1262 | P | mRNA_4155 | 936 | T | mRNA_4156 | NA | NA | 0.6646 |
| mRNA_4159, mRNA_4155, mRNA_4154, mRNA_4152, mRNA_4151, mRNA_4157, mRNA_4158, mRNA_4160, mRNA_4156, mRNA_4153 | 2198 | 534 | P | mRNA_4155 | 1664 | T | mRNA_4156 | NA | NA | 0.5554 |
| mRNA_4133, mRNA_4131, mRNA_4134, mRNA_4130, mRNA_4132, mRNA_4135, mRNA_4129, mRNA_4128, mRNA_4127 | 1831 | 1231 | P | mRNA_4131 | 600 | P | mRNA_4132 | NA | NA | 0.4322 |
| mRNA_4113, mRNA_4111, mRNA_4109, mRNA_4108, mRNA_4107, mRNA_4110 | 5970 | 429 | P | mRNA_4111 | 5541 | P | mRNA_4113 | NA | NA | 0.5584 |
| mRNA_3753, mRNA_3756, mRNA_3754, mRNA_3758, mRNA_3751, mRNA_3755, mRNA_3757, mRNA_3750 | 1081 | 632 | T | mRNA_3753 | 449 | T | mRNA_3754 | NA | NA | 0.5732 |
| mRNA_3753, mRNA_3756, mRNA_3754, mRNA_3758, mRNA_3751, mRNA_3755, mRNA_3757, mRNA_3750 | 1081 | 584 | T | mRNA_3753 | 497 | T | mRNA_3754 | NA | NA | 0.5468 |
| mRNA_3748, mRNA_3747, mRNA_3749, mRNA_3746, mRNA_3745 | 39112 | 38376 | P | mRNA_3744 | 736 | P | mRNA_3745 | NA | NA | 0.7041 |
| mRNA_3448, mRNA_3454, mRNA_3453, mRNA_3456, mRNA_3449, mRNA_3452, mRNA_3451, mRNA_3455, mRNA_3450 | 1427 | 1075 | T | mRNA_3452 | 352 | T | mRNA_3451 | 1 | NA | 0.6011 |
| mRNA_3484, mRNA_3485, mRNA_3482, mRNA_3480, mRNA_3483, mRNA_3481, mRNA_3487, mRNA_3486 | 946 | 507 | T | mRNA_3484 | 439 | P | mRNA_3483 | NA | NA | 0.5692 |
| mRNA_3484, mRNA_3485, mRNA_3482, mRNA_3480, mRNA_3483, mRNA_3481, mRNA_3487, mRNA_3486 | 946 | 465 | T | mRNA_3484 | 481 | P | mRNA_3483 | NA | NA | 0.6412 |
| mRNA_3490, mRNA_3488, mRNA_3493, mRNA_3489, mRNA_3492, mRNA_3491, mRNA_3487, mRNA_3486 | 1781 | 1286 | P | mRNA_3490 | 495 | T | mRNA_3489 | NA | NA | 0.7375 |
| mRNA_3549, mRNA_3548, mRNA_3543, mRNA_3547, mRNA_3550, mRNA_3544, mRNA_3545, mRNA_3546 | 6322 | 4012 | T | mRNA_3547 | 2310 | P | mRNA_3546 | 1 | NA | 0.3306 |
| mRNA_3675, mRNA_3677, mRNA_3676 | 41828 | 3234 | P | mRNA_3675 | 38594 | P | mRNA_3674 | NA | NA | 0.7174 |
| mRNA_3471, mRNA_3470, mRNA_3472, mRNA_3468, mRNA_3467, mRNA_3469, mRNA_3466 | 1066 | 310 | T | mRNA_3468 | 756 | T | mRNA_3469 | NA | NA | 0.5857 |
| mRNA_3304, mRNA_3301, mRNA_3309, mRNA_3305, mRNA_3300, mRNA_3302, mRNA_3303, mRNA_3306, mRNA_3307, mRNA_3308 | 989 | 242 | T | mRNA_3304 | 747 | P | mRNA_3305 | NA | NA | 0.5 |
| mRNA_3229, mRNA_3226, mRNA_3231, mRNA_3230, mRNA_3227, mRNA_3232, mRNA_3228, mRNA_3225 | 754 | 227 | T | mRNA_3229 | 527 | P | mRNA_3230 | NA | NA | 0.481 |
| | - | 54387 | P | mRNA_4353 | - | - | - | NA | NA | 0.3936 |
| | - | 54386 | P | mRNA_4353 | - | - | - | NA | NA | 0.3443 |
| | - | 20551 | P | mRNA_4353 | - | - | - | NA | NA | 0.5422 |
| | - | - | - | - | 131962 | T | mRNA_4353 | NA | NA | 0.4671 |
| | - | - | - | - | 225865 | T | mRNA_4353 | NA | NA | 0.4583 |
| | - | - | - | - | 342291 | T | mRNA_4353 | NA | NA | 0.5402 |
| | - | - | - | - | 409363 | T | mRNA_4353 | NA | NA | 0.4654 |
| | - | - | - | - | 515669 | T | mRNA_4353 | NA | NA | 0.3209 |
| | - | - | - | - | 516176 | T | mRNA_4353 | NA | NA | 0.5246 |
| | - | - | - | - | 529219 | T | mRNA_4353 | NA | NA | 0.5082 |
| | - | - | - | - | 736948 | T | mRNA_4353 | NA | NA | 0.5037 |
| | - | 738949 | T | mRNA_4353 | - | - | - | NA | NA | 0.5684 |
| | - | 738633 | T | mRNA_4353 | - | - | - | NA | NA | 0.4649 |
| | - | 735633 | T | mRNA_4353 | - | - | - | NA | NA | 0.5648 |
| | - | 718158 | T | mRNA_4353 | - | - | - | NA | NA | 0.4827 |
| | - | 594167 | T | mRNA_4353 | - | - | - | NA | NA | 0.5696 |
| | - | 581977 | T | mRNA_4353 | - | - | - | NA | NA | 0.4461 |
| | - | 569622 | T | mRNA_4353 | - | - | - | NA | NA | 0.4143 |
| | - | 516031 | T | mRNA_4353 | - | - | - | NA | NA | 0.6178 |
| | - | 190752 | T | mRNA_4353 | - | - | - | NA | NA | 0.4816 |
| mRNA_4873, mRNA_4875, mRNA_4872, mRNA_4876, mRNA_4871 | 2821 | 1496 | P | mRNA_4875 | 1325 | T | mRNA_4873 | 1 | NA | 0.5688 |
| mRNA_4873, mRNA_4875, mRNA_4872, mRNA_4876, mRNA_4871 | 2821 | 1172 | P | mRNA_4875 | 1649 | T | mRNA_4873 | 1 | NA | 0.6522 |
| mRNA_4873, mRNA_4875, mRNA_4872, mRNA_4876, mRNA_4871 | 2821 | 1081 | P | mRNA_4875 | 1740 | T | mRNA_4873 | 1 | NA | 0.544 |
| mRNA_4873, mRNA_4875, mRNA_4872, mRNA_4876, mRNA_4871 | 2821 | 1045 | P | mRNA_4875 | 1776 | T | mRNA_4873 | 1 | NA | 0.6634 |
| mRNA_4892, mRNA_4891, mRNA_4889, mRNA_4890 | 1998 | 641 | P | mRNA_4890 | 1357 | P | mRNA_4889 | NA | NA | 0.3126 |
| mRNA_4904, mRNA_4902, mRNA_4903, mRNA_4905, mRNA_4907, mRNA_4906, mRNA_4900, mRNA_4901 | 4950 | 1951 | T | mRNA_4903 | 2999 | T | mRNA_4902 | NA | NA | 0.6342 |
| mRNA_4904, mRNA_4902, mRNA_4903, mRNA_4905, mRNA_4907, mRNA_4906, mRNA_4900, mRNA_4901 | 4950 | 1134 | T | mRNA_4903 | 3816 | T | mRNA_4902 | NA | NA | 0.4009 |
| mRNA_4932, mRNA_4931, mRNA_4937, mRNA_4936, mRNA_4938, mRNA_4934, mRNA_4933, mRNA_4935 | 969 | 451 | T | mRNA_4935 | 518 | P | mRNA_4934 | 1 | NA | 0.5048 |
| mRNA_4949, mRNA_4947, mRNA_4948, mRNA_4950, mRNA_4945, mRNA_4946 | 1506 | 583 | T | mRNA_4949 | 923 | P | mRNA_4948 | NA | NA | 0.5909 |
| mRNA_4953, mRNA_4954, mRNA_4952, mRNA_4956, mRNA_4955 | 1941 | 1049 | T | mRNA_4954 | 892 | P | mRNA_4953 | NA | NA | 0.5049 |
| mRNA_4953, mRNA_4954, mRNA_4952, mRNA_4956, mRNA_4955 | 1941 | 792 | T | mRNA_4954 | 1149 | P | mRNA_4953 | NA | NA | 0.4292 |
| mRNA_4983, mRNA_4989, mRNA_4986, mRNA_4984, mRNA_4985, mRNA_4988, mRNA_4987 | 3551 | 2080 | T | mRNA_4985 | 1471 | T | mRNA_4984 | NA | NA | 0.3278 |
| mRNA_4983, mRNA_4989, mRNA_4986, mRNA_4984, mRNA_4985, mRNA_4988, mRNA_4987 | 3551 | 1830 | T | mRNA_4985 | 1721 | T | mRNA_4984 | NA | NA | 0.459 |
| mRNA_4991, mRNA_4990, mRNA_4989, mRNA_4986, mRNA_4985, mRNA_4995, mRNA_4994, mRNA_4993, mRNA_4992, mRNA_4988, mRNA_4987 | 1015 | 365 | T | mRNA_4990 | 650 | P | mRNA_4989 | NA | NA | 0.5387 |
| mRNA_4991, mRNA_4990, mRNA_4989, mRNA_4986, mRNA_4985, mRNA_4995, mRNA_4994, mRNA_4993, mRNA_4992, mRNA_4988, mRNA_4987 | 1015 | 249 | T | mRNA_4990 | 766 | P | mRNA_4989 | NA | NA | 0.665 |
| mRNA_4991, mRNA_4990, mRNA_4989, mRNA_4986, mRNA_4985, mRNA_4995, mRNA_4994, mRNA_4993, mRNA_4992, mRNA_4988, mRNA_4987 | 1015 | 161 | T | mRNA_4990 | 854 | P | mRNA_4989 | NA | NA | 0.3354 |
| mRNA_5010, mRNA_5000, mRNA_5009, mRNA_5001, mRNA_5005, mRNA_5002, mRNA_5004, mRNA_5011, mRNA_5006, mRNA_5008 | 1239 | 787 | T | mRNA_5006 | 452 | T | mRNA_5005 | NA | NA | 0.4955 |
| mRNA_5021, mRNA_5015, mRNA_5022, mRNA_5016, mRNA_5019, mRNA_5017, mRNA_5018, mRNA_5020 | 2248 | 1603 | T | mRNA_5020 | 645 | T | mRNA_5019 | NA | NA | 0.4925 |
| mRNA_5021, mRNA_5015, mRNA_5022, mRNA_5016, mRNA_5019, mRNA_5017, mRNA_5018, mRNA_5020 | 2248 | 1543 | T | mRNA_5020 | 705 | T | mRNA_5019 | NA | NA | 0.4846 |
| mRNA_5021, mRNA_5015, mRNA_5022, mRNA_5016, mRNA_5019, mRNA_5017, mRNA_5018, mRNA_5020 | 2248 | 1493 | T | mRNA_5020 | 755 | T | mRNA_5019 | NA | NA | 0.6055 |
| mRNA_5037, mRNA_5042, mRNA_5038, mRNA_5036, mRNA_5035, mRNA_5034, mRNA_5040, mRNA_5039, mRNA_5041, mRNA_5043 | 1375 | 733 | P | mRNA_5039 | 642 | P | mRNA_5038 | NA | NA | 0.454 |
| mRNA_5037, mRNA_5042, mRNA_5038, mRNA_5036, mRNA_5035, mRNA_5034, mRNA_5040, mRNA_5039, mRNA_5041, mRNA_5043 | 1375 | 505 | P | mRNA_5039 | 870 | P | mRNA_5038 | NA | NA | 0.5751 |
| mRNA_5037, mRNA_5042, mRNA_5038, mRNA_5036, mRNA_5035, mRNA_5034, mRNA_5040, mRNA_5039, mRNA_5041, mRNA_5043 | 1375 | 478 | P | mRNA_5039 | 897 | P | mRNA_5038 | NA | NA | 0.5024 |

| | | | | | | | | | | |
|---|---|---|---|---|---|---|---|---|---|---|
| mRNA_5037, mRNA_5042, mRNA_5038, mRNA_5036, mRNA_5035, mRNA_5034, mRNA_5040, mRNA_5039, mRNA_5041, mRNA_5043 | 1375 | 462 | P | mRNA_5039 | 913 | P | mRNA_5038 | NA | NA | 0.6197 |
| mRNA_5046, mRNA_5042, mRNA_5047, mRNA_5040, mRNA_5044, mRNA_5051, mRNA_5050, mRNA_5041, mRNA_5049, mRNA_5052, mRNA_5048, | 4264 | 743 | P | mRNA_5046 | 3521 | T | mRNA_5044 | NA | NA | 0.3631 |
| mRNA_5105, mRNA_5103, mRNA_5101, mRNA_5104, mRNA_5102, mRNA_5100, mRNA_5099 | 822 | 331 | T | mRNA_5103 | 491 | P | mRNA_5102 | NA | NA | 0.4071 |
| mRNA_5105, mRNA_5103, mRNA_5101, mRNA_5104, mRNA_5102, mRNA_5100, mRNA_5099 | 822 | 188 | T | mRNA_5103 | 634 | P | mRNA_5102 | NA | NA | 0.7147 |
| mRNA_5127, mRNA_5128 | 16343 | 9066 | T | mRNA_5126 | 7277 | T | mRNA_5127 | NA | 0 | 0.5474 |
| mRNA_5118, mRNA_5113, mRNA_5110, mRNA_5111, mRNA_5117, mRNA_5115, mRNA_5116, mRNA_5114, mRNA_5112 | 2223 | 762 | T | mRNA_5115 | 1461 | P | mRNA_5116 | NA | NA | 0.2872 |
| mRNA_5105, mRNA_5106, mRNA_5103, mRNA_5108, mRNA_5107, mRNA_5104, mRNA_5102 | 1737 | 129 | T | mRNA_5104 | 1608 | P | mRNA_5105 | NA | NA | 0.6852 |
| mRNA_5105, mRNA_5103, mRNA_5101, mRNA_5104, mRNA_5102, mRNA_5100, mRNA_5099 | 822 | 518 | P | mRNA_5102 | 304 | T | mRNA_5103 | NA | NA | 0.6081 |
| mRNA_5046, mRNA_5042, mRNA_5047, mRNA_5040, mRNA_5044, mRNA_5050, mRNA_5041, mRNA_5049, mRNA_5048, mRNA_5043 | 4264 | 1903 | T | mRNA_5044 | 2361 | P | mRNA_5046 | NA | NA | 0.7586 |
| mRNA_5046, mRNA_5042, mRNA_5047, mRNA_5040, mRNA_5044, mRNA_5050, mRNA_5041, mRNA_5049, mRNA_5048, mRNA_5043 | 4264 | 1564 | T | mRNA_5044 | 2700 | P | mRNA_5046 | NA | NA | 0.5624 |
| mRNA_5046, mRNA_5042, mRNA_5047, mRNA_5040, mRNA_5044, mRNA_5050, mRNA_5041, mRNA_5049, mRNA_5048, mRNA_5043 | 4264 | 1543 | T | mRNA_5044 | 2721 | P | mRNA_5046 | NA | NA | 0.4511 |
| mRNA_5037, mRNA_5042, mRNA_5038, mRNA_5036, mRNA_5035, mRNA_5034, mRNA_5040, mRNA_5039, mRNA_5041, mRNA_5043 | 1375 | 944 | P | mRNA_5038 | 431 | P | mRNA_5039 | NA | NA | 0.6371 |
| mRNA_5037, mRNA_5042, mRNA_5038, mRNA_5036, mRNA_5035, mRNA_5034, mRNA_5040, mRNA_5039, mRNA_5041, mRNA_5043 | 1375 | 671 | P | mRNA_5038 | 704 | P | mRNA_5039 | NA | NA | 0.5898 |
| mRNA_5037, mRNA_5042, mRNA_5038, mRNA_5036, mRNA_5035, mRNA_5034, mRNA_5040, mRNA_5039, mRNA_5041, mRNA_5043 | 1375 | 668 | P | mRNA_5038 | 707 | P | mRNA_5039 | NA | NA | 0.5536 |
| mRNA_5037, mRNA_5042, mRNA_5038, mRNA_5036, mRNA_5035, mRNA_5034, mRNA_5040, mRNA_5039, mRNA_5041, mRNA_5043 | 1375 | 657 | P | mRNA_5038 | 718 | P | mRNA_5039 | NA | NA | 0.6602 |
| mRNA_5021, mRNA_5015, mRNA_5022, mRNA_5016, mRNA_5019, mRNA_5017, mRNA_5018, mRNA_5020 | 2248 | 778 | T | mRNA_5019 | 1470 | T | mRNA_5020 | NA | NA | 0.5221 |
| mRNA_4983, mRNA_4990, mRNA_4989, mRNA_4986, mRNA_4984, mRNA_4985, mRNA_4988, mRNA_4987 | 1105 | 591 | P | mRNA_4985 | 514 | P | mRNA_4986 | NA | NA | 0.5378 |
| mRNA_4983, mRNA_4990, mRNA_4989, mRNA_4986, mRNA_4984, mRNA_4985, mRNA_4988, mRNA_4987 | 1105 | 590 | P | mRNA_4985 | 515 | P | mRNA_4986 | NA | NA | 0.6665 |
| mRNA_4983, mRNA_4990, mRNA_4989, mRNA_4986, mRNA_4984, mRNA_4985, mRNA_4988, mRNA_4987 | 1105 | 458 | P | mRNA_4985 | 647 | P | mRNA_4986 | NA | NA | 0.3641 |
| mRNA_4983, mRNA_4989, mRNA_4986, mRNA_4984, mRNA_4985, mRNA_4988, mRNA_4987 | 3551 | 2823 | T | mRNA_4984 | 728 | T | mRNA_4985 | NA | NA | 0.5427 |
| mRNA_4983, mRNA_4989, mRNA_4986, mRNA_4984, mRNA_4985, mRNA_4988, mRNA_4987 | 3551 | 2806 | T | mRNA_4984 | 745 | T | mRNA_4985 | NA | NA | 0.6093 |
| mRNA_4983, mRNA_4989, mRNA_4986, mRNA_4984, mRNA_4985, mRNA_4988, mRNA_4987 | 3551 | 1620 | T | mRNA_4984 | 1931 | T | mRNA_4985 | NA | NA | 0.6022 |
| mRNA_4983, mRNA_4989, mRNA_4986, mRNA_4984, mRNA_4985, mRNA_4988, mRNA_4987 | 3551 | 1616 | T | mRNA_4984 | 1935 | T | mRNA_4985 | NA | NA | 0.5178 |
| mRNA_4983, mRNA_4984, mRNA_4980, mRNA_4978, mRNA_4982, mRNA_4985, mRNA_4981, mRNA_4979 | 2842 | 749 | P | mRNA_4982 | 2093 | P | mRNA_4983 | NA | 0.16 | 0.3001 |
| mRNA_4949, mRNA_4947, mRNA_4948, mRNA_4950, mRNA_4951 | 1531 | 794 | P | mRNA_4949 | 737 | P | mRNA_4950 | 1 | NA | 0.7222 |
| mRNA_4949, mRNA_4947, mRNA_4948, mRNA_4950, mRNA_4951 | 1531 | 793 | P | mRNA_4949 | 738 | P | mRNA_4950 | 1 | NA | 0.502 |
| mRNA_4949, mRNA_4947, mRNA_4948, mRNA_4950, mRNA_4945, mRNA_4946 | 1506 | 810 | P | mRNA_4948 | 696 | T | mRNA_4949 | NA | NA | 0.665 |
| mRNA_4949, mRNA_4947, mRNA_4948, mRNA_4950, mRNA_4945, mRNA_4946 | 1506 | 653 | P | mRNA_4948 | 853 | T | mRNA_4949 | NA | NA | 0.5603 |
| mRNA_4944, mRNA_4947, mRNA_4948, mRNA_4942, mRNA_4943, mRNA_4945, mRNA_4946 | 1514 | 591 | P | mRNA_4945 | 923 | P | mRNA_4946 | 1 | NA | 0.6361 |
| mRNA_4932, mRNA_4929, mRNA_4931, mRNA_4930, mRNA_4928, mRNA_4927, mRNA_4926, mRNA_4933 | 1398 | 929 | P | mRNA_4929 | 469 | T | mRNA_4930 | NA | NA | 0.3611 |
| mRNA_4904, mRNA_4908, mRNA_4902, mRNA_4903, mRNA_4905, mRNA_4907, mRNA_4906 | 1631 | 1451 | P | mRNA_4905 | 180 | T | mRNA_4906 | NA | NA | 0.7208 |
| mRNA_4904, mRNA_4902, mRNA_4903, mRNA_4905, mRNA_4907, mRNA_4906, mRNA_4900, mRNA_4901 | 4950 | 2980 | T | mRNA_4902 | 1970 | T | mRNA_4903 | NA | NA | 0.283 |
| mRNA_4892, mRNA_4891, mRNA_4889, mRNA_4890, mRNA_4888 | 1998 | 942 | P | mRNA_4889 | 1056 | P | mRNA_4890 | NA | NA | 0.5974 |
| mRNA_4889, mRNA_4887, mRNA_4885, mRNA_4888, mRNA_4886, mRNA_4884 | 1652 | 1467 | T | mRNA_4887 | 185 | T | mRNA_4888 | 1 | NA | 0.7439 |
| mRNA_4889, mRNA_4887, mRNA_4885, mRNA_4888, mRNA_4886, mRNA_4884 | 1652 | 704 | T | mRNA_4887 | 948 | T | mRNA_4888 | 1 | NA | 0.65 |
| mRNA_4873, mRNA_4875, mRNA_4872, mRNA_4876, mRNA_4871 | 2821 | 1699 | T | mRNA_4873 | 1122 | P | mRNA_4875 | 1 | NA | 0.4087 |
| mRNA_4873, mRNA_4875, mRNA_4872, mRNA_4876, mRNA_4871 | 2821 | 1584 | T | mRNA_4873 | 1237 | P | mRNA_4875 | 1 | NA | 0.3698 |
| | 99853 | 31396 | T | mRNA_7412 | 68457 | P | mRNA_7354 | NA | NA | 0.5362 |
| | 99853 | 25530 | T | mRNA_7412 | 74323 | P | mRNA_7354 | NA | NA | 0.7313 |
| mRNA_7412 | - | - | - | - | 672 | P | mRNA_7412 | NA | NA | 0.6462 |
| | - | - | - | - | 9856 | P | mRNA_7412 | NA | NA | 0.5362 |
| | 99853 | 74052 | P | mRNA_7354 | 25801 | T | mRNA_7412 | NA | NA | 0.5882 |
| | 99853 | 67470 | P | mRNA_7354 | 32383 | T | mRNA_7412 | NA | NA | 0.6633 |
| | 99853 | 45988 | P | mRNA_7354 | 53865 | T | mRNA_7412 | NA | NA | 0.5597 |
| | 99853 | 17067 | P | mRNA_7354 | 82786 | T | mRNA_7412 | NA | NA | 0.5171 |
| | - | - | - | - | 9998 | P | mRNA_7350 | NA | NA | 0.6309 |
| | - | - | - | - | 92254 | P | mRNA_7350 | NA | NA | 0.2837 |
| | - | - | - | - | 99811 | P | mRNA_7350 | NA | NA | 0.6018 |
| | - | - | - | - | 123692 | T | mRNA_7673 | NA | NA | 0.5602 |
| | - | - | - | - | 22725 | T | mRNA_7672 | NA | NA | 0.5019 |
| | - | - | - | - | 115865 | T | mRNA_7672 | NA | NA | 0.416 |
| mRNA_7906, mRNA_7905, mRNA_7908, mRNA_7907 | - | 289 | T | mRNA_7905 | - | - | - | 1 | NA | 0.4659 |
| mRNA_7919, mRNA_7925, mRNA_7924, mRNA_7923, mRNA_7920, mRNA_7922, mRNA_7918, mRNA_7917 | 1178 | 728 | P | mRNA_7920 | 450 | P | mRNA_7922 | NA | 45.83 | 0.6093 |
| | - | 202117 | T | mRNA_8651 | - | - | - | NA | NA | 0.5544 |
| | - | 169465 | T | mRNA_8651 | - | - | - | NA | NA | 0.5723 |
| | - | 133423 | T | mRNA_8651 | - | - | - | NA | NA | 0.5122 |
| | - | 129383 | T | mRNA_8651 | - | - | - | NA | NA | 0.5851 |
| | - | 51340 | T | mRNA_8651 | - | - | - | NA | NA | 0.5304 |
| mRNA_8651 | - | - | - | - | 1360 | T | mRNA_8651 | NA | NA | 0.5725 |
| | - | - | - | - | 86778 | T | mRNA_8651 | NA | NA | 0.591 |
| | - | - | - | - | 190383 | T | mRNA_8651 | NA | NA | 0.6384 |
| mRNA_8881, mRNA_8880 | - | 4769 | T | mRNA_8881 | - | - | - | NA | NA | 0.4558 |
| mRNA_9084, mRNA_9083, mRNA_9081, mRNA_9080, mRNA_9082 | 7971 | 5111 | P | mRNA_9084 | 2860 | P | mRNA_9083 | NA | 0 | 0.424 |
| mRNA_9084, mRNA_9083, mRNA_9082 | 7971 | 898 | P | mRNA_9084 | 7073 | P | mRNA_9083 | NA | 0 | 0.5669 |
| mRNA_9084, mRNA_9083, mRNA_9081, mRNA_9082 | 7971 | 5297 | P | mRNA_9083 | 2674 | P | mRNA_9084 | NA | 0 | 0.5271 |
| mRNA_9084, mRNA_9083, mRNA_9081, mRNA_9080, mRNA_9082 | 7971 | 2750 | P | mRNA_9083 | 5221 | P | mRNA_9084 | NA | 0 | 0.4786 |
| mRNA_9220 | 16740 | 7416 | T | mRNA_9220 | 9324 | T | mRNA_9219 | NA | NA | 0.5335 |
| mRNA_9220 | 16740 | 6953 | T | mRNA_9220 | 9787 | T | mRNA_9219 | NA | NA | 0.5508 |
| mRNA_9220 | 16740 | 6749 | T | mRNA_9220 | 9991 | T | mRNA_9219 | NA | NA | 0.3212 |
| mRNA_9220 | 16740 | 6723 | T | mRNA_9220 | 10017 | T | mRNA_9219 | NA | NA | 0.5495 |
| mRNA_9220 | 16740 | 6722 | T | mRNA_9220 | 10018 | T | mRNA_9219 | NA | NA | 0.5192 |
| mRNA_9220 | 16740 | 789 | T | mRNA_9220 | 15951 | T | mRNA_9219 | NA | NA | 0.4474 |
| mRNA_9220 | 16740 | 10059 | T | mRNA_9219 | 6681 | T | mRNA_9220 | NA | NA | 0.5467 |
| mRNA_9220 | 16740 | 10044 | T | mRNA_9219 | 6696 | T | mRNA_9220 | NA | NA | 0.5016 |
| mRNA_9220 | 16740 | 10033 | T | mRNA_9219 | 6707 | T | mRNA_9220 | NA | NA | 0.5784 |
| mRNA_9220 | 16740 | 9829 | T | mRNA_9219 | 6911 | T | mRNA_9220 | NA | NA | 0.4818 |
| mRNA_9220 | 16740 | 9820 | T | mRNA_9219 | 6920 | T | mRNA_9220 | NA | NA | 0.606 |
| mRNA_9338 | - | 1018 | P | mRNA_9338 | - | - | - | NA | NA | 0.5599 |
